# Comparative genomics of *Cryptococcus* and *Kwoniella* reveals pathogenesis evolution and contrasting karyotype dynamics via intercentromeric recombination or chromosome fusion

**DOI:** 10.1101/2023.12.27.573464

**Authors:** Marco A. Coelho, Márcia David-Palma, Terrance Shea, Katharine Bowers, Sage McGinley-Smith, Arman W. Mohammad, Andreas Gnirke, Andrey M. Yurkov, Minou Nowrousian, Sheng Sun, Christina A. Cuomo, Joseph Heitman

## Abstract

A large-scale comparative genomic analysis was conducted for the global human fungal pathogens within the *Cryptococcus* genus, compared to non-pathogenic *Cryptococcus* species, and related species from the sister genus *Kwoniella*. Chromosome-level genome assemblies were generated for multiple species of both genera, resulting in a dataset encompassing virtually all of their known diversity. Although *Cryptococcus* and *Kwoniella* have comparable genome sizes (about 19.2 and 22.9 Mb) and similar gene content, hinting at pre-adaptive pathogenic potential, our analysis found evidence in pathogenic *Cryptococcus* species of specific examples of gene gain (via horizontal gene transfer) and gene loss, which might represent evolutionary signatures of pathogenic development. Genome analysis also revealed a significant variation in chromosome number and structure between the two genera. By combining synteny analysis and experimental centromere validation, we found that most *Cryptococcus* species have 14 chromosomes, whereas most *Kwoniella* species have fewer (11, 8, 5 or even as few as 3). Reduced chromosome number in *Kwoniella* is associated with formation of giant chromosomes (up to 18 Mb) through repeated chromosome fusion events, each marked by a pericentric inversion and centromere loss. While similar chromosome inversion-fusion patterns were observed in all *Kwoniella* species with fewer than 14 chromosomes, no such pattern was detected in *Cryptococcus*. Instead, *Cryptococcus* species with less than 14 chromosomes, underwent chromosome reductions primarily through rearrangements associated with the loss of repeat-rich centromeres. Additionally, *Cryptococcus* genomes exhibited frequent interchromosomal translocations, including intercentromeric recombination facilitated by transposons shared between centromeres. Taken together, our findings advance our understanding of genomic changes possibly associated with pathogenicity in *Cryptococcus* and provide a foundation to elucidate mechanisms of centromere loss and chromosome fusion driving distinct karyotypes in closely related fungal species, including prominent global human pathogens.

## Introduction

Human fungal pathogens are estimated to be responsible for over 1.5 million deaths worldwide annually [1]. Growing concern regarding invasive fungal diseases led the World Health Organization (WHO) to publish its first list of priority fungal pathogens, with *Cryptococcus neoformans* ranked first in the critical group [2]. Even though direct selection of traits favoring human virulence or infection is not expected because humans are not the natural hosts for any pathogenic *Cryptococcus* species, interactions of these yeasts with other eukaryotes in their natural environments could select traits that enhance virulence in mammals [3-6]. Exploring the evolutionary trajectories of pathogenic and non-pathogenic *Cryptococcus* species and closely related genera can provide insights into the broader mechanisms that govern fungal evolution and shed light on why some species, but not others, have evolved traits that cause pathogenesis in humans [7, 8].

The genus *Cryptococcus* encompasses diverse species, including both pathogenic and closely related non-pathogenic saprobic species [9-12]. Within the pathogenic clade, there are seven recognized species that can be pathogenic for both immunocompromised and immunocompetent individuals [13-15]: *Cryptococcus neoformans*, *Cryptococcus deneoformans*, and five species within the *Cryptococcus gattii* species complex [9]. The *C. gattii* complex also includes another recently discovered lineage described as *C. gattii* VGV [16], which has not yet been linked to human infection. The non-pathogenic species include *Cryptococcus wingfieldii*, *Cryptococcus amylolentus*, *Cryptococcus floricola*, *Cryptococcus depauperatus* and *Cryptococcus luteus* [11, 12, 17, 18]. Phylogenetically, the genus *Kwoniella* is considered the closest relative to *Cryptococcus* [17-19], and all known *Kwoniella* species are saprophytic in nature [10, 20-22].

Although *Cryptococcus* and *Kwoniella* genome assemblies have improved in quality and completeness in recent years [11, 12, 23-25], there has been no comprehensive comparative genomic study so far that leverages complete genome assemblies. By comparing chromosome-level genome assemblies of multiple strains and species, we can now identify the extent of collinearity between the chromosomes of different species and infer structural variation, as well as identify specific regions where variation occurs, with a high degree of accuracy. Chromosomal rearrangements, including inversions, translocations, fusions, and fissions, underlie the extensive karyotypic variability seen across eukaryotes [26-28]. These sources of structural variation aid organisms in adapting to diverse environments [29, 30], influence changes in mating systems [23, 31-33], drive the evolution of pathogenic traits [34-36], and play roles in speciation [37-41].

Chromosome number reduction is common in eukaryotes and often results from chromosome fusion events. This is exemplified in several muntjac deer species, where karyotype changes occurred through chromosome fusion during speciation [41], in the formation of the extant human chromosome 2 [42, 43], in chromosome number reduction in the plant *Arabidopsis thaliana* [44, 45], and in the nematode *Diploscapter pachys* that achieved a single-chromosome karyotype through fusion of six ancestral chromosomes [46]. In fungi, telomere-to-telomere fusions altering chromosome numbers are also observed, such as in Ascomycota yeast species [47], and in *Fusarium graminearum*, where chromosome fusion has led to a reduced karyotype compared to related species, with the sites of fusion corresponding to former subtelomeric regions retaining high genetic diversity and heterochromatin marks [48-50]. Similarly, chromosome fusion is suggested to underlie karyotype reduction in Basidiomycota *Malassezia* species [51, 52]

Here, we explored the genomic characteristics of *Cryptococcus* and *Kwoniella* species, focusing on karyotype and gene variation. We generated high-quality chromosome-level genome assemblies for 22 species, capturing most of the known diversity of the two genera. Analysis of structural variation across species revealed contrasting mechanisms of karyotype evolution. In *Kwoniella*, karyotypic variation was primarily driven by chromosome fusion events involving pericentric inversions and centromere loss, leading to chromosome numbers varying from 14 to as few as 3, without notable loss of genomic information. Interestingly, in species with only three chromosomes, one has evolved into a “giant” chromosome (∼16 to 18 Mb) formed by successive fusion events. In stark contrast, *Cryptococcus* species largely retained the ancestral karyotypic arrangement of 14 chromosomes, apart from two species where chromosome number reduction entailed substantial interchromosomal rearrangements and centromere inactivation through loss of repeat-rich sequences. We also examined gene content variation, finding that genes related to canonical pathogenesis phenotypes are highly conserved across all species, although some notable pathogen-specific gene signatures were identified. This study highlights key genetic characteristics of both genera and establishes a foundation for understanding pathogenesis evolution, centromere loss, and the role of chromosome fusions in the evolution and adaptation of these fungal species.

## Results

### Chromosome-level assemblies, genomic features, and phylogeny of *Cryptococcus* and *Kwoniella* species

We generated 22 chromosome-level genome assemblies for 7 *Cryptococcus* and 15 *Kwoniella* species by combining long- (Oxford Nanopore or PacBio) and short-read (Illumina) sequencing (**Fig 1A** and **S1 Fig**). This dataset includes two new *Cryptococcus* (isolates OR849 and OR918 [53]) and four new *Kwoniella* species (isolates B9012, CBS6097, CBS9459, and DSM27149) that will be formally described elsewhere (**Fig 1**). Five of the newly obtained *Cryptococcus* assemblies belong to pathogenic species known to cause disease in humans and animals: four of these are improved assemblies of species within the *Cryptococcus gattii* complex and the other one is an updated assembly of *C. deneoformans* reference strain JEC21. All of these assemblies have higher contiguity, particularly at centromeric and subtelomeric regions. We also incorporated the genome assemblies of seven other *Cryptococcus* species generated in previous studies [11, 12, 16, 24, 25, 39, 54] for comparative genomic analysis. This resulted in a comprehensive dataset of 14 *Cryptococcus* and 15 *Kwoniella* genomes, covering most of the currently known diversity within the two groups (**Fig 1A**, **S1 Fig** and **S1 Appendix**).

**Fig 1.**
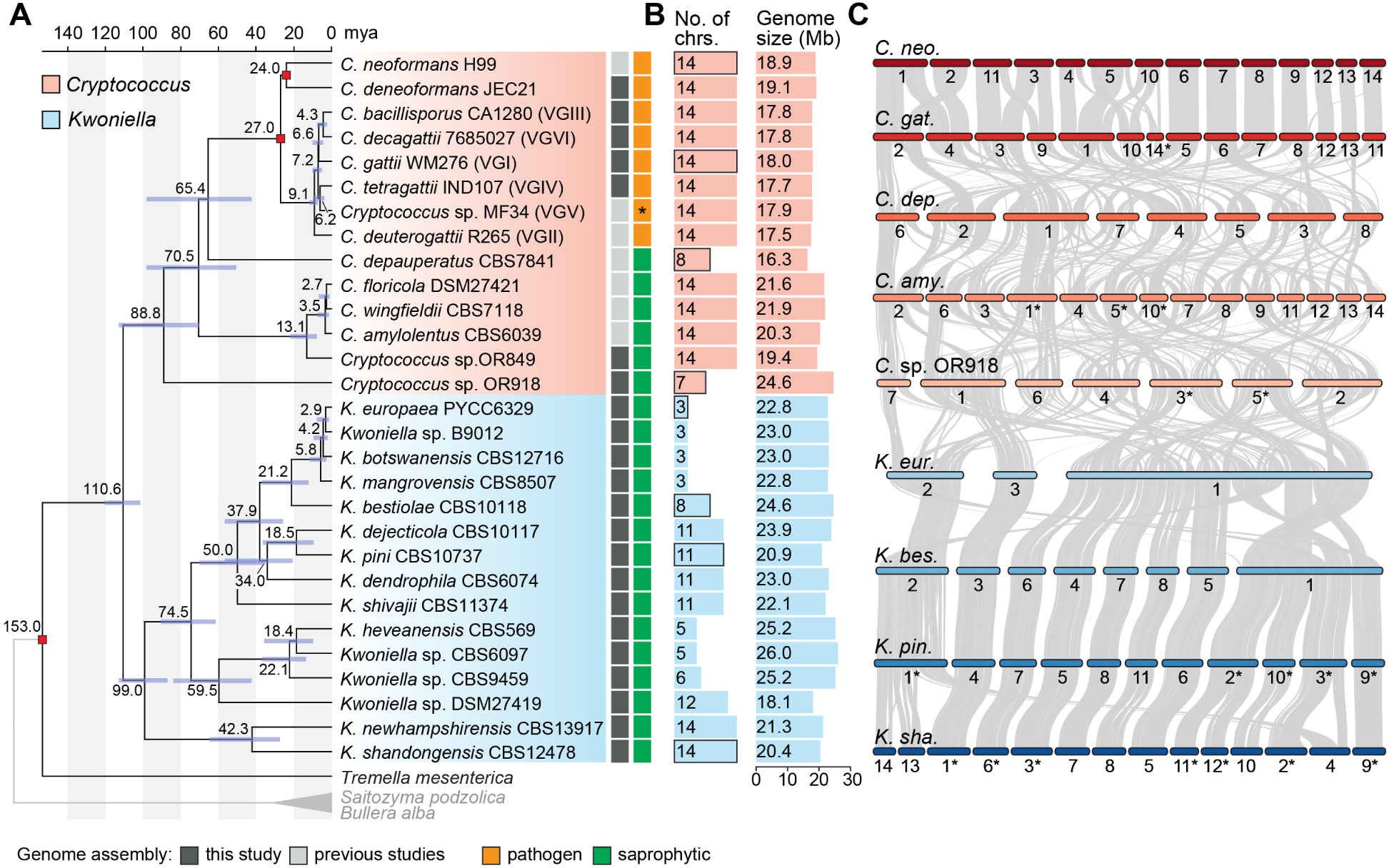
Molecular time tree for *Cryptococcus* and *Kwoniella* and genome composition. **(A)** Time tree including all *Cryptococcus* and *Kwoniella* strains analyzed in this study. Different genera are depicted by different-colored boxes as shown in the key. Divergence time estimation was conducted with the RelTime algorithm employing a phylogeny inferred by maximum likelihood analysis using a concatenation-based approach on a data matrix composed of protein alignments of 3,430 single-copy genes shared across all species and three outgroups. All branches are 100% supported (SH-aLRT and UFboot tests; see S1 Fig). For dating estimation, *T. mesenterica* was included as ingroup and absolute divergence time estimates were calibrated using the following constraints: separation of *T. mesenterica* from other species [153.0 million years ago (mya)], the origin of the pathogenic *Cryptococcus* species (27.0 mya) and the *C. neoformans* and *C. deneoformans* split (24 mya). Blue boxes around each internode correspond to 95% divergence time confidence intervals for each branch of the phylogeny. Complete genome sequences obtained in this study are marked in dark gray. An asterisk indicates that VGV has not as yet been linked to human infection. **(B)** Number of chromosomes and genome size (Mb, megabase pair). **(C)** Pairwise synteny relationships between representative *Cryptococcus* and *Kwoniella* species with different number of chromosomes highlighting markedly distinct routes of karyotypic evolution (many interchromosomal rearrangements in *Cryptococcus* vs. chromosome fusion events in *Kwoniella*). Links represent the boundaries of syntenic gene blocks identified by MCScanX with pairwise homologous relationships determined by SynChro. Chromosomes were reordered and/or inverted (marked with asterisks) relative to their original assembly orientations to maximize collinearity.

Gene set evaluation, following gene prediction and annotation (see Material and Methods and **S1 Appendix**), revealed a high level of completeness, with an average of 97.8% presence of Benchmarking Universal Single-Copy Orthologs (BUSCO) genes from the fungal tremellomycetes_odb10 database, ranging from 93.9% to 99.7% (**S1C Fig**). This underscores the high-quality of the assembled genomes and their gene sets. Further analysis showed that, on average, *Cryptococcus* genomes are smaller (16.3 to 24.6 Mb, average 19.2 Mb) compared to *Kwoniella* genomes (18.1 to 26.0 Mb, average 22.9 Mb) (**Fig 1B** and **S1H Fig;** *P* = 0.0004, Mann-Whitney U Test). The larger sizes of *Kwoniella* genomes can be attributed to both a higher average number of predicted genes (*P* = 0.004, Mann-Whitney U Test) and longer introns (*P* < 0.0001, Mann-Whitney U Test) (**S1I-K Figs**). However, while larger genome sizes among *Cryptococcus* species is strongly correlated with a higher number of genes (*P* < 0.0001, *R*^2^ = 0.824), this association is less pronounced in *Kwoniella* (*P* = 0.0446, *R*^2^ = 0.275; **S1L Fig**), where longer introns appear to be a significant factor in genome size (*P* < 0.0001, *R*^2^ = 0.793; **S1M Fig**). Additionally, no significant correlation was found between the number of introns per coding sequence and genome size changes within or between the two groups (**S1K** and **S1N Figs**).

Comparative genomic analysis also revealed striking differences in chromosome numbers between the two groups. With the exception of two species, all other *Cryptococcus* species analyzed have 14 chromosomes, in contrast to *Kwoniella* where chromosome numbers range widely. Some species, like *Kwoniella shandongensis* and *Kwoniella newhampshirensis*, have 14 chromosomes, whereas others, such as *Kwoniella mangrovensis*, have as few as 3 chromosomes (**Fig 1B**). Owing to these variations in *Kwoniella*, we carried out pulsed-field gel electrophoresis and Hi-C mapping as additional validation measures to ensure accuracy of a subset of the assemblies, particularly for those species with fewer chromosomes. The combined results of both methods corroborated our assemblies (**S2 Fig**).

Prompted by these findings we conducted an in-depth investigation of the karyotypic evolution within and across these two groups. To provide a phylogenetic framework for subsequent evolutionary analysis, we first established phylogenetic relationships by identifying 3,430 single-copy genes shared across all *Cryptococcus* and *Kwoniella* species, and three outgroups. Phylogenetic reconstructions based on a concatenation-based approach differentiated *Cryptococcus* and *Kwoniella* species into distinct clades and validated the status of some of the new isolates as distinct species, as evidenced by their clear phylogenetic separation and divergence (**Fig 1A** and **S1 Fig**). Estimation of the divergence times using the RelTime method and three calibration points [55, 56] suggests that *Cryptococcus* and *Kwoniella* diverged from a shared last common ancestor approximately 110 million years ago (mya), and the initial split within *Cryptococcus* and *Kwoniella* occurred around 90 and 100 mya, respectively (**Fig 1A**). While the estimated divergence times are in general consistent with other studies [57-59], we note that previous estimates for the last common ancestor of disease-causing Cryptococci pointed to an earlier divergence time ranging from 40 to 100 mya [60, 61]. Nevertheless, even if our analysis underestimates the divergence times, it still suggests that the initial divergence within the two groups occurred at roughly the same time.

### Distinct modes of chromosome evolution in *Cryptococcus* and *Kwoniella*

To understand the pronounced differences in chromosome number between *Cryptococcus* and *Kwoniella*, a detailed analysis of chromosomal rearrangements was conducted based on whole-genome alignments from species with varying chromosome numbers in both groups. As shown in **Figs 1A** and **1C**, although the crown node times of *Cryptococcus* and *Kwoniella* are relatively similar, the two groups have experienced distinct types of chromosomal rearrangements throughout evolution. Within *Cryptococcus*, interchromosomal rearrangements predominate, whereas chromosome fusions are the dominant type of rearrangement within *Kwoniella* and seem to account for the extensive variation in chromosome number across species (**Fig 1C**). Previous studies have linked large-scale interchromosomal rearrangements in *Cryptococcus* to recombination within gene-devoid, transposable element-rich, centromeres [23, 39]. Combining this genomic hallmark of *Cryptococcus* centromeres [11, 12, 24, 62-64] with synteny analysis, we assigned *in silico* each of the 14 centromeres of *C. neoformans* to a predicted centromeric region in *K. shandongensis*, a *Kwoniella* species with 14 fully assembled chromosomes (**S3 Fig**). This karyotypic similarity between *C. neoformans* and *K. shandongensis* supports the hypothesis that both lineages descended from a common ancestor with a 14-chromosome karyotype.

### Karyotype reduction in *Kwoniella* occurred over multiple speciation events through recurrent chromosome-chromosome fusions

To trace the sequence of chromosomal rearrangements that unfolded throughout *Kwoniella* evolution, synteny blocks were reconstructed with SynChro [65], employing *K. shandongensis* as reference. As detailed in **Fig 2**, karyotype reduction within *Kwoniella* occurred both progressively and independently through multiple speciation events, primarily through chromosome fusions. We identified fusion events that have emerged more ancestrally, such as the fusion between chromosomes corresponding to *K. shandongensis* chrs. 11 and 12, occurring between 99 and 74.5 mya (event A in **Fig 2** and **S4A Fig**), which is consistently seen as an individual chromosome in 8 of 13 species with fewer than 14 chromosomes. Another ancestral event combined chrs. 14, 13, and 2 (**S4B Fig)**. The outcome of this fusion persists as a single chromosome (chr. 1) in *Kwoniella* sp. DSM27419 (**S4B Fig**), but in other species this chromosome became more rearranged due to intrachromosomal rearrangements (as observed in *K. heveanensis* and closely related species) or underwent a translocation with another chromosome prior to diversification (event B in **Fig 2**). These fusion events led to the emergence of an 11-chromosome state in one of the lineages that branched off from the *Kwoniella* common ancestor and persisted in four extant *Kwoniella* species (*K. dejecticola*, *K. pini*, *K. dendrophila*, and *K. shivajii*). The other lineage, comprising *K. shandongensis* and *K. newhampshirensis*, retained the 14-chromosome karyotype.

**Fig 2.**
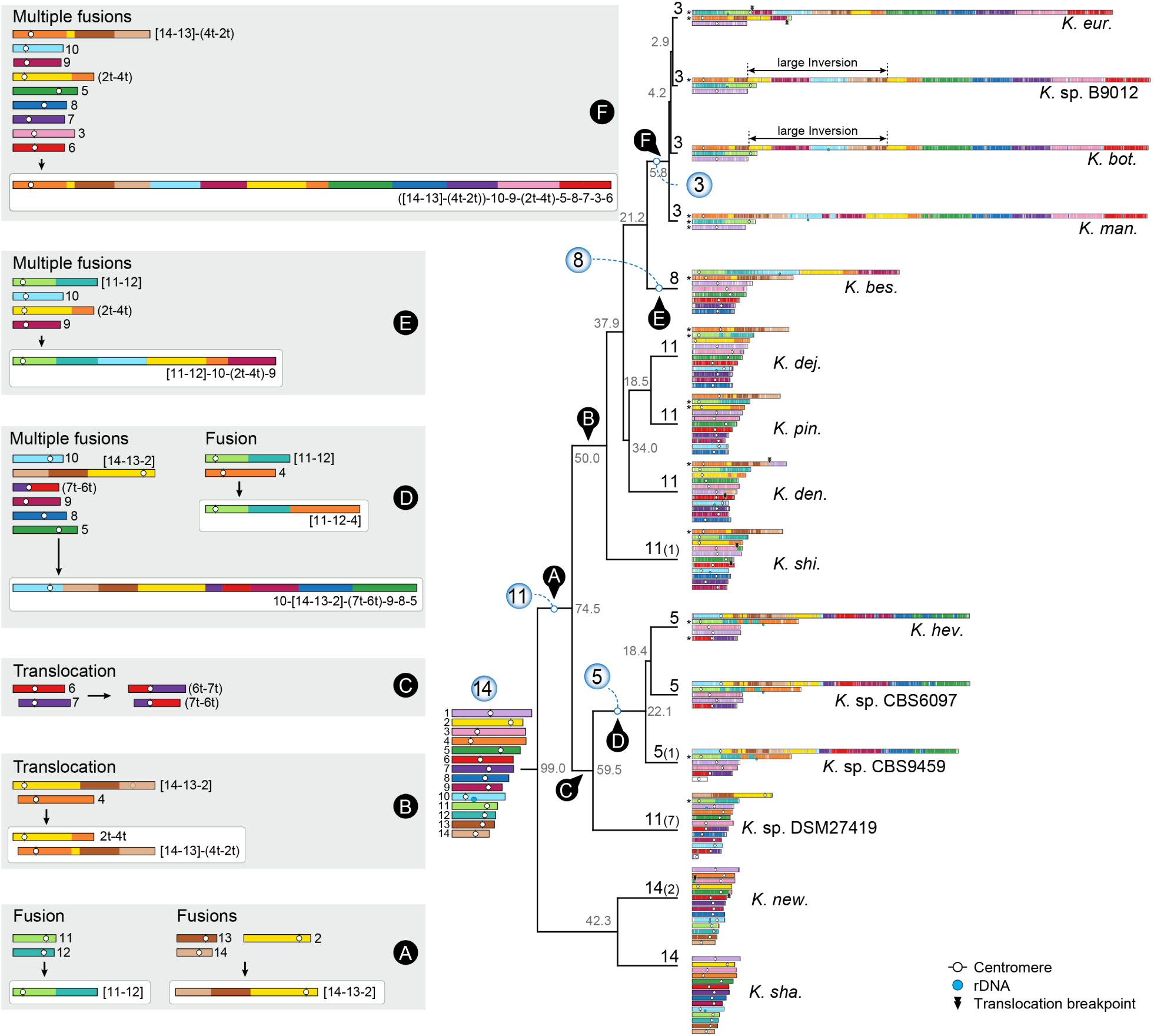
Karyotype reduction in *Kwoniella* occurred independently through recurrent chromosome fusions across multiple speciation events. Major chromosomal rearrangement events, labeled from A to F, are illustrated along the evolution of *Kwoniella*. The karyotype of *K. shandongensis* (with 14 chromosomes) served as the reference for reconstructing synteny blocks in pairwise comparisons. The number of chromosomes in each species is depicted at the tips of the tree, with numbers in parentheses indicating additional small chromosomes lacking clear syntenic relationships with any other chromosome; these may constitute supernumerary, mini-chromosomes, or recently formed chromosomes. Only chromosomes above 100 kb are illustrated, with asterisks marking those inverted from their original assembly orientations. Numbers in gray around each internode represent median divergence time estimates in millions of years, obtained from Fig 1. Chromosomal fusions and translocations are labeled, respectively, within brackets or parentheses, based on their original IDs (e.g., “[11-12]” indicates a chromosome resulting from fusion of ancestral chrs. 11 and 12; “(7t-6t)” represents one of the chromosomes resulting from a reciprocal translocation between ancestral chrs. 6 and 7).

More recent fusion events, between 59.5 to 5.8 mya, led to unique chromosomal arrangements in different lineages. In *K. heveanensis*, *Kwoniella* sp. CBS6097, and *Kwoniella* sp. CBS9459, our analysis suggests their common ancestor had a 5-chromosome karyotype. Specifically, chr. 2 emerged through fusion of the pre-fused chr. 11-12 with a chromosome corresponding to chr. 4 of *K. shandongensis* (event D in **Fig 2**), followed by species-specific intrachromosomal rearrangements, mainly inversions (**S4A Fig**). Likewise, chr. 1 seems to be the product of a past event where six chromosomes underwent fusion, generating a ∼13-Mb chromosome (event D in **Fig 2**, and **S5 Fig**). Parallel events involving multiple chromosome fusions also took place in *K. bestiolae*, where a different set of four chromosomes fused together (event E in **Fig 2**, and **S6 Fig**). Yet, the most striking event, entailing the fusion of 9 chromosomes, is inferred to have occurred between 21.2 to 5.8 mya, in the shared ancestor of *K. europaea*, *Kwoniella* sp. B9012, *K. botswanensis*, and *K. mangrovensis*, leading to a 3-chromosome karyotype with an exceptionally large ∼18.2 Mb chromosome (event F in **Fig2**). Post-branching from *K. mangrovensis*, this “giant” chromosome, which is 7 to 8 times larger than the other two, underwent structural changes, including a large inversion and a reciprocal translocation with chr. 2 in *K. europaea*, along with smaller, independent inversions in each species (**Fig 3A**). The data at hand does not allow, however, determining whether the 8 fusion events underlying the formation of the “giant” chromosome occurred in a single simultaneous step or through multiple consecutive events. This naturally formed “giant” chromosome is unusual and novel, particularly given its size, and its formation has clearly been tolerated despite some evidence suggesting that larger chromosomes may face additional challenges during mitotic segregation and replication [66].

**Fig 3.**
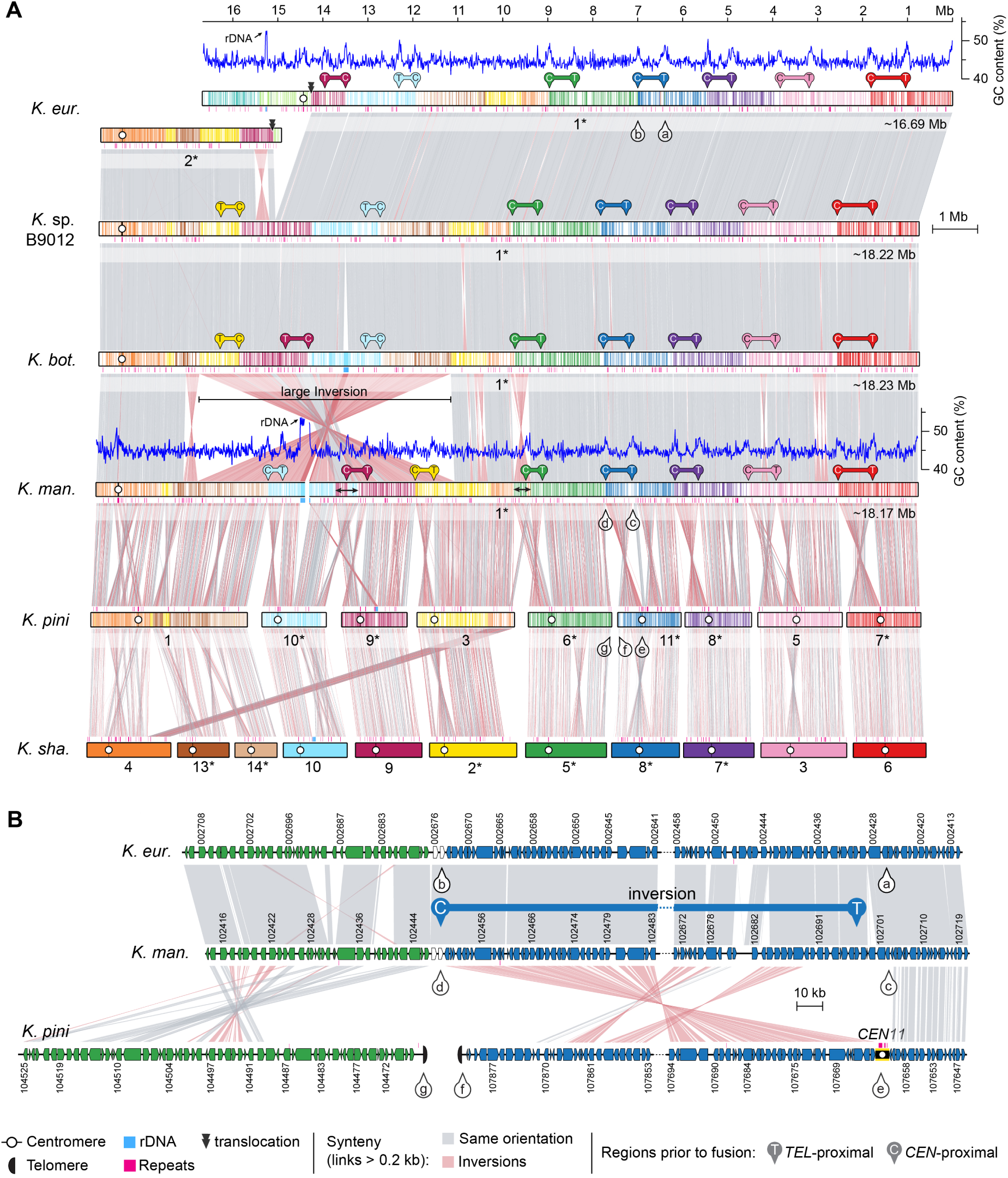
Giant chromosome formation in the ancestor of *K. mangrovensis* and sibling species resulted from multiple chromosome fusions events and inversions between centromere and telomere proximal regions. **(A)** Synteny comparison showing that the giant chromosome of *K. mangrovensis*, *K. botswanensis*, *Kwoniella* sp. B9012 and *K. europaea* (chr. 1) resulted from fusion of 9 chromosomes extant in *K. pini* (equivalent to 11 ancestral chromosomes). Note that the centromere-proximal regions of *K. pini* chromosomes correspond to regions near the fusion points on the giant chromosomes, whereas the telomere-proximal regions match more internalized regions, suggesting that a large pericentric inversion targeting the centromere-adjacent region is associated with each fusion event. Small inversions in *K. mangrovensis* relocating the telomere- and centromere-proximal regions after fusion are marked by a double-sided black arrow, and a large inversion is predicted to have occurred after the split from *K. mangrovensis* but before divergence of the other three species. In *K. europaea* the progenitor giant chromosome subsequently underwent a translocation with chr. 2. Chromosomes inverted relative to their original assembly orientations are marked with asterisks. **(B)** Zoomed-in synteny view of the genomic regions marked in panel A (pins with lowercase letters from a – g), shown as an example.

### Chromosome fusion events are associated with inversions between centromere- and telomere-proximal regions

To elucidate how these chromosomal fusion events occurred, all resulting junction sites were inspected. First, the possible presence of interstitial telomeric repeat sequences were examined near the fusion sites, as they could have been retained as remnants of past telomere-to-telomere fusion events [42]. However, no telomeric arrays, consisting of a minimum of two repeats of the telomeric motif TAAC(4,5), were detected beyond the chromosome ends.

Next, by aligning the giant chromosome of *K. mangrove*nsis with the individual chromosomes of *K. pini*, we unexpectedly found that at each fusion point, one side had sequences matching the end of one of the fused chromosomes, while the sequences on the other side aligned with an internal region of the other chromosome, rather than its end (illustrated in **Fig 3B** for *K. pini* chrs. 6 and 11). Further examination revealed that these internal sequences are chromosomal regions adjacent to *in silico* predicted centromeres (**Fig 3B**), an arrangement that suggests large inversions between telomere- and centromere-proximal regions. This pattern, prevalent in most chromosomal fusion events contributing to the giant chromosome formation (**Fig 3A**), was also observed in fusion events across other *Kwoniella* species (**S4-S6 Figs**), demonstrating a widespread occurrence. Predictably, this pattern is more discernable in more recent chromosome fusion events or when comparing chromosomes of closely related species. In a few instances, subsequent post-fusion secondary inversions may have obscured this pattern (e.g., the fusion of chrs. 9-10 and 3-6 in the giant chromosome, **Fig 3A**; or the fusion at the origin of chr. 2 of *Kwoniella* sp. CBS9459, **S4A Fig**).

### *Kwoniella* species have significantly shorter centromeres

The finding of extensive karyotypic variation in *Kwoniella*, along with large inversions involving centromeric regions associated with chromosome fusion events, led us to inspect (i) the conservation of kinetochore components essential for accurate chromosome segregation, and (ii) experimentally validate *in silico* predicted centromeres in selected *Kwoniella* species with different chromosome numbers.

The centromere-specific histone H3 variant CENP-A, a key epigenetic marker for centromeres and kinetochore formation, previously characterized in *Cryptococcus* and other fungi [23, 24, 51, 62, 64, 67-71], was confirmed to be conserved in both *Cryptococcus* and *Kwoniella* species, together with most outer kinetochore proteins such as those of the KMN (Knl1, Mis12 and Ndc80 complexes) network and Dam1/DASH complex (**S7 Fig** and **S2 Appendix**). Despite most canonical inner kinetochore proteins being absent in the Agaricomycotina subphylum, to which *Cryptococcus* and *Kwoniella* belong, bridgin (Bgi1), which along with CENP-C connects outer kinetochore network to centromeric chromatin [72], was found in all species. Together, this indicates that machinery for accurate chromosomal segregation has been largely retained.

To characterize centromeres in *Kwoniella*, N-terminally mCherry-tagged CENP-A proteins were functionally expressed in *K. europaea* (3 chrs.), *K. bestiolae* (8 chrs.), and *K. pini* (11 chrs.). A genetic construct expressing the fusion protein from its native promoter was randomly inserted into each strain via biolistic transformation. Live cell imaging showed the mCherry-tagged proteins exhibit centromere localization patterns consistent with those reported in *Cryptococcus* species (**S7C Fig**) [23, 62], indicating that the mCherry-CENP-A alleles are functional. We also attempted to express these constructs in *K. shandongensis* (with 14 chrs.) but were unsuccessful in obtaining transformants. To identify functional centromeres, we performed CENP-A chromatin immunoprecipitation sequencing (ChIP-seq). Because *C. neoformans* and *C. deuterogattii* centromeric regions are also enriched for other epigenetic marks, including 5-methylcytosine (5mC) DNA methylation, and heterochromatic histone modification H3K9me2 [62-64, 73, 74], ChIP-seq with an antibody specific to H3K9me2 and whole-genome bisulfite-sequencing (WGBS) were also conducted.

A single CENP-A bound region was significantly enriched on each of the chromosomes of the three *Kwoniella* species (**Fig 4**, and **S8**-**S10 Figs**), matching the centromeric regions initially predicted by synteny analysis, thereby validating for our *in-silico* approach. All centromeres also exhibited enrichment for H3K9me2, but the presence of 5mC was less consistent. For instance, high levels of 5mC were detected across all centromeres of *K. europaea* (**S8 Fig**), but only on *CEN3*, *CEN4*, and *CEN7* of *K. bestiolae* (**S9 Fig**). In *K. pini*, 5mC was absent from all centromeres, despite being present in a few other genomic regions (**S10 Fig**). Notably, this variation was not strictly correlated with the presence/absence of transposable elements (TEs) as *K. pini* centromeres still retain some TE remnants (**S10 Fig**). Building on recent research [73], we have also determined that all *Kwoniella* species possess both *de novo* and maintenance-type DNA methyltransferases (Dnmt5 and DnmtX) for cytosine DNA methylation, unlike all *Cryptococcus* species, including *Cryptococcus* sp. OR918 representing the earliest-branching lineage of this group, which only retained Dnmt5, and *C. depauperatus* that has specifically lost both proteins (**S11 Fig** and **S2 Appendix**). An independent loss event of DnmtX is also noted in the outgroup species, *Bullera alba* (**S11 Fig**). Considering all of the evidence, we conclude that the genomic regions identified in *Kwoniella* serve as binding sites for the centromeric histone CENP-A, affirming their role as *bona fide* centromeres.

**Fig 4.**
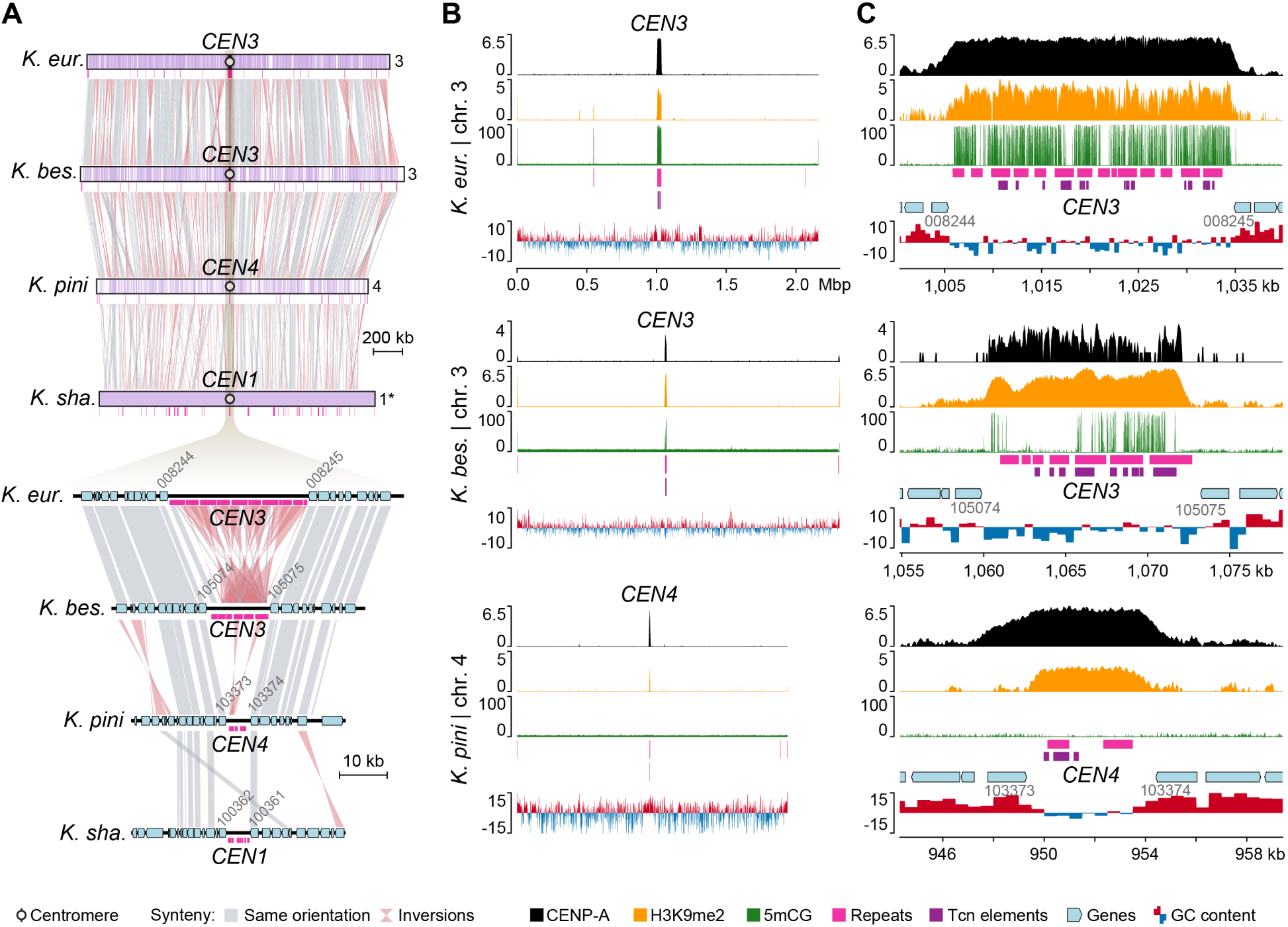
Experimental validation of *Kwoniella* centromeres in three species with different numbers of chromosomes. **(A)** Gene synteny conservation spanning a predicted centromeric region in four *Kwoniella* species (*CEN3* of *K. europaea*, *CEN3* of *K. bestiolae*, *CEN4* of *K. pini*, and *CEN1* of *K. shandongensis*). **(B)** Plots of the chromosomes depicted in panel A (except for *K. shandongensis*) displaying CENP-A (black) and H3K9me2 (orange) enrichment, fraction of CG cytosine DNA methylation (5mCG, green), repeat content (pink), Tcn-like LTR elements (purple), and GC content (shown as deviation from the genome average – red, above; blue, below). The fold enrichment of each sample over the input DNA is shown on the left of each panel for CENP-A and H3K9me2. **(C)** Zoomed-in sections show the regions spanning the centromeres with adjacent genes (light blue). In panels B and C, data is computed in 5-kb non-overlapping windows.

As established in *C. neoformans*, *C. deneoformans*, *C. deuterogattii*, and *C. amylolentus* [23, 62, 64] and here demonstrated for three *Kwoniella* species, the lengths of the CENP-A-bound regions largely coincide with the ORF-free regions predicted to be centromeres. As such, the distance between centromere-flanking genes was leveraged as a metric to quantify centromere length across species. This analysis revealed significantly shorter centromeres in *Kwoniella* compared to *Cryptococcus* (*P* < 0.0001, Mann-Whitney U Test), with *Kwoniella* centromeres averaging 6.7 kb in median length (mean 8.3 kb) versus *Cryptococcus* centromeres that have a median length of 32.9 kb (mean 40.3 kb) (**Fig 5A** and **S1 Appendix**).

**Fig 5.**
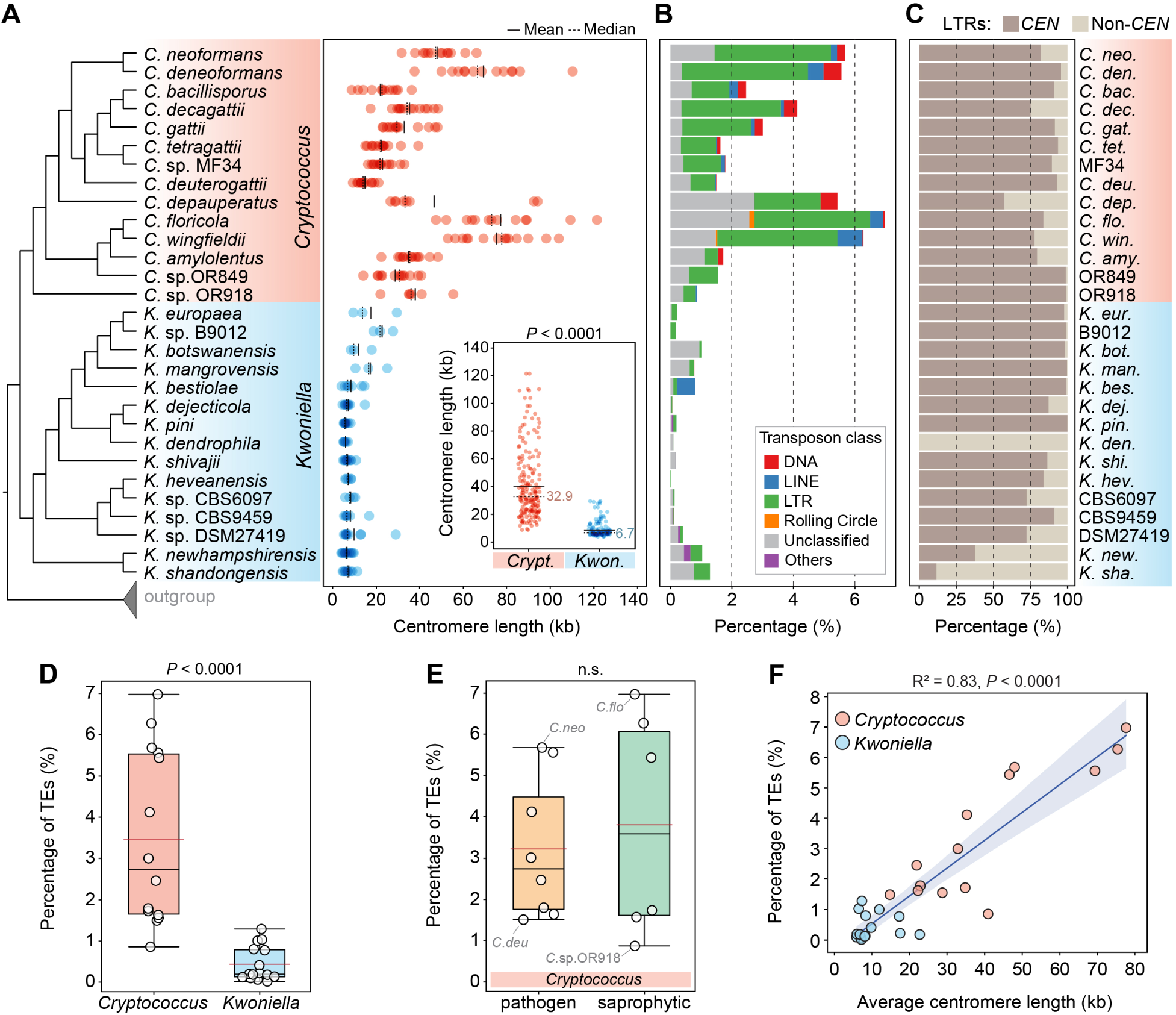
Centromere length and transposable element (TE) content in *Cryptococcus* and *Kwoniella*. **(A)** Comparison of estimated centromere lengths (in kb) along the phylogeny of *Cryptococcus* and *Kwoniella* species. Each dot represents a single centromere, and solid and dashed black lines represent mean and median lengths, respectively. A comparison between the two groups is summarized in the inset, showing significantly smaller centromeres in *Kwoniella* compared to the *Cryptococcus* lineage (*P*-value obtained by Wilcoxon’s Test). **(B)** Estimated TE content within each genome. **(C)** Relative percentage of LTR retrotransposons found in centromeric (*CEN*) versus non-centromeric (non-*CEN*) regions (normalized by the total percentage of LTRs). **(D-E)** Box-plots comparing TE content between *Cryptococcus* and *Kwoniella*, and between pathogenic and non-pathogenic *Cryptococcus* species (*P* values obtained by Mann-Whitney U Test; n.s., not significant). The red line, black line, and boxes denote the mean value, median value, and interquartile range, respectively. **(F)** Correlation between TE abundance and average centromere length. The plot shows a linear regression with confidence values (blue area) of the fitted line, *R^2^*correlation coefficients values, and *P* values from the *t*-test for the slope of the regression line.

### *Kwoniella* centromeres exhibit markedly fewer TEs aligned with reduced genomic TE load

Although centromere identity and function are typically defined by CENP-A binding rather than by specific DNA sequences, repetitive sequences, such as TEs, are frequently observed in the centromeres of plants, animals, and fungi [75-77]. As established in previous studies, *Cryptococcus* species exhibit large regional centromeres that are enriched in specific long-terminal-repeat retrotransposons (LTRs) from the Ty3/*Gypsy* and Ty1/*copia* families [23, 24, 62]. It is thus not surprising that a comparative analysis of repetitive elements across the two groups revealed a significantly higher prevalence of TEs in *Cryptococcus* species (*P* < 0,0001, Mann-Whitney U Test), accounting for ∼0.9 to 7% of the genome (median 2.7%). On the other hand, *Kwoniella* species contain substantially fewer elements, ranging from 0.02 to 1.3% (median 0.2%) (**Figs 5B** and **5D**). Among the pathogenic *Cryptococcus* clade, *C. neoformans* and *C. deneoformans* are the two species situated at the higher end of transposon density (estimated at ∼5.7 and 5.6%, respectively) while *C. deuterogattii* represents the lower end (∼1.5%). No substantial differences in transposon load were found between pathogenic and non-pathogenic *Cryptococcus* species (**Fig 5E**; *P* = 0.95, Mann-Whitney U Test).

Interesting, a positive correlation was detected between TE content and the average length of centromeres (*P* < 0.0001, *R*^2^ = 0.814; **Fig 5F**), with most LTRs being located within the predicted centromeric regions in both groups (**Fig 5C**). However, *K. dendrophila* deviates from this pattern; this species exhibits low TE density genome-wide, and the predicted centromeres are devoid of LTRs and 5mC DNA methylation, while still exhibiting enrichment for H3K9me2 (**S12 Fig**). Additional exceptions were detected in *K. shandongensis* and *K. newhampshirensis*, as in these species, remnants of LTRs are only present in a subset of the predicted centromeres, along with other unclassified repeat elements (**S13 Fig**). These findings highlight the complex and diverse nature of centromere structures, emphasizing significant genetic and epigenetic characteristics of these critical chromosomal regions, even among closely related species.

### Shorter centromeres in *Kwoniella* do not appear to be a result of RNAi loss

Previous studies in *C. neoformans and C. deneoformans* identified centromeres as primary sources for production of small-interfering RNA (siRNA) for transposon silencing via the RNA interference (RNAi) pathway [62], with these species retaining the canonical RNAi components: Argonaute (Ago), Dicer (Dcr), and RNA-dependent RNA polymerase (Rdp) [78, 79]. In contrast, the RNAi-deficient species *C. deuterogattii* has lost many RNAi components [80] and harbors shortened centromeres containing only LTRs remnants [62]. This led us to question whether *Kwoniella* shortened centromeres were due to a lack of active RNAi.

To address this, we conducted an in-depth analysis of the key RNAi genes across *Cryptococcus* and *Kwoniella*, which revealed that the common ancestor of the two lineages likely had a functional RNAi pathway, with two Argonaute proteins (Ago1 and Ago4), one Dicer (Dcr1), and one RNA-dependent RNA polymerase (Rdp1) (**S14 Fig**). Additionally, we uncovered a complex pattern of gene duplication and loss across lineages (refer to **S1 Text** for details), yet, excluding *C. deuterogattii*, all species seem to be RNAi-proficient based on the presence of canonical RNAi genes (**S14 Fig**). Beyond the core components of the RNAi-pathway, we also examined eight additional genes (*ZNF3 GWC1*, *QIP1*, and *RDE1* to *RDE5*) essential for global siRNA production in *C. neoformans* [80-82] (**S1 Text** and **S2 Appendix**). While most genes were consistently conserved, *ZNF3* (CNAG_02700) was an exception, having been lost independently three times in *Cryptococcus* (**S1 Text**, **S15 Fig**, and **S2 Appendix**). In *Kwoniella*, Znf3 proteins are shorter but retain essential zinc finger domains, suggesting potential functionality (**S1 Text**, **S15 Fig**, and **S2 Appendix**). These findings suggest that *Kwoniella* retains both canonical and auxiliary RNAi components, pointing to an active RNAi pathway. Thus, the shorter centromeres in *Kwoniella* likely result from factors other than RNAi pathway deficiencies, underscoring the need for further research into factors influencing centromere length.

### Identification of mini-chromosomes in *Kwoniella*

Our genomic analyses also detected unusually small chromosomes (< 100 kb) in some *Kwoniella* species, which we termed “mini-chromosomes”. Specifically, six mini-chromosomes were identified in *Kwoniella* sp. DSM27419 (ranging from ∼43.7 to 83 kb), two in *K. newhampshirensis* (∼39 kb each), and one in *K. shivajii* (∼87.6 kb). This was substantiated by PFGE and read coverage analyses from both Illumina and long-read sequencing, along with the identification of telomeric repeats at both ends of each mini-chromosome (**S16A-B Figs**). Compared to regular chromosomes, these mini-chromosomes have a lower gene density and reduced GC content (**S16C-H Figs**). Interestingly, *Kwoniella* sp. DSM27419 has an extra chromosome (chr. 12) ∼228 kb in size, and *Kwoniella* sp. CBS9459 also features a smaller chromosome (chr. 6) ∼745 kb. Both chromosomes show no synteny with other *Kwoniella* chromosomes (**Fig. 2** and **S16F Fig**), leaving their origin and function currently unclear and subject to further investigation.

### Chromosome number reduction in *Cryptococcus* via centromere inactivation and transposon loss

Unlike the broad spectrum of chromosome numbers in *Kwoniella*, *Cryptococcus* species maintain a more uniform chromosome number, with only two species having fewer than 14 chromosomes (*C. depauperatus* and *Cryptococcus* sp. OR918, **Fig 1B**). To investigate if chromosome number reduction in *Cryptococcus* follows mechanisms similar to those identified in *Kwoniella* (chromosome fusions with pericentric inversions), we focused on comparisons between *C. depauperatus* (8 chromosomes), and two other *Cryptococcus* species, *C. neoformans* and *C. amylolentus* (each with 14 chromosomes).

Synteny analysis revealed substantial interchromosomal rearrangements in these species after diverging from their last common ancestor (**Fig 6A**). Analysis of the centromere-flanking regions indicates that *C. depauperatus* centromeres have served as frequent target sites for chromosomal arm exchanges through intercentromeric recombination, similar to *C. neoformans* and *C. amylolentus* [23, 62]. However, whereas only three centromere-mediated translocations are estimated since the last shared ancestor of *C. neoformans* and *C. amylolentus*, in *C. depauperatus*, 7 out of 8 extant centromeres appear to be products of repeated intercentromeric recombination (**Fig 6B**).

**Fig 6.**
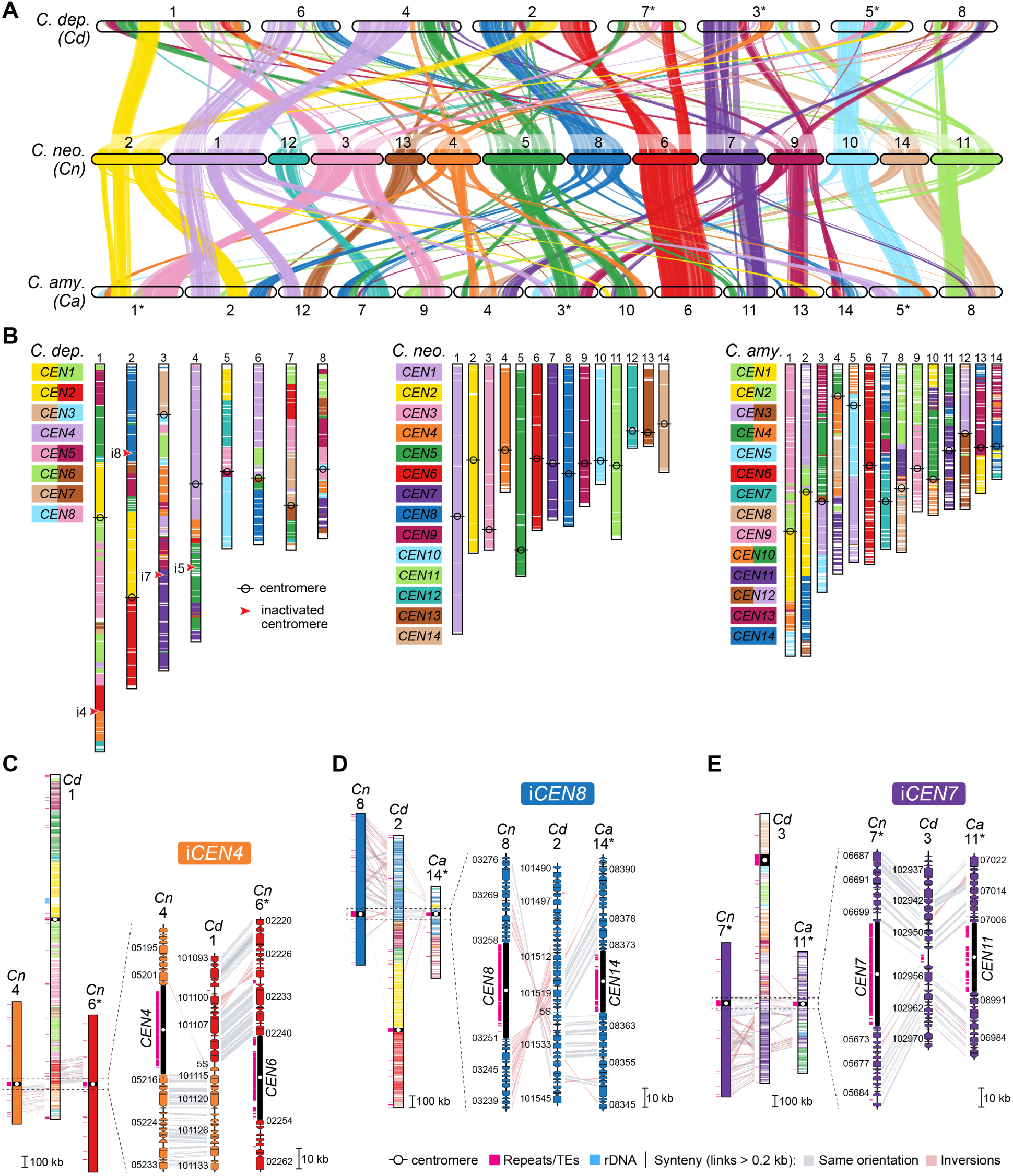
Chromosome number reduction in *Cryptococcus* involves centromere inactivation via loss of LTR-rich regions. **(A)** Pairwise synteny relationships between *C. depauperatus* (8 chromosomes), and *C. neoformans* and *C. amylolentus* (14 chromosomes each). Links represent the boundaries of syntenic gene blocks identified by MCScanX with pairwise homologous relationships determined by SynChro. Chromosomes are color-coded per *C. neoformans* and were reordered or inverted (marked with asterisks) from their original assembly orientations to maximize collinearity. **(B)** Overlay of synteny blocks and centromere locations, underscoring several intercentromeric rearrangements between *C. neoformans* and the other species. Red arrowheads pinpoint predicted inactivated centromeric regions. (**C-E**) Linear chromosome plots depicting gene synteny conservation between *C. depauperatus* (*Cd*), *C. neoformans* (*Cn*), and *C. amylolentus* (*Ca*) chromosomes in regions corresponding to inactivated *CEN4* (i*CEN4*), *iCEN8,* and *iCEN7* in *C. depauperatus*, respectively.

Synteny analysis also suggests that the reduction from 14 to 8 chromosomes in *C. depauperatus* did not result from sequential chromosome fusions as in *Kwoniella*. This chromosome number reduction involved inactivating or losing 6 centromeres. We traced the fate of 4 of these centromeres, aligning with *CEN4*, *CEN5*, *CEN7*, and *CEN8* of *C. neoformans*, with reasonable accuracy (**Fig 6B**). The other two, linked to recombined *CEN6*/*CEN9* and *CEN12* in *C. neoformans*, seem to have been lost due to additional rearrangements. Our previous work reported that the inactivation of the centromere in *C. depauperatus*, aligning with *CEN5* in *C. neoformans* (or *CEN10*/*CEN4* in *C. amylolentus*), likely resulted from gross chromosomal rearrangements linked to the complex evolution of the mating-type locus [11]. Consequently, we focused on losses in *C. depauperatus* corresponding to *CEN4*, *CEN7*, and *CEN8* of *C. neoformans*.

Examination of the gene content in *C. depauperatus* that aligns with *C. neoformans CEN4* suggests that inactivation of this centromere might have resulted from intercentromeric recombination followed by loss of the LTR-rich region. This is supported by the gene organization in C. *depauperatus*, which aligns well with a juxtaposition of the genes flanking the opposite sides of *CEN4* and *CEN6* in *C. neoformans* (**Fig 6C**). Similarly, inactivation of *CEN7* and *CEN8* appears to be associated with the removal of their LTR-rich regions, without major loss of flanking genes or rearrangements (**Figs 6D** and **6E**). Notably, the *C. depauperatus* region corresponding to inactivated *CEN7* still contains TE remnants, suggesting it may represent a more recently inactivated centromere. These findings collectively indicate that reduction in chromosome number in *Cryptococcus* resulted from chromosomal rearrangements other than full-chromosome fusions, and centromere loss is primarily attributable to excisions of centromeric DNA sequences.

### Surveying genomic signatures of karyotypic variation

In light of the marked karyotypic differences between *Cryptococcus* and *Kwoniella*, along with the unique chromosome dynamics in the *Kwoniella* clade, we leveraged our high-resolution genome assemblies to identify potential variations in gene families related to telomere length maintenance and protection (shelterin complex). To examine gene conservation, orthologs were identified across 32 genomes (14 *Cryptococcus*,15 *Kwoniella*, and 3 outgroup species) (**Fig 7**). A total of 4,756 orthogroups (OGs) were found in all the *Cryptococcus* and *Kwoniella* genomes. Key telomere-length regulation proteins such as Tel2, Tel1, Tert, and Blm helicase were conserved across species (**S2 Appendix**). However, except for *POT1*, *TPP1*, and a *TEN1*-like gene, no other clear shelterin complex orthologs were found (see **S2 Text** for details and **S2 Appendix**). This suggests either that other shelterin components are absent, or that other proteins fulfilling these roles have evolved, in line with the reported diversity of fungal shelterin proteins [83-85]. Additionally, the analysis of 152 genes known to impact telomere length [86, 87] in the budding yeast *Saccharomyces cerevisiae* also showed a high degree of conservation (**S2 Appendix**), with an average of ∼82% of these genes present across species. Significantly, species with “giant” chromosomes, showed a similar conservation rate, highlighting the lack of major gene composition differences across species with varied chromosome numbers (**S2 Text** and **S2 Appendix**).

**Fig 7.**
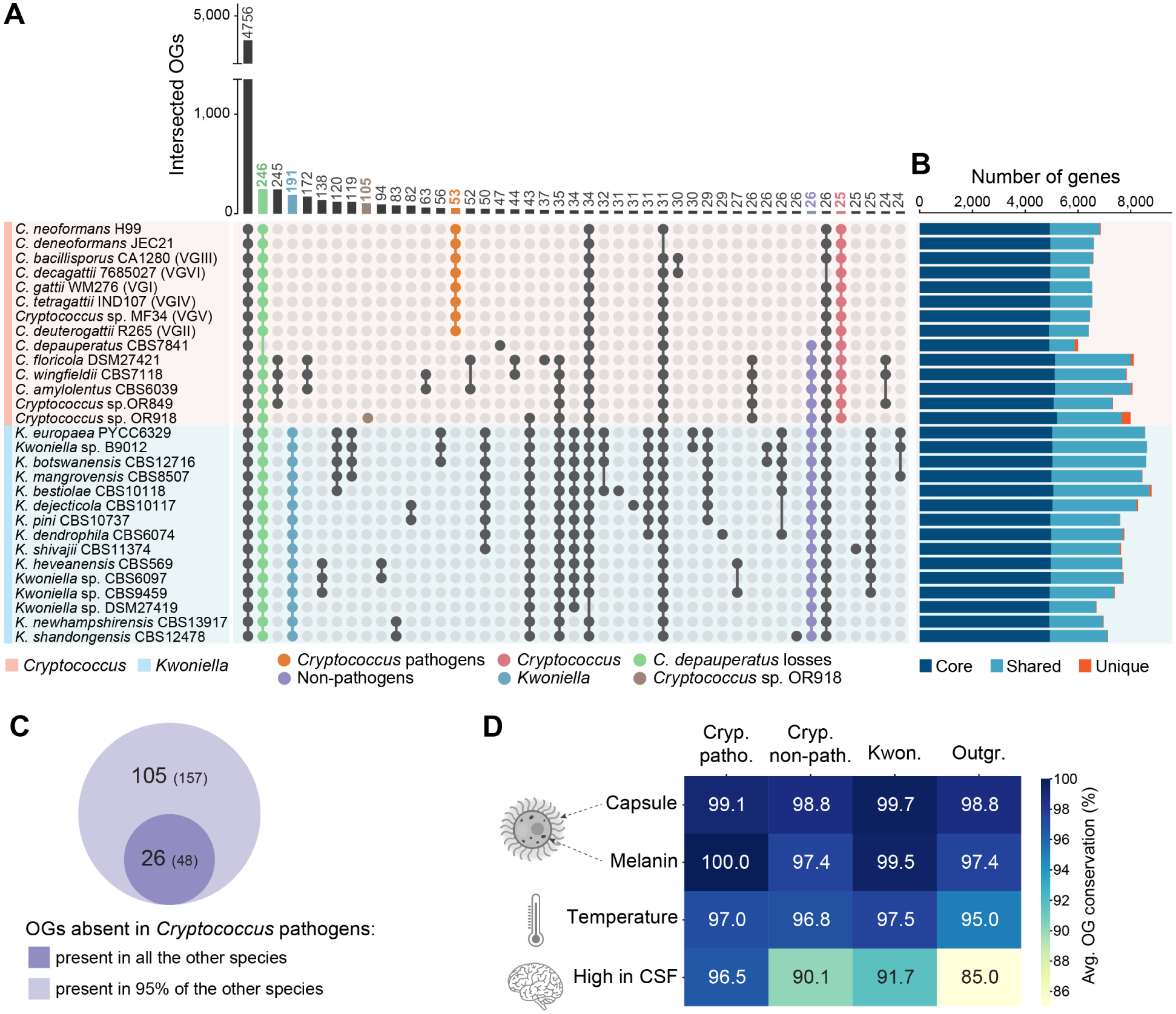
Gene family analysis in *Cryptococcus* and *Kwoniella*. **(A)** UpSet plot displaying protein family overlaps between *Cryptococcus* and *Kwoniella*. The numbers of protein families (orthogroups, OGs) defined by OrthoFinder are indicated for each species intersection, with key intersections emphasized as per the accompanying key. **(B)** Bar plot categorizing orthology classes into core genes (found in all genomes; dark blue), shared genes (present in multiple but not all genomes; light blue), and unique genes (present within species-specific OGs; dark orange). **(C)** Venn diagram comparing OGs absent in all pathogenic *Cryptococcus* species but present in all or 95% of non-pathogenic *Cryptococcus* and *Kwoniella* species. **(D)** Heatmap depicting the average conservation rate in *Cryptococcus* pathogens, non-pathogenic *Cryptococcus*, *Kwoniella*, and outgroup species, of OGs identified in *C. neoformans* known to be involved in capsule and melanin production, growth at 37 °C, and high expression in human cerebrospinal fluid (CSF).

To examine more closely if differences in selection may have contributed to, or resulted from, karyotypic variation, we carried out a branch selection analysis. We compared the evolutionary patterns of orthologous genes in four *Kwoniella* species with 3 chromosomes (foreground branch) to a sister clade of four species with 8 to 11 chromosomes (background branch). Genes in the foreground clade with higher dN/dS ratios compared to the background clade indicate a greater degree of selection. Top scoring genes included those associated with cell structure, ribosomal functions, and DNA modification or repair (**S3 Appendix**). Two genes related to centromere function in the top 30 corresponded to *C. neoformans* CNAG_02218, a putative homolog of *SCM3* (suppressor of chromosome mis-segregation) [88, 89] and CNAG_01334, an ortholog of *S. cerevisiae SLI15*, encoding a subunit of the Chromosomal Passenger Complex (CPC) that regulates kinetochore-microtubule interactions and is important for chromosome segregation [90, 91]. These findings may suggest a model in which instability in chromosome segregation could be a driving event in karyotype instability.

### Gene family losses in pathogenic and non-pathogenic *Cryptococcus* species

Complete genomes sequences of *Cryptococcus* and *Kwoniella* species also provide a new window into the evolution of their gene content. Beyond the 4,756 shared OGs between both groups, the next most frequent pattern is the specific absence of 246 OGs in *C. depauperatus*, which has the smallest gene set (**Fig 7, S1 Fig, S1 Appendix,** and **S4 Appendix**). This substantial gene loss in *C. depauperatus* is aligned with its reduced genome size (**Fig 1**, **S1 Fig** and **S1 Appendix**) and possibly associated with loss of yeast phase growth [11]. Notable gene losses include major facilitator superfamily proteins, glycosyl hydrolase family 3 proteins, and DNA-interacting proteins, such as Msh4 and Msh5 mismatch repair proteins, and the Rad8 DNA repair protein. Also missing are genes encoding uracil/uridine permeases (CNAG_04632 and CNAG_07917) (**S4 Appendix**), aligning with our own experimental observations that no 5-FOA resistant mutants could be isolated, suggesting an inability of *ura5* mutants to import uracil or uridine to compensate for auxotrophy.

In analyzing genes absent in all pathogens yet present in all non-pathogens, we identified 26 OGs, corresponding to 48 genes in non-pathogenic *C. floricola*. These genes displayed diverse enzymatic activities, including a large set of short chain dehydrogenase/reductases and amino acid permeases (**Figs 7A** and **7C**, and **S4 Appendix**). Recognizing that some gene losses in pathogens might also sporadically occur in non-pathogens due to environmental adaptations or specific organismic interactions, we adjusted our criteria to include genes absent in all pathogens but present in at least 95% of non-pathogens. This adjustment resulted in identifying 79 additional OGs (109 genes in *C. floricola*) (**Fig 7C**). Two genes, *PRA1* and *ZRT1*, stand out for their roles in zinc acquisition in *C. albicans*, with the loss of *PRA1* being recently suggested as a possible key evolutionary step for fungal pathogenesis [92]. Pra1, a zincophore [93], and Zrt1, a zinc transporter [94], are adjacent and divergently transcribed, and this arrangement was found to predate the Basidiomycota-Ascomycota split, with *PRA1* having subsequently experienced multiple losses across fungal clades [95, 96]. The absence of the *PRA1/ZRT1* cluster was previously noted in some pathogenic *Cryptococcus* species, contrasting with six non-pathogenic species surveyed that retain it [94]. We have now broadened this analysis by confirming the absence of this gene cluster in all pathogenic *Cryptococcus* species, and its presence in most, though not all, non-pathogenic *Cryptococcus* and *Kwoniella* species (**S16A Fig**). Exceptions include *Cryptococcus* sp. OR849 and *K. newhampshirensis*, which have recently lost the cluster as inferred from synteny analysis (**S16A, F-G Figs**). Additionally, *Cryptococcus* sp. OR918 and most (13 of 15) *Kwoniella* species possess a second *PRA1* variant (*PRA1-2*) with significant sequence differences and no adjacent zinc transporter (**S16B-C Figs**). The absence of *PRA1*/*ZRT1* gene cluster in pathogenic species, and its selective loss in non-pathogens hinting at a pre-pathogenic state, underscore its potential significance in fungal pathogen evolution, warranting further investigation.

### Gene families in pathogenic *Cryptococcus*

Among the genes specifically present in the *Cryptococcus* pathogens (53 OGs, corresponding 57 *C. neoformans* genes), there is a large group predicted to interact with DNA, including six predicted transcription factors or DNA binding proteins (**S4 Appendix**) and two of the four genes annotated as B-glucuronidases (PFAM glycosyl hydrolase family 79C), implicated in cell wall modification [97]. As strong signatures of pathogenesis were not detected in this analysis, the conservation of genes associated with the canonical features of cryptococcal pathogenesis were evaluated. Genes associated with capsule biosynthesis [98, 99], melanin production [100-104], and ability to grow at 37 degrees [105] were all found to be highly conserved across all species (**Fig 7D**, **S5 Appendix**). Only a gene set identified as highly expressed in human cerebrospinal fluid (CSF) across diverse *C. neoformans* strains [106] was found to be less conserved in non-pathogenic *Cryptococcus*, *Kwoniella*, and the outgroup species (**Fig 7D**, **S5 Appendix**).

Gene family analysis also found an exclusive gene in pathogenic *Cryptococcus*, residing ∼82 kb from the right end of chr. 1 in *C. neoformans* H99, predicted to encode a D-lactate dehydrogenase, expressed under various conditions (see CNAG_00832 in FungiDB) (**Fig 8**). Surprisingly, BLAST analysis shows its closest homologs are *Aspergillus* proteins with about 87% identity, indicating possible horizontal gene transfer (HGT) from these ascomycetous molds. Investigating further, we used the CNAG_00832 protein sequence to search the NCBI clustered nr database and conducted phylogenetic analysis of the top 1,000 hits. The resulting tree (**Fig 8A**) places *Cryptococcus* proteins within the genus *Aspergillus* (**Fig 8B**), while also suggesting a bacterial origin for this fungal D-lactate dehydrogenase. In line with this, AlphaFold 3D structure predictions following pairwise structure alignments revealed similar structures among three representative sequences of each of the three lineages (**Fig 8D** and **S18 Fig**). The precise function of this gene in *Cryptococcus* pathogens, particularly in interconverting D-lactate and pyruvate, along with the reduction/oxidation of NAD+ and NADH, is subject of ongoing work, but could be important for *Cryptococcus* growth in glucose-limited environments like the brain during cryptococcal meningoencephalitis, potentially via gluconeogenesis [107].

**Fig 8.**
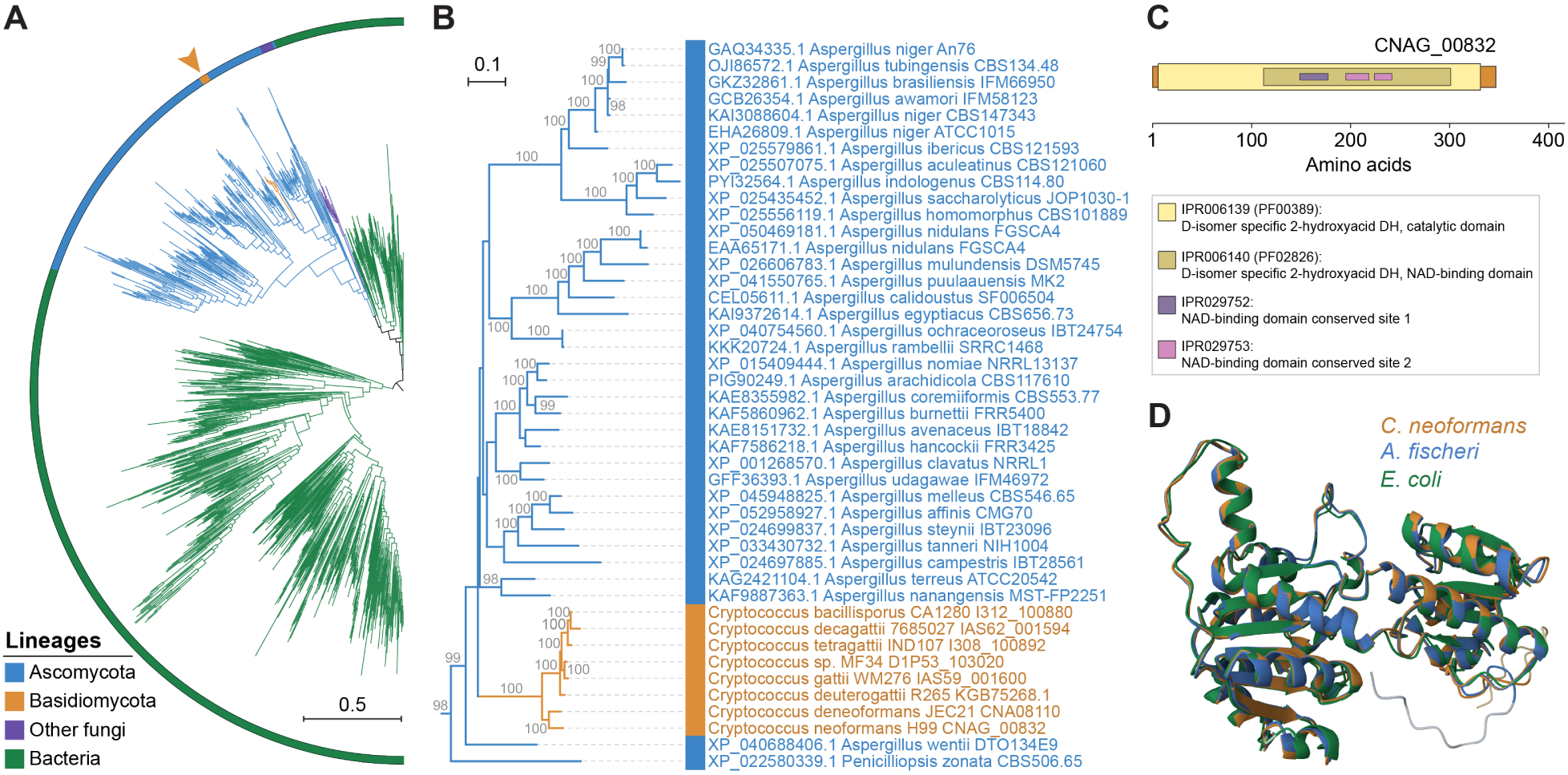
Horizontal gene transfer and the origin of a putative D-lactate dehydrogenase specific to pathogenic *Cryptococcus* species. **(A)** Maximum likelihood (ML) phylogeny encompassing 1,007 protein sequences obtained from a BLASTP search against the NCBI clustered nr database, using the *C. neoformans* CNAG_00832 protein sequence as query and selecting the top 1,000 hits. The identified sequences in pathogenic *Cryptococcus* species were also included. Tree branches are colored as per the key, depicting major groups of organisms. The tree, visualized with iTOL v6, was constructed with IQ-TREE2, with internal branch support assessed by 10,000 replicates of Shimodaira–Hasegawa approximate likelihood ratio test (SH-aLRT) and ultrafast bootstrap (UFboot), and is rooted at the midpoint. Branch lengths are given in number of substitutions per site. **(B)** Pruned ML phylogeny showing the position of *Cryptococcus* proteins clustering within the presumed Aspergilli donor lineage. **(C)** Domain organization of the protein sequence encoded by CNAG_00832 with identified InterPro (IPR) and Pfam (PF) domains highlighted. **(D)** Overlay of AlphaFold predicted structures of D-lactate dehydrogenease proteins from *C. neoformans* (UniProt J9VFV7), *Aspergillus fischeri* (UniProt A1D163), and *Escherichia coli* (UniProt P52643) showing structural similarity. Pairwise structure alignments were performed and visualized in the Protein Data Bank (PDB) website (https://www.rcsb.org/alignment) using the JjFATCAT-rigid algorithm.

## Discussion

Our broad comparative genomics analysis of *Cryptococcus* and *Kwoniella* has uncovered extensive karyotypic variation, particularly within *Kwoniella*, and significant gene content conservation in pathogenic and non-pathogenic species. Our findings support a non-pathogenic common ancestor, with pathogenic traits evolving more recently in *Cryptococcus*. While pathogenic *Cryptococcus* species display distinct phenotypic traits, including higher temperature tolerance and melanin and capsule virulence factors [10], genes responsible for these characteristics are largely conserved across all species. This suggests that gene sets enabling adaptation to diverse environmental niches might predispose certain species to develop pathogenicity in humans. Echoing the minimal gene content differences between humans and great apes [108], our study indicates that differences in pathogenic *Cryptococcus* species may stem from finer scale gene variation or differential genetic and epigenetic regulation, rather than major changes in gene content. While it seems plausible that shared traits underlie emergence of pathogenic species, it is also likely that finer differences will emerge within the pathogens given that some species (*C. neoformans*, *C. deneoformans*, *C. bacillisporus*, *C. tetragattii*) predominantly infect HIV/AIDS patients whereas others (*C. gattii* and *C. deuterogattii*) infect largely non-HIV/AIDS patients, such as those with autoantibodies to the cytokine GM-CSF [109-112]. Understanding these differences will require incorporating further population, transcriptomic, epigenetic, and functional data.

The notable plasticity of fungal genomes, particularly evident in *Cryptococcus* species, is highlighted by a variety of mechanisms and processes that drive genetic variation (such as hybridization [113-115], ploidy variation [116, 117], transposon mobilization [118, 119], gene loss [120], and gene gain via duplication [121] or horizontal gene transfer [122]) and also by diverse reproductive strategies (e.g. [123]). Our analysis found a few, but likely significant gene losses, especially within *Cryptococcus*. First, we expanded the previously reported loss in *Cryptococcus* of a *de novo* methyltransferase (DnmtX) [73] involved in 5mCG methylation, by identifying its absence in an earlier-derived *Cryptococcus* lineage (strain OR918). Contrastingly, DnmtX persists across *Kwoniella* species. Alongside the ubiquitous presence of a maintenance methyltransferase (Dnmt5) in both genera (with *C. depauperatus* as an exception, lacking both genes), this suggests the possibility of distinct DNA methylation landscapes in the two genera, potentially impacting gene regulation and TE control. Secondly, in examining the genetic network encoding RNAi components across the two groups, we uncovered a complex evolutionary history characterized by both ancestral and recent, species/clade-specific, gene duplications and losses (*AGO1/2/3/4*, *DCR1/2*, and *RDP1/2*). Besides *C. deuterogattii*, which uniquely lost multiple RNAi genes [80], these genes are otherwise retained in all other species, suggesting they have active RNAi pathways. This is of interest given the dramatic differences in centromere length between *Cryptococcus* and *Kwoniella*, and previous studies associating RNAi loss and centromere length contractions [62]. This may imply alternative mechanisms in *Kwoniella* for transposon control and centromere length regulation, as discussed further below. Interestingly, *Kwoniella* has maintained a higher number of Argonaute genes since diverging from a shared ancestor with *Cryptococcus*, which might indicate redundancy or functional specialization as observed in other organisms [124, 125]. The third example of gene loss is in the unusual species *C. depauperatus*, which lacks a yeast phase and grows exclusively as a hyphal organism engaged in continuous sexual reproduction [11]. This species has lost genes for uracil/uridine import and DNA repair, underscoring distinct evolution. Finally, our analysis broadens previous findings on the *PRA1/ZRT1* gene cluster, implicated in zinc acquisition and recently hypothesized as a key factor in fungal pathogen evolution [92]. We show this gene cluster was lost in all pathogenic *Cryptococcus* species, yet present in most, but not all, non-pathogens. This dichotomy indicates potential evolutionary pathways that could have facilitated pathogenic trait development in *Cryptococcus*, aligning with recent hypotheses on pathogenesis evolution in fungi [8].

Our analysis provides evidence that gene gain via HGT might have contributed to pathogenicity evolution in *Cryptococcus*. Specifically, we identified a gene encoding a putative D-lactate dehydrogenase that is unique to *Cryptococcus* pathogenic species and appears to have been acquired from the genus *Aspergillus*. While the function of this *Cryptococcus* gene is still under investigation, it could promote growth via gluconeogenesis during glucose deprivation (common during brain infections) [126, 127]. This aligns with proposed hypotheses that lactate, a C_3_ substrate, might be more favorable for biomass production in brain infection than C_2_ substrates such as acetate [107]. Supporting this, the glyoxylate shunt pathway, which uses C_2_ substrates, is not essential for *C. neoformans* virulence [128, 129]. These findings could suggest that *Cryptococcus* pathogens may have a pre-existing advantage in glucose-limited environments, such as the human brain. However, given that the *Cryptococcus* pathogenic group predates their interaction with humans, these traits might have stemmed from broader environmental pressures rather than solely due to human host interactions. Further studies of gene loss/gain, allied with comparative transcriptomic analyses under various conditions, will be needed to unravel other genetic and regulatory changes leading from non-pathogenicity to pathogenicity in these fungi.

Our extensive comparative genomic analysis also identified distinct karyotypic evolution in *Kwoniella* compared to *Cryptococcus*. In *Kwoniella*, chromosome fusion is the major driving force, occurring repeatedly and independently throughout the genus. Starting from an inferred ancestral chromosome number of 14, various extant *Kwoniella* species show reduced chromosome numbers – 11, 8, 5, and as few as 3 – due to successive chromosome fusions. At the extreme, these fusions have formed giant chromosomes, 15-18 Mb in size, which are up to 8 times larger than other extant chromosomes, comprising as much as 80% of the genome. Investigating what drove repeated chromosome-chromosome fusion events in *Kwoniella*, we observed that each fusion typically involves a pericentric inversion extending from one telomere to just beyond the centromere. As shown in **Fig 9**, these pericentric inversions may have occurred before (model A) or after chromosome fusion (model B).

**Fig 9.**
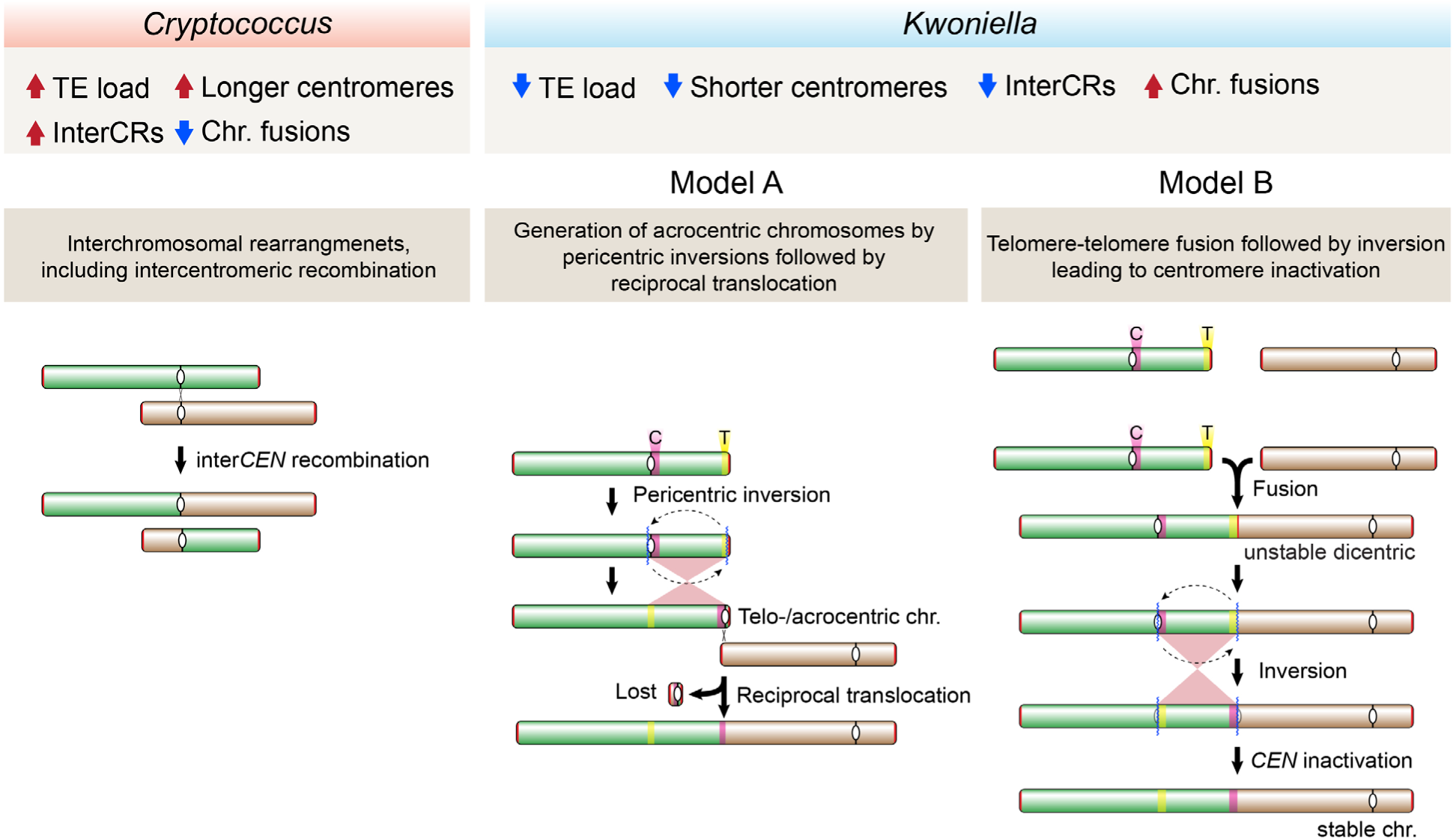
Proposed model of chromosome evolution in *Cryptococcus* and *Kwoniella*. The hypothesized mechanisms driving chromosome evolution in *Cryptococcus* and *Kwoniella* account for the distinct patterns of chromosomal rearrangements observed in these two groups. *Cryptococcus* (left), is characterized by frequent interchromosomal rearrangements, including intercentromeric recombination likely influenced by longer centromeres and higher transposable element (TE) content. *Kwoniella* (right), however, predominantly shows chromosome fusions, associated with lower TE content and shorter centromeres, and fewer interchromosomal rearrangements. Two mechanisms for chromosome fusions in *Kwoniella* are proposed: Model A, involving a pericentric inversion followed by a reciprocal translocation between chromosomes, typically resulting in the loss of the smaller translocation product; and Model B, based on "end-to-end fusion" leading to a dicentric chromosome that becomes monocentric through a pericentric inversion (see text for details).

Model A proposes an initial pericentric inversion shifting the centromere of one chromosome to the end, resulting in a telo- or acrocentric chromosome. This is followed by a symmetric reciprocal translocation with another chromosome, involving breakpoints near the centromere on the long arm of the telo- or acrocentric chromosome and near the end of the second chromosome. The resulting larger translocation product, combining most of both chromosomes, is retained, while the smaller product comprising the centromere of the telo- or acrocentric chromosome plus two telomeres, often devoid of essential genes, is lost due to unstable transmission. Supporting this model, no evidence was found for interstitial telomeric repeats sequences at any of the junctions, albeit such sequences are known to be unstable [130], and might have decayed if initially present. Interestingly, some chromosome fusions described in *Arabidopsis* may have originated through a similar process [45].

A derived hypothesis for Model A, particularly considering the shorter centromeres in *Kwoniella* compared to *Cryptococcus*, is the potential loss of centromere function due to telomeric silencing or DNA sequence erosion resulting from pericentric inversion. In this context, a possible outcome following the pericentric inversion could be the restoration of viability through chromosomal fusion. This hypothesis aligns with findings in *C. deuterogattii*, where centromere deletion led to both neocentromere formation (without karyotype change) and chromosome fusion, reducing the karyotype from 14 to 13 chromosomes [64]. Interestingly, in this case no interstitial telomeric sequences were found at the junctions, which points towards microhomology-mediated end joining (MMEJ) as the likely fusion mechanism [64]. Thus, events impairing centromere function may have driven chromosome fusion in *Kwoniella*.

Model B, on the other hand, proposes an “end-to-end fusion” hypothesis, possibly in cells with dysfunctional or critically shortened telomeres, engaging in nonhomologous end-joining. This process potentially results in an unstable dicentric chromosome, which may be stabilized into a monocentric chromosome through an inversion targeting one of the centromeres (leading to its inactivation) and the initial fusion point (potentially a fragile site). This contrasts with telomere-telomere style chromosome fusion events observed in other species, such as the formation of chromosome 2 in humans [42], the fusion events leading to the 4-chromosome karyotype in *F. graminearum* [48], and similar events during laboratory crosses in *C. deneoformans* [131].

In both models, pericentric inversions might have been mediated by repetitive sequences like transposable elements in inverted configurations near centromeres and telomere-proximal regions. While no such inverted repeats were detected in our study, possibly due to decay over time, this hypothesis finds a parallel in recent chromosome fusion events in muntjac deer. In these species, telomeric and centromeric repeats at the fusion sites of ancestral chromosomes are still present, indicating their role in driving illegitimate recombination leading to chromosome fusions [41]. Notably, the events in muntjac deer occurred roughly 3 mya, thus significantly more recent than the divergence of the last common ancestor between two *Kwoniella* species with different karyotypes, such as *K. europaea* (1n=3) and *K. bestiolae* (1n=8), estimated ∼21.2 mya. The potential discovery of more recently diverged *Kwoniella* species exhibiting karyotypic differences due to similar chromosomal fusion events could provide insights into these processes.

Moving forward, experimental approaches such as CRISPR-mediated pericentric inversion, could model chromosomal fusion events observed in *Kwoniella* under laboratory conditions. Alternatively, using CRISPR to induce chromosome fusions, generating dicentrics similar to recent experiments in *C. deuterogattii* [63], could test if post-fusion pericentric inversions occur. Successfully reducing chromosome numbers in both budding (*S. cerevisiae*) and fission (*Schizosaccharomyces pombe*) yeasts, yielding functional single-chromosome organisms, was recently achieved [132, 133]. These unique yeasts showed comparable vitality to their wild-type counterparts under various conditions and stresses, with at most a slight reduction. However, when mixed with a normal strain, the single-chromosome budding yeast was rapidly outcompeted [132], suggesting that the mild fitness differences observed in the lab might be more detrimental in natural environments. Additionally, several genes involved in DNA replication were upregulated in the single-chromosome budding yeast, indicating challenges in replicating the giant chromosome [132]. Interestingly, our selection analysis comparing *Kwoniella* species with 3 chromosomes to a sister clade with 8 to 11 chromosomes reveals differential selection in proteins associated with the centromere and kinetochores. This suggests centromere instability might be a catalyst for chromosome fusion in *Kwoniella,* and the potential mitotic segregation and replication challenges associated with larger chromosomes [66] have been either overcome or significantly mitigated in *Kwoniella*. Future research will examine chromosome stability in these species under different conditions, including meiosis.

Other mechanisms for chromosome fusion include dysfunction in telomere protection, as seen in human cells with compromised shelterin complexes leading to dicentric fusions [134-136]. While similar defects in shelterin subunits or telomerase in *Kwoniella* could be promoting chromosome fusions, our analysis did not reveal obvious defects in shelterin subunits or telomerase compared with *Ustilago maydis* [83, 137, 138], although the RNA subunit of telomerase in *Kwoniella* remains unidentified. While challenging to detect bioinformatically, innovative approaches like those used in *U. maydis* [139, 140] could be key. However, telomeric repeat sequences present at chromosome termini in *Kwoniella* suggests functional telomerase and intact telomeres.

In contrast to chromosome fusions leading to giant chromosome formation in *Kwoniella* species, a strikingly different mode of karyotype evolution emerged from our comparative genomic analysis of pathogenic and nonpathogenic *Cryptococcus* species. The ancestral karyotype of 14 chromosomes has remained largely conserved in all of the pathogenic species and most (4 of 6) nonpathogenic species. Despite this conservation, *Cryptococcus* species have experienced significantly more interchromosomal rearrangements, both within and beyond centromeric regions. Intercentromeric recombination between abundant and shared centromeric TEs has been well-documented in *C. neoformans* and *C. amylolentus* [23, 24, 62]. Such recombination leads to balanced chromosomal translocations, resulting in stable monocentric chromosomes rather than unstable dicentric resulting from chromosome fusions [39]. In *C. depauperatus*, the chromosome number has reduced from 14 to 8, by a process different from simple chromosome fusion. Instead, this reduction results from different types of rearrangements, including intercentromeric recombination followed by loss of repeat-rich centromeres. DNA double-stranded breaks in these regions can promote loss of centromeric sequence [39], and in *Malassezia* species centromere fission followed by fusion of the acentric chromosome arms to other chromosomes has driven chromosome number reduction [51].

Why is the pattern of karyotype evolution so strikingly different between *Cryptococcus* and *Kwoniella* species? These differences may be attributable to the size and complexity of their centromeres, as well as the presence or activity of mechanisms constraining TE movement. *Cryptococcus* centromeres are, on average, nearly five times larger than *Kwoniella* centromeres, with the largest *Cryptococcus* centromere exceeding 120 kb, compared to ∼30 kb in *Kwoniella*. This size difference, along with a higher number of shared TEs in *Cryptococcus* centromeres, likely increases the frequency of homologous recombination leading to more chromosomal translocations. Additionally, our analysis also shows significantly higher TE density in *Cryptococcus* compared to *Kwoniella* (over tenfold), suggesting a more active role of TEs in *Cryptococcus* genomic rearrangements. The disparity in TE prevalence between the two genera points to different mechanisms of TE control. One notable difference is the absence of the *de novo* DNA methyltransferase (DnmtX) gene in all *Cryptococcus* species, whereas all *Kwoniella* species have retained this gene. Based on analysis of 5mC DNA methylation patterns across the *Kwoniella* genus, it is clear TEs are methylated, and in species where H3K9me2 was analyzed, this also correlates with heterochromatin formation. Thus, the combination of *de novo* 5mC DNA methylation and heterochromatin formation may operate to dramatically reduce TE activity in *Kwoniella*, resulting in more compact centromeres and a lower genome-wide TE density. In contrast, without DnmtX, *Cryptococcus* species are unable to establish new methylation patterns on recently mobilized TEs, leading to less controlled TE activity and increased density in the genome. It is currently unclear if loss of *de novo* methylation in *Cryptococcus* is compensated by other TE suppression mechanisms, such as histone modifications (H3K9me2) or RNA interference. Future research aimed at deciphering the interplay between DNA methylation and other epigenetic mechanisms in regulating TE activity in these species could involve expressing the DnmtX gene in *C. neoformans* strains, both with and without active RNAi and that exhibit high TE loads [119, 141], and observing if this decreases transposon mobilization.

A final interesting facet that emerged from our comparative genomic analysis was the finding that several *Kwoniella* species harbor mini-chromosomes as linear pieces of DNA with telomeric repeats at both ends. To our knowledge, this is a novel finding for a yeast. Previous studies in other fungi have revealed similar examples of what have been termed accessory, dispensable, or B-chromosomes, and these have been associated with host range of plant fungal pathogens [142-145]. At present, the origin and biological function(s) of the *Kwoniella* mini-chromosomes are unknown and it is unclear if they represent remnants resulting from past chromosome fusion events. Future studies can be directed to test their stability during mitosis, transmission following genetic crosses, and possible functions of genes they encode through gene deletion or chromosome loss analyses.

This research provides a robust platform for further studies. Incorporating comparative transcriptomic data could refine current annotations and aid in functional gene characterization, particularly those relevant to pathogenicity across different species. Future studies may also include comparison of the closely related species comprising the *C. gattii* species complex to define genetic factors associated with more prevalent infections in immunocompromised patients by some species and in immunocompetent patients by others. Assessing the pathogenic potential of species of these two genera is crucial, especially as human encroachment into natural habitats exposes us to new opportunistic pathogens. This knowledge will be vital in anticipating and mitigating future health threats posed by these fungi.

## Materials and methods

### Strains and media

Strains studied in this work were grown on YPD (10 g/L yeast extract, 20 g/L Bacto Peptone, 20 g/L dextrose and 20 g/L agar) media unless specified otherwise. *Cryptococcus* strains were incubated at 30°C while *Kwoniella* strains were grown at room temperature (20-23°C). *E. coli* strains were grown on FB media (25 g/l tryptone, 7.5 g/L yeast extract, 1 g/L glucose, 6 g/L NaCl, 50mM Tris HCl pH 7.6) with added ampicillin (100 μg/ml) and kanamycin (50 μg/ml) at 37°C. Strains studied are listed in **S6 Appendix**.

### Genomic DNA extraction

High-molecular weight (HMW) DNA was prepared with a cetyltrimethylammonium bromide (CTAB) extraction as previously described [39], avoiding vortexing during sample preparation. Where necessary, DNA samples for Oxford Nanopore or PacBio long read sequencing were enriched for HMW DNA (> 25 kb) employing the Short Read Eliminator Kit (Circulomics/PacBio). Quality control was performed by determining A260/A280 and A260/A230 ratios on NanoDrop, and quantification was done with Qubit dsDNA Assay Kit (Invitrogen) on the Qubit fluorometer. The size and integrity of the DNA were confirmed by clamped homogeneous electric fields (CHEF) electrophoresis carried out at 6V/cm with an initial switch time (IST) of 1 sec and final switch time (FST) of 6 sec, for 18 h at 14 °C, in a CHEF-DR III system apparatus (Bio-Rad). CHEF gels were prepared with 1% pulsed field certified agarose (BioRad) in 0.5X TBE or 1X TAE, with CHEF DNA 8-48 kb and CHEF DNA 5 kb (BioRad) size standards. For some samples, gDNA extraction for Illumina sequencing was done with a phenol:chloroform-based protocol previously described [146], with minor modifications. Briefly, equivalent amounts of cell pellet and 0.5 mm acid-washed beads (approximately 250 µL) were mixed and washed with sterile bi-distilled water. After centrifugation and removal of the supernatant, the pellet and beads were resuspended in 500 µL of DNA lysis buffer (10 mM Tris, 1 mM EDTA, 100 mM NaCl, 1% SDS, 2% Triton X-100 in water) and 500 µL of phenol:chloroform:isoamyl alcohol (25:24:1) solution. After cell disruption by bead beating at 4°C, centrifugation and collection of the supernatant, an additional chloroform extraction was performed. Supernatants were then precipitated in 100% ethanol for 1h and gDNA pellets were then collected by centrifugation. After performing pellet clean-up with 70% ethanol, gDNA was dissolved in 10 mM Tris-Cl (pH 8), and treated with RNase A for 30 minutes at 37°C. After a final chloroform extraction, ethanol precipitation and washing, the gDNA pellet was resuspended in 10 mM Tris-Cl (pH 8).

### Illumina, Nanopore, and PacBio sequencing

Whole-genome sequencing was performed with Nanopore, PacBio and Illumina technologies. Nanopore sequencing was carried out both in-house (Duke) and at the Broad Institute Technology Labs. PacBio sequencing was conducted at the Duke University Sequencing and Genomic Technologies (SGT) core, and Illumina sequencing was performed either at the Broad Institute Genomics Platform or at the Duke SGT. For PacBio sequencing, 15-to-20-kb insertion-size libraries were prepared and run on a PacBio RS II or Sequel (2.0 chemistry) system. For nanopore sequencing, a single strain was sequenced using the SQK-LSK108 kit, or up to four different DNA samples were barcoded using the SQK-LSK109 and EXP-NBD103/EXP-NBD104 kits. These libraries, either single or pooled, were sequenced on R9 flow-cells (FLO-MN106) for 48 h or 72 h at default voltage in a MinION system using the latest MinION software. For some strains, two Illumina libraries were constructed. A fragment library was prepared from 100 ng of genomic DNA, sheared to ∼250 bp using a Covaris LE instrument, and adapted for sequencing as previously described [147]. A 2.5-kb ‘jumping’ library was prepared using the 2-to-5-kb insert Illumina Mate-pair library prep kit (V2; Illumina). These libraries were sequenced on an Illumina HiSeq 2000, producing 101-base paired reads. Specific details on sequencing platforms, basecalling, and de-multiplexing are provided in **S1 Appendix** for each genome.

### Genome assembly

Initial assemblies were conducted with Illumina data using Allpaths [148] for preliminary investigations of genome architecture. Complete genomes were then assembled with Canu [149] using default parameters and Nanopore or PacBio data, followed by polishing with Illumina short reads (see **S1 Appendix** for details). The consensus accuracy of Nanopore-based assemblies was improved by first correcting errors with Nanopolish v0.11.2 (https://nanopolish.readthedocs.io/en/latest/), and then with up to five rounds of polishing with Pilon v1.22 [150] *(--fix all*) with Illumina reads mapped to the first pass-polished assembly using BWA-MEM v0.7.17-r1188 [151]. PacBio-based assemblies were only polished with Pilon as above. Contigs containing exclusively rDNA sequences detected by Barrnap (https://github.com/tseemann/barrnap) *(--kingdom euk*) or that could be assigned to mitochondrial DNA, were removed from the final nuclear assemblies. Assembly integrity (including telomeric regions) was confirmed by aligning Canu-corrected and Illumina reads with minimap2 v2.9-r720 [152] and BWA-MEM v0.7.17-93 r1188, respectively, and examining read coverage profiles in the Integrative Genomics Viewer (IGV) [153]. Genome assemblies and sequencing data are available at DDBJ/EMBL/GenBank, with accession numbers given in **S1 Appendix**.

### Gene prediction, annotation, and statistical analyses

Gene models were predicted *ab initio* with BRAKER2 v2.1.5 [154] as previously described [11]. BRAKER2 was run in ETP-mode when RNA-seq data was available, leveraging both RNA-seq and protein data for GeneMark training. Otherwise, BRAKER2 was run in EP-mode, relying only on protein data for training. Protein sets from *C. neoformans* H99 [24] and *C. amylolentus* CBS6039 [23], along with RNA-seq data from two growth conditions (see below), were used as input. Naming of protein-coding genes combined results from HMMER PFAM/TIGRFAM, Swiss-Prot, and KEGG products. Gene set completeness was assessed with BUSCO v4.0.6 against the tremellomycetes_odb10 database [155, 156]. Genomic features (number of genes, the number of introns in coding sequences (CDSs), and mean intron length) were calculated using AGAT (https://github.com/NBISweden/AGAT), utilizing the “agat_sp_statistics.pl” tool. Statistical analyses comparing these genomic features between *Cryptococcus* and *Kwoniella*, employed Python3 with Pandas, Seaborn, Matplotlib, and SciPy libraries. Differences between the two groups were assessed using the two-sided Mann-Whitney U test. To evaluate correlations within each group, linear regression analysis was conducted between genome size and other genomic metrics. The strength of these correlations was quantified using the coefficient of determination (R²), and their significance was determined using the t-test for the slope of the regression line.

### RNA extraction, sequencing, and data processing

RNA-seq libraries were prepared from *C. bacillisporus* CA1280, *C. decagattii* 7685027, *C. gattii* WM276, and *C. tetragattii* IND107 cells grown in 50 ml YPD, at 30°C or 37°C, conducted in duplicate. Each of these *Cryptococcus* cell preparations was spiked in with one-tenth (OD/OD) of *S. cerevisiae* strain S288C cells grown in YPD at 30°C, followed by washing and snap freezing. Total RNA was extracted with TRIzol as previously described [24] and then adapted for sequencing using the TagSeq protocol [157], in which ribosomal RNA was depleted using the RiboZero Yeast reagent. Sequencing was performed on an Illumina HiSeq 2000 at the Broad Institute Genomics Platform, producing 101-base paired reads. TagSeq adapters were removed and RNA-seq data was preprocessed with Trim Galore v0.6.7 (https://github.com/FelixKrueger/TrimGalore) discarding reads shorter than 75 nt after quality or adapter trimming (parameters: *--paired --quality 20 --phred33 --length 75*). Splice read alignment was performed with STAR aligner v2.7.4a [158] (indexing: *--genomeSAindexNbases 11*; aligning: *--alignIntronMin 10 --alignIntronMax 2000 --outSAMtype BAM SortedByCoordinate*). Spike-in reads from *S. cerevisiae* were first removed by keeping the reads that did not align to *S. cerevisiae* S288C genome. The remaining reads were then mapped to the respective *Cryptococcus* genome assemblies, and the resulting BAM files were input into the BRAKER pipeline.

### Whole-genome bisulfite sequencing (WGBS) and 5mCG analysis

Genomic DNA from *Kwoniella* strains CBS8507, CBS10737, CBS12478, CBS6074, CBS10118, and PYCC6329 was isolated following the CTAB method. Following quantification and quality control, whole-genome bisulfite sequencing was performed at the Duke University SGT on a NovaSeq 6000 system to generate 50-base paired-end reads. Bisulfite-treated library reads were trimmed with Trim Galore v0.6.7 and analyzed with Bismark v0.22.3 [159] employing bowtie2 and the respective reference genome. Methylation was called using Bismark default settings. Additionally, 5mCG calls were obtained from the Nanopore data using Nanopolish [160]. Output files were converted to bedGraph format for visualization in IGV or for plotting with pyGenomeTracks [161].

### Ortholog identification, alignment and selection analysis

A phylogenomic data matrix was constructed with single-copy orthologs determined by OrthoFinder v2.5.2 [162] across all *Cryptococcus* and *Kwoniella* species, and three outgroups: *Tremella mesenterica* ATCC28783 (GCA_004117975.1), *Saitozyma podzolica* DSM27192 (GCA_003942215.1), and *Bullera alba*JCM2954 (GCA_001600095.1). OrthoFinder was run with default setting and with BLAST as the search tool. The amino acid sequences of 3,430 single-copy orthologs, identified as shared among all species, were individually aligned with MAFFT v7.310 [163] (arguments: *--localpair --maxiterate 1000*) and, subsequently trimmed with TrimAl v1.4.rev22 [164] (parameters: *-gappyout -keepheader*). For selection analysis, nucleotide sequences of single-copy orthologs were backaligned to their amino acid alignments using Egglib v3 [165]. For branch model analysis, a foreground species clade with three chromosomes (*K. europaea* PYCC6329, *Kwoniella* sp. B9012, *K. botswanensis* CBS12716, and *K. mangrovensis* CBS8507) was compared to a background species clade with 8 to 11 chromosomes (*K. bestiolae* CBS10118, *K. dejecticola* CBS10117, *K. pini* CBS10737, and *K. dendrophila* CBS6074), using an un-rooted tree with codeml (with model=2 NSsites=0 CodonFreq=7 and estFreq=0) [166-168]. A null M0 model was run on the un-rooted tree with no clade labels. Significance of fit between these models measured by the likelihood ratio test was corrected for multiple testing using FDR correction.

### Species phylogeny and time tree estimation

A Maximum likelihood (ML) phylogeny was with IQ-TREE v2.1.3 [169], employing a concatenation approach with gene-based partitioning. Individual protein alignments were input to IQ-TREE with *"-p"* argument to form a supermatrix (of 32 taxa, with 3,430 partitions and 1,803,061 sites) for partition analysis. This approach employs an edge-linked proportional partition model to account for variances in evolutionary rates across different partitions. The best amino acid substitution model for each partition was identified by ModelFinder applying the Bayesian information criterion (BIC). The highest-scoring ML tree was determined using the parameters “*--seed 54321 -m MFP -msub nuclear -B 1000 -alrt 1000 -T 20*”, incorporating 1,000 iterations of the Shimodaira–Hasegawa approximate likelihood ratio test (SH-aLRT) and ultrafast bootstrap (UFboot) for branch support.

A time tree was computed in MEGA11 [170] utilizing the RelTime method [171]. This method relaxes the strict molecular clock assumptions in phylogenetic analysis and converts relative node ages into absolute dates using calibration constraints on one or more nodes. The inferred ML species phylogeny was taken as input and transformed into an ultrametric tree with relative times. Absolute dates were then assigned by applying three calibration constraints obtained from other studies [55, 56] and the TimeTree project [172] (http://www.timetree.org/): the separation of *T. mesenterica* from other species (153.0 mya), the emergence of the pathogenic *Cryptococcus* species (27.0 mya) and the split between *C. neoformans* and *C. deneoformans* (24 mya). In RelTime, divergence times for the outgroup are not estimated when applying calibration constraints, as it relies on ingroup evolutionary rate to estimate divergence times, without presuming the evolutionary rates in the ingroup clade are applicable to the outgroup. Therefore, to utilize the divergence of *T. mesenterica* from other species as a calibration point, this species was incorporated as part of the ingroup for this analysis.

### Gene genealogies

Protein sequences for the genes of interest were retrieved from the relevant orthogroups identified by OrthoFinder, subjected to manual inspection, and reannotated as required. The curated protein sequences were subsequently aligned, trimmed, and ML phylogenies were generated with IQ-TREE2. The resulting phylogenies were visualized with iTOL v5.6.3 [173]. The specific model parameters for phylogenetic reconstruction are provided in the corresponding figure legends.

### Synteny analyses

Conserved synteny blocks between pairwise comparisons of *Cryptococcus* and *Kwoniella* genomes were determined using SynChro [65] with synteny block stringency (delta parameter) set to 3. Comparisons within *Kwoniella* and *Cryptococcus* respectively employed the genomes of *K. shandongensis* and *C. neoformans* as references. Synteny blocks determined by SynChro were also input to MCScanX_h [174] and visualized with SynVisio (https://synvisio.github.io/#/) for representation purposes (**Figs 1C** and **6A**, and **S3A Fig**). Detailed linear synteny plots comparing chromosomes and specific genomic regions, such as centromeres, were generated with EasyFig [175] using BLASTN and retaining hits above 200 bp. Fusion events within *Kwoniella* were defined based on synteny plots and the positional information of centromeres in chromosomes as determined by *in silico* analysis and experimentally validated for selected species. By leveraging these visual representations, established phylogenetic relationships, and adhering to the principle of parsimony, chromosomal alterations were inferred with the following rationale: changes observed in identical order and orientation in two sister species were presumed to have been present in their common ancestor. Any changes failing to meet this criterion were categorized as lineage-specific alterations. To enhance the clarity and readability of the figures, we modified the color schemes and labels in the SynChro, SynVisio, and EasyFig plots using Adobe Illustrator.

### Analysis of repeat sequences and transposable elements

Repetitive elements were identified independently for all genomes by leveraging widely used library-based and *de novo* TE annotation tools as implemented in the EarlGrey pipeline [176]. Briefly, known repeats are first identified with RepeatMasker (CONS-Dfam_withRBRM_3.7) and *de novo* TE identification is performed with RepeatModeler2. Next, the set of consensus sequences obtained *de novo* are clustered using CD-HIT-EST to reduce redundancy. The resulting TE consensus sequences are extended through an interactive “BLAST, Extract, Extend” (BEE) process and redundancy is removed again with CD-HIT-EST. The resulting TE consensus sequences are classified into specific families or labelled as “unclassified” if not matching a known family. The final non-redundant library, which combines both the known TE library and the TE consensus sequences obtained *de novo*, is utilized to annotate TEs across the genome using RepeatMasker, by applying a conservative threshold (“-cutoff 400’). Lastly, spurious hits less than 100 bp in length are excluded from the TE annotations prior to final quantification. Genome-wide TE density plots were visualized using R with the package ggplot2. Overlaps between centromeres and repetitive elements were assessed with bedtools v2.27.1 [177]. The uncharacterized status of significant TE numbers in some species may be attributable to the progressive erosion of TE sequences by mutational processes, diminishing their recognizability and identifiability. TE data associated with Fig 5 is provided in **S6 Appendix.**

### Generation of mCherry-tagged Cse4 (CEN-A) strains in different *Kwoniella* species

The Cse4/CENP-A gene was identified in *Kwoniella* genomes by TBLASTN, with *C. neoformans* (CNAG_00063) and *C. amylolentus* (L202_03810) Cse4 sequences as queries. A mCherry-tagged Cse4 fusion protein was generated via overlap extension-PCR (OE-PCR). For this, the upstream/promoter region of *CSE4* and the *CSE4*-ORF with its respective downstream/terminator region were amplified from strain CBS10118, and the mCherry ORF was amplified from plasmid pVY50 [23] using primers with specific complementary 5’ ends of 60 bp each (**S7 Appendix**). These fragments were then purified and assembled via OE-PCR into a complete amplicon encoding the N-terminally mCherry-tagged Cse4/CENP-A gene, regulated by its endogenous promoter and terminator regions. The amplicon, further amplified with primers MP253/MP254 (containing ApaI and XhoI restriction sites; **S7 Appendix**), was digested with both restriction enzymes and gel purified. Similarly, plasmid pVY50 was digested with the same enzymes, followed by gel purification, to retrieve a 5,809 bp fragment composed by the backbone of the plasmid encoding, among other elements, a fungal neomycin (NEO) resistant gene. The complete digested amplicon was then cloned into the corresponding sites of pVY50 to generate plasmid pMP01 (**S7 Appendix**). The same approach was employed to generate plasmids pMP02 and pMP03 encoding mCherry-tagged Cse4/CENP-A of *K. europaea* PYCC6329 and *K. dendrophila* CBS6074, respectively (**S7 Appendix**). The resulting plasmids were introduced into TOP10 *E. coli*, grown overnight at 37°C, and post-miniprep recovery, validated through restriction analysis and Sanger sequencing.

A minimum of 10 µg of circular plasmid DNA was used for biolistic transformation [178] of each *Kwoniella* strain. Transformants were selected on YPD media supplemented with 200 μg/ml of neomycin. Transformation of *K. bestiolae* and *K. europaea* successfully yielded transformants but attempts to transform *K. dendrophila* CBS6074 with plasmid pMP03 were unsuccessful. Consequently, *K. pini* CBS10737, phylogenetically close to *K. dendrophila* and with 11 chromosomes, was chosen as an alternative. Biolistic transformation of *K. pini* with pMP03 successfully recovered transformants. At least two independent transformants of each species underwent fluorescent microscopy screening to ascertain the expression of the tagged protein. Selected transformants were stained with Hoechst 33342 and imaged using a Delta Vision Elite deconvolution microscope with a CoolSNAP HQ2 CCD camera at Duke University Light Microscopy Core Facility. Images were processed using Fiji-ImageJ (https://imagej.net/Fiji) (RRID:SCR_002285).

### Chromatin immunoprecipitation followed by high throughput sequencing (ChiP-seq)

Chromatin immunoprecipitation followed by sequencing (ChIP-seq) was conducted to identify Cse4 enriched regions in tagged species (*K. bestiolae*_mCherry-Cse4, *K. europaea*_mCherry-Cse4, and *K. pini*_mCherry-Cse4), using a polyclonal antibody against mCherry (ab183628, Abcam) as previously described [64]. Similarly, ChIP-seq for *K. bestiolae* CBS10118, *K. europaea* PYCC6329, *K. pini* CBS10737, *K mangrovensis* CBS8507, and *K. dendrophila* CBS6074 was performed to detect histone H3K9me2 enriched regions, employing a monoclonal antibody against histone H3K9me2 (ab1220, Abcam). Libraries were prepared and sequenced at the Duke University SGT, using either a NovaSeq 6000 or a HiSeq 4000 instrument to produce 50-bp paired-end reads. ChIP-seq sequencing reads were trimmed with Trim Galore v0.6.7, and subsequently aligned to each respective genome assembly with bowtie2. Read duplicates were removed with Picard and SAMtools, and bamCompare v3.5.4 was used to normalize the ChIP-seq data against the input control. The resulting files were converted to bedGraph format for visualization in IGV or for plotting with pyGenomeTracks.

### CHEF electrophoresis of *Kwoniella* chromosomes

Spheroplasts with intact chromosomal DNA were generated in plugs, as previously described [179], for *Kwoniella* strains CBS10118, CBS8507 and PYCC6329 with minor modifications: i) cells were grown in Yeast Nitrogen Base (YNB) liquid minimal medium supplemented with NaCl (1 M), and ii) zymolase (25 mg/mL), or *Trichoderma harzianum* lysing enzymes (50 mg/mL for *K. mangrovensis*) were used to lyse the cells embedded in the agarose, with overnight reactions at 37°C. Gels were prepared with 0.8 or 0.9% of Megabase agarose (BioRad) and chromosomal separation was conducted with the CHEF DR-II System and the CHEF Mapper XA System (Bio-Rad, Richmond CA) using different running and buffer conditions, and different size markers selected based on the chromosome size range of each assay (details in **S2 Fig**). Following electrophoresis, gels were stained with ethidium bromide, unstained using the running buffer, and photographed under a UV transillumination imaging system.

### Hi-C mapping

Hi-C mapping of *Kwoniella* strains CBS10118, CBS8507 and PYCC6329 was performed as previously described [180], with minor modifications. Cells were grown overnight in YPD, after which a solution of 37% formaldehyde was added to a final concentration of 3% of the total volume. After incubation at 25°C for 20 minutes, the cross-linking reaction was quenched by adding 2.5 M glycine at 2X the volume of formaldehyde used, followed by incubation at 25°C for 20 minutes. Washed cell pellets were resuspended in 50 mL of 1x NEBuffer 2, flash frozen in liquid nitrogen, grinded up to a powder, and then resuspended in the same buffer to an OD600 of 10.0. In situ Hi-C sequencing libraries [181] were generated from 0.75 mL cell suspensions. Cell pellets were resuspended in 250 µL ice-cold Hi-C lysis buffer (10 mM Tris-HCl pH 8.0, 10 mM NaCl, 0.2% Igepal CA630, 1 tablet/10 mL Roche complete mini EDTA-free protease inhibitor), kept on ice for 5 minutes, centrifuged (5 min at 2,500 x g at 4°C) and washed in 500 µL Hi-C lysis buffer. Pellets were resuspended in 50 µL 0.5% SDS and incubated for 10 minutes at 62°C to permeabilize nuclei. After adding 140 µL H_2_O and 25 µL 10% Triton X-100 and 15 minutes at 37°C to quench the SDS, 25 µL 10x CutSmart buffer and 10 µl 10 U/µL MseI restriction enzyme (New England Biolabs) were added, and chromatin digested overnight at 37°C with rotation. MseI was heat-inactivated (20 minutes at 62°C) and fragment ends filled-in and biotinylated by adding 27.5 µl of a cocktail containing 15 µL 1 mM biotin-14-dUTP (Jena BioScience), dCTP, dGTP, dATP (3x 1.5 µL of 10 mM solutions) and 8 µL of 5 U/µl Klenow fragment of DNA polymerase I (New England Biolabs) and a 30-minute incubation at 37°C with rotation. For blunt-end ligation, 900 µL of a cocktail containing 663 µL H_2_O, 100 µL 10% Triton X-100, 120 µL 10x T4 DNA ligase buffer and 5 µL of 400 U/µL T4 DNA ligase (New England Biolabs) was added. After 2 hours at room temperature with rotation, tubes were centrifuged (5 minutes at 2,500 x g), supernatants were carefully removed and the pellets resuspended in 300 µL 1% SDS, 10 mM Tris-HCl pH 8.0, 0.5 M NaCl. After adding 10 µL of 20 mg/µL proteinase K (New England Biolabs), proteins were degraded for 30 minutes at 55°C followed by an overnight incubation at 68°C with shaking to reverse crosslinks. Insoluble material including apparently intact cells were spun down (5 min at 2,500 x g). DNA in the supernatant was precipitated in the presence of 1 ul (20 mg) glycogen with 2 volumes of ethanol at -80°C for 15 minutes and spun down for 15 minutes at 13,000 rpm at 4°C. The pellet was washed with 800 µL 70% ethanol and dissolved in 130 µL TE buffer (10 mM Tris-HCl pH 8.0, 0.1 mM EDTA). The DNA was sheared to ∼450 bp in a 130-µL vial on a Covaris S2 instrument set to 7°C, duty cycle 10%, intensity 4, 200 cycles/burst, 2 cycles of 35 seconds. Sheared DNA was cleaned-up with 0.55 volumes of AmPure XP beads (Beckman Coulter) and eluted in 300 µL TE buffer. DNA molecules containing biotinylated ligation junctions were pulled down on Dynabeads MyOne Streptavidin T1 beads (Life Technologies). For each capture, 150 µL beads were washed with 400 µL 1x Tween Washing Buffer (TWB; 1M NaCl, 5 mM Tris-HCl pH 7.5, 0.5 mM EDTA; 0.05% Tween-20) and resuspended in 300 µL 2x Binding Buffer (2xBB; 2M NaCl, 10 mM Tris-HCl pH 7.5, 1 mM EDTA) and mixed with 300 µL sheared DNA for 15 minutes at room temperature. Beads were washed twice in 600 µL TWB for 2 minutes at 55°C and once in 100 µL TE buffer. On-bead end repair, adapter ligation and PCR amplification was performed using the Kapa Hyper Prep kit (Roche). Beads were resuspended in 60 µL of a cocktail containing 50 µL H_2_O, 7 µL end repair & A-tailing buffer and 3 µL enzyme mix and incubated for 30 minutes at 20°C and 30 minutes at 65°C. A cocktail containing 5 µL H_2_O, 30 µL ligation buffer, 10 µL T4 DNA ligase and 5 µL undiluted (15 µM) Unique Dual Indexed adapters were added and the reactions incubated for 15 minutes at room temperature. Beads were collected, washed twice for 2 minutes at 55°C in TWB and once in 100 µL TE buffer, resuspended in a cocktail containing 40 µL H_2_O, 50 µL 2x Kapa HiFi Hot Start Ready Mix and 10 µL Kapa Illumina amplification primers, split in 2×50 µL in strip tubes and thermocycled: 30 s at 98°C; 8 cycles of 10 s at 98°C, 30 s at 55°C, 30 s at 72°C; 7 minutes at 72°C. Beads were pelleted on a magnet and the supernatant cleaned up with 0.7 volumes of AMPure XP beads. Libraries were characterized by BioAnalyzer, pooled and sequenced on an Illumina HiSeq 2500 instrument. Hi-C plots were generated with Juicebox, using Juicer v1.5.6 after alignment of the Illumina reads with BWA 0.7.12-r1039.

## Acknowledgments

This study was supported by NIH/NIAID R01 grant AI050113-18 and NIH/NIAID R01 grant AI039115-26 awarded to JH, R01 grant AI33654-06 awarded to JH, and Paul Magwene, and NIH/NHGRI grant U54HG003067 and U19AI110818-08 to the Broad Institute. MN acknowledges support from German Research Foundation, DFG grant NO407/7-2. JH is also Co-Director and Fellow of the CIFAR program Fungal Kingdom: Threats & Opportunities. We thank Klaas Schotanus, Vikas Yadav, and Anna Floyd Averette for technical assistance and constructive feedback, Fred Dietrich for computational resources, Chris Todd Hittinger from providing yeast strains, and Duncan Wilson for communication and discussions on the implications of his discovery of retention and loss of the *PRA1/ZRT1* gene cluster in pathogenic fungi. We appreciate Rytas Vilgalys, Teun Boekhout, and Kaustuv Sanyal for their advice, inspiration, and discussions. We thank the Broad Institute Genomics Platform for generating Illumina sequence for this project, the Broad Institute Microbial Omics Core for assistance with RNA sequencing, and Robert Lintner for his assistance with Oxford Nanopore sequencing.

## Author contributions

**Conceptualization:** Marco A. Coelho, Márcia David-Palma, Sheng Sun, Christina A. Cuomo, Joseph Heitman

**Data Curation:** Marco A. Coelho, Minou Nowrousian, Christina A. Cuomo

**Formal analysis:** Marco A. Coelho, Terrance Shea, Katharine Bowers, Sage McGinley-Smith, Minou Nowrousian, Christina A. Cuomo

**Funding acquisition:** Christina A. Cuomo, Joseph Heitman

**Investigation:** Marco A. Coelho, Márcia David-Palma, Sheng Sun, Arman W. Mohammad, Andreas Gnirke

**Project Administration:** Sheng Sun, Christina A. Cuomo, Joseph Heitman

**Resources:** Andrey M. Yurkov, Christina A. Cuomo, Joseph Heitman

**Supervision:** Christina A. Cuomo, Joseph Heitman

**Visualization:** Marco A. Coelho

**Writing – original draft:** Marco A. Coelho, Márcia David-Palma, Joseph Heitman

**Writing – review & editing:** Marco A. Coelho, Márcia David-Palma, Sheng Sun, Christina A. Cuomo, Joseph Heitman

