## Supplementary Information for "Comparative genomics of *Cryptococcus* and *Kwoniella* reveals pathogenesis evolution and contrasting karyotype dynamics via intercentromeric recombination or chromosome fusion"

**This PDF file includes:**

S1 to S18 Figs
Legends for S1 and S2 Text
Legends for S1 to S7 Appendix

**Other supplementary materials for this manuscript include the following:**

S1 and S2 Text
S1 to S7 Appendix

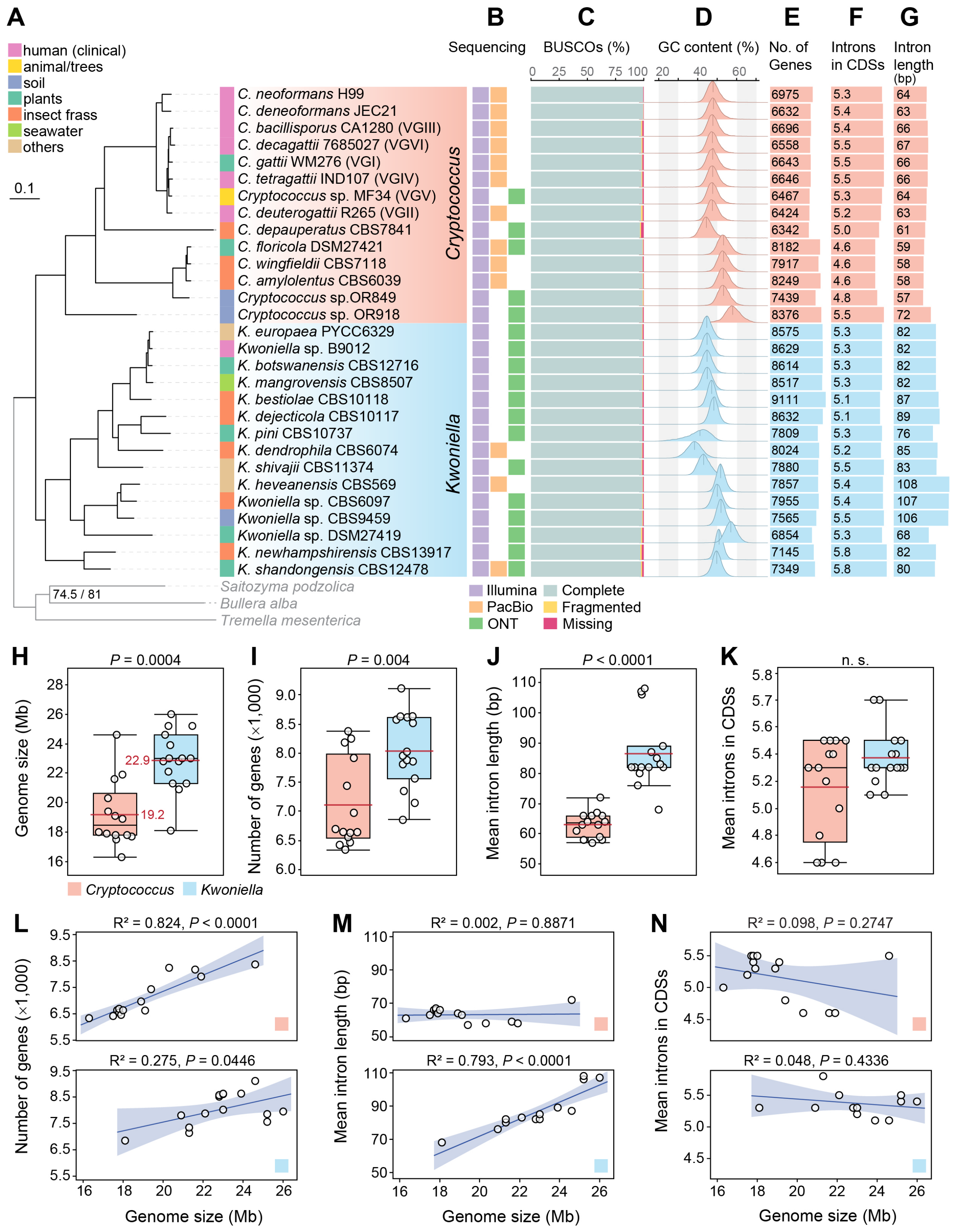


**S1 Fig. *Cryptococcus* and *Kwoniella* phylogeny and genomic features. (A)** Maximum likelihood phylogeny of *Cryptococcus* and *Kwoniella* inferred through a concatenation-based approach on a data matrix composed of protein alignments of 3,430 single-copy genes shared across all species and three outgroups (depicted in grey). Except where indicated, all branches are 100% supported (SH-aLRT and UFboot tests). Branch lengths are given in number of substitutions per site (scale bar). The isolation origin of each strain is indicated as given in the key. **(B)** Genome sequencing approach for each of the strains. **(C)** BUSCO completeness assessment of each genome gene set. **(D)** Frequency distribution of GC content across species, with mean GC values represented by vertical lines. **(E)** Number of genes, **(F)** mean number of introns within coding sequences (CDSs) and **(G)** their mean length (in base pairs, bp). Box-plot comparisons of **(H)** genome sizes, **(I)** number of genes, **(J)** mean intron length, and **(K)** mean number of introns in CDSs, between *Cryptococcus* and *Kwoniella*. The red line, black line, and boxes denote the mean value, median value, and interquartile range, respectively. (*P*-values obtained by Mann-Whitney U test). **(L-M)** Comparative analysis of gene count, intron length, and mean number of introns relative to total genome size. Each plot shows linear regression with confidence values, $R^2$ correlation coefficients values, and *P* values from the *t*-test for the slope of the regression line.

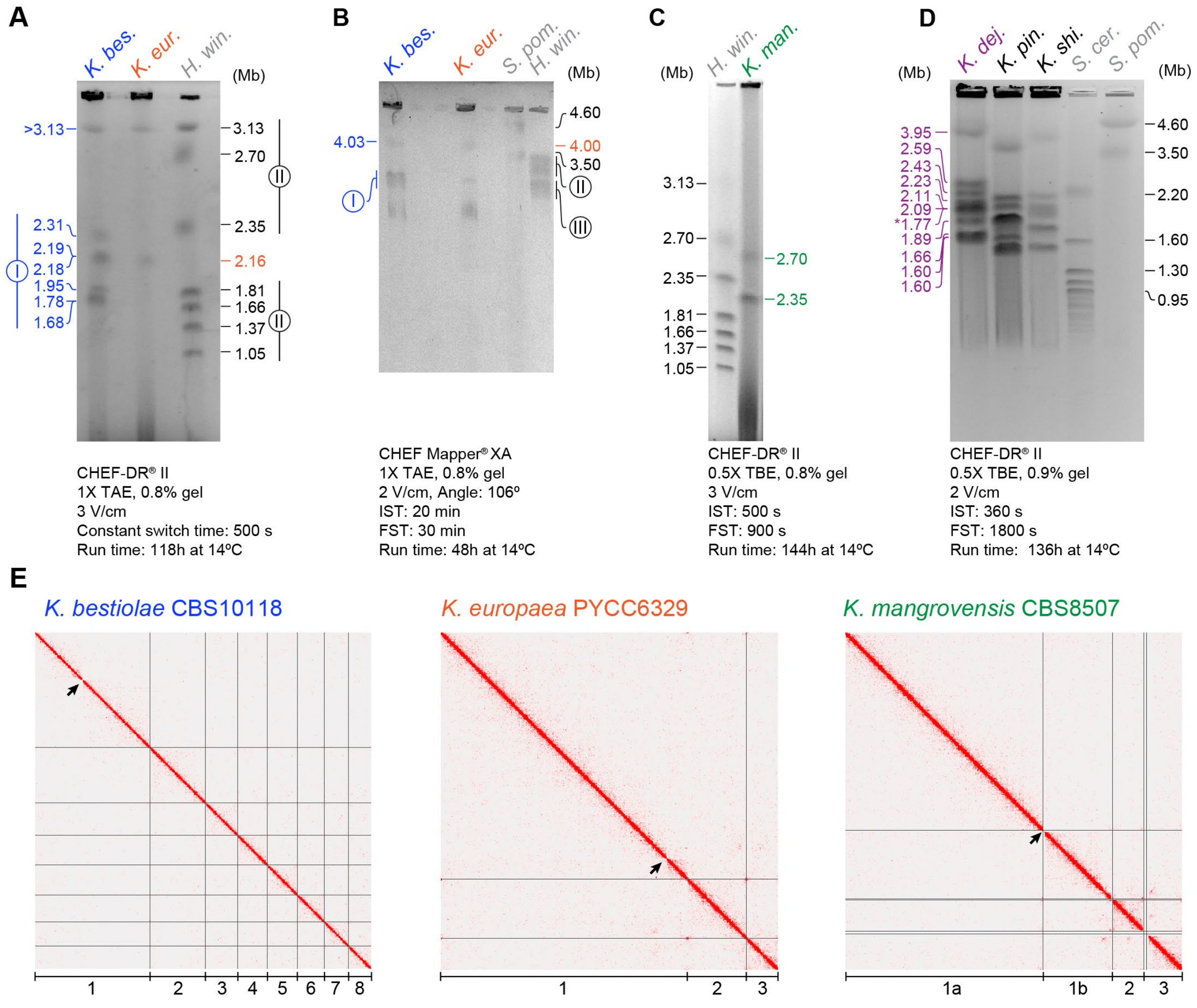


**S2 Fig. Genome assembly validation of representative *Kwoniella* species by clamped homogeneous electrical field (CHEF) electrophoresis and Hi-C mapping. (A-D)** Electrophoretic karyotypes of selected *Kwoniella* species with different number of chromosomes: *K. bestiolae* with 8 chrs (blue); *K. europaea* and *K. mangrovensis* with 3 chrs. each (orange and green, respectively); and *K. dejecticola* (purple), *K. pini* and *K. shivajii* with 11 chrs. each. *Saccharomyces cerevisiae*, *Schizosaccharomyces pombe*, and *Hansenula wingei* chromosomes serve as markers, with their sizes in megabase pairs (Mb) shown in black. Color-coded numbers indicate contigs sizes in each respective assembly. Two running conditions were used for better separation of small and large chromosomes in *K. bestiolae* and *K. europaea* (panels A and B). The largest chromosomes in *K. bestiolae* (8.41 Mb), *K. europaea* (16.67 Mb) and *K. mangrovensis* (18.17 Mb) are too large to be resolved by this approach. The karyotypes of *K. dejecticola*, *K. pini* and *K. shivajii* confirm 11 chrs. in each species, aligning with the contig number and length. The contig size harboring the rDNA array in *K. dejecticola* (indicated by an asterisk) is likely underestimated. **(E)** Hi-C contact matrix showing interaction frequencies between genomic regions, with pixel intensity indicating how often a pair of loci interact. Most of the links are nearby intrachromosomal, validating our assemblies. Interaction frequencies produced with Juicer Tools v1.7.6 are summarized along the genome. Chromosome numbers are given at the bottom of each plot. In *K. mangrovensis*, chr. 1 is broken at the rDNA array (black arrow in each plot) and shown as two contigs (1a and 1b).

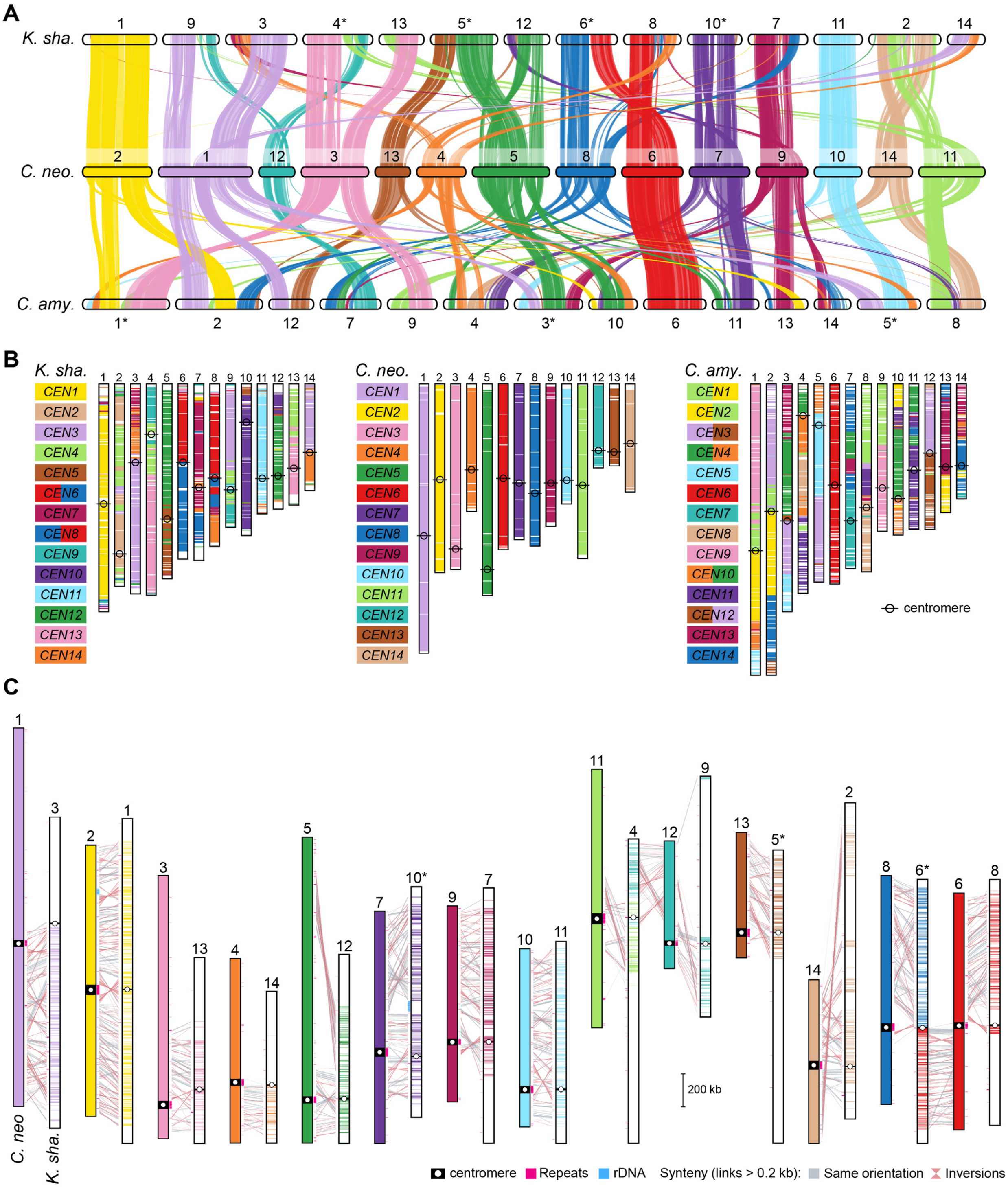


**S3 Fig. Centromere conservation between *C. neoformans*, *C. amylolentus* and *K. shandongensis*.** **(A)** Pairwise synteny relationships between *C. neoformans*, *C. amylolentus* and *K. shandongensis*, all with 14 chromosomes. Links depict boundaries of syntenic gene blocks identified by MCScanX, with pairwise homologous relationships determined by SynChro. Chromosomes are color-coded based on *C. neoformans* and were reordered or inverted (marked with asterisks) from their original assembly orientations to maximize collinearity. **(B)** Superimposition of synteny blocks and centromere locations reveals three intercentromeric rearrangements between *C. neoformans* and *C. amylolentus*, as opposed to a single one between *C. neoformans* and *K. shandongensis*. **(C)** Synteny analysis based on BLASTN comparing chromosomal regions encompassing centromeres in *K. shandongensis* (predicted *in silico*) relative to previously determined centromeres of *C. neoformans*. Despite the numerous intrachromosomal rearrangements between these two species, centromere-flanking regions exhibit full (e.g. *CnCEN2* ) or at least partial (e.g. *CnCEN6/8*) synteny.

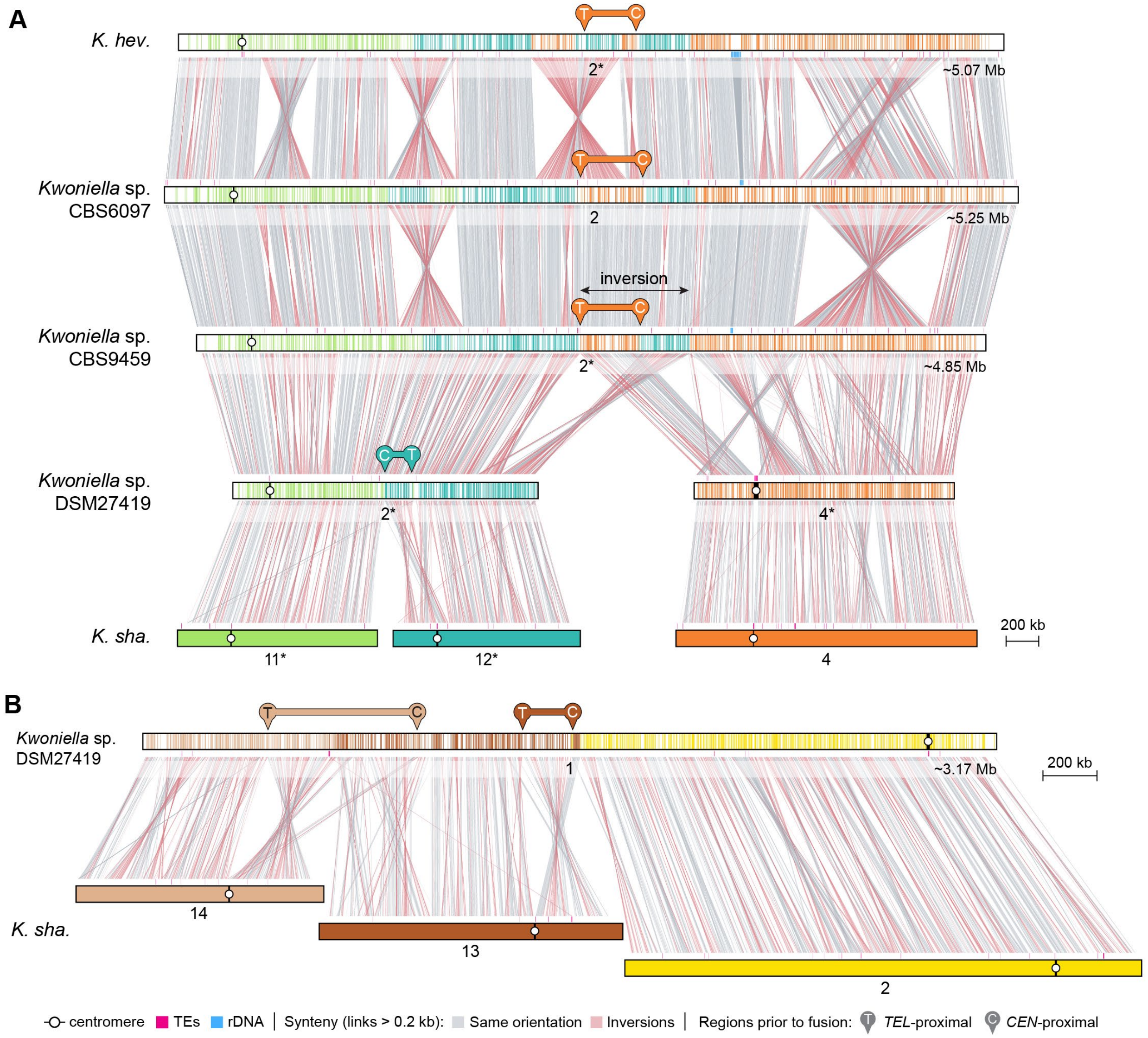


**S4 Fig. Ancestral chromosome fusion events in *Kwoniella*.** Synteny comparison showing that **(A)** chr. 2 of *Kwoniella* sp. DSM27419 resulted from the fusion of *K. shandongensis* chrs. 11 and 12. and that **(B)** chr. 1 of *Kwoniella* sp. DSM27419 emerged from the fusion of three chromosomes extant in *K. shandongensis* (chrs. 2, 13 and 14), followed by several intrachromosomal rearrangements. These two fusion events, inferred as the oldest within *Kwoniella* (event A in Fig 2), resulted in chromosome arrangements consistent across all species after the split from *K. shandongensis*/*K. newhampshirensis* (albeit with a few subsequent species-specific rearrangements). Note that the centromere-proximal regions of *K. shandongensis* chromosomes align at or near the fusion points on *Kwoniella* sp. DSM27419 fused chromosomes, while telomere-proximal regions are more internal, suggesting large inversions targeting centromeric regions accompanied each fusion event. In *K. heveanensis*, and sibling species *Kwoniella* sp. CBS6097 and *Kwoniella* sp. CBS9495, chr. 2 results from a subsequent fusion of *K. shandongensis* chr. 4 to the already fused chr. 11-12 (event D in Fig. 2), followed by several intrachromosomal rearrangements. This event occurred in the common ancestor of these three species and the centromere- and telomere-proximal regions have been inverted back by a secondary inversion (double-sided black arrow) that occurred after the initial fusion. Chromosomes inverted from their original assembly orientations are marked with asterisks.

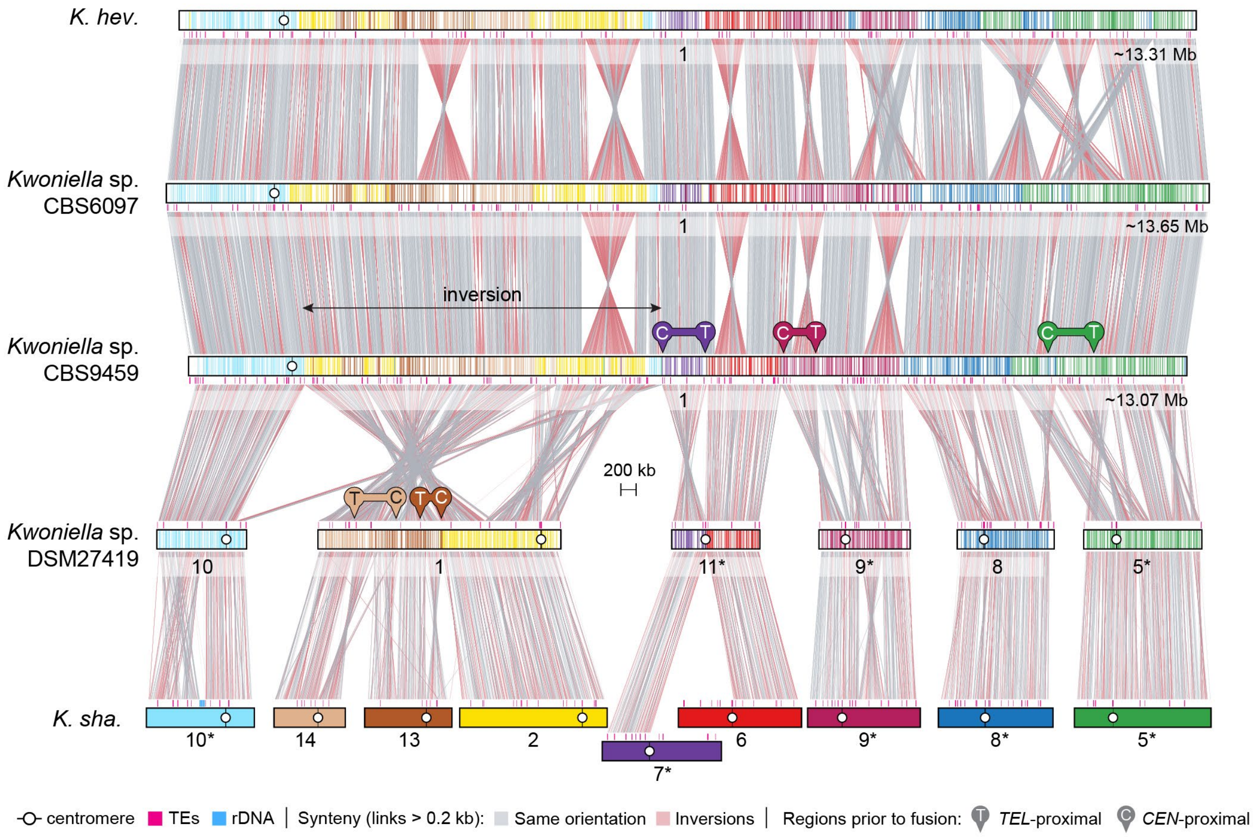


**S5 Fig. Formation of a giant chromosome in *K. heveanensis* and sibling species.** Synteny comparison showing that chr. 1 of *K. heveanensis* and sibling species *Kwoniella* sp. CBS6097 and *Kwoniella* sp. CBS9495 resulted from fusion of 6 chromosomes, followed by several intrachromosomal rearrangements. Three of the ancestral chromosomes had been fused prior to this event (chrs. 2-13-14). Note that most of centromere-proximal regions of *K. shandongensis* chromosomes are located at or near the fusion points on the giant chromosome, whereas the telomere-proximal regions are more internalized, suggesting that a large inversion targeting the centromeric region is associated with each fusion event. Chr. 11 of *Kwoniella* sp. DSM27419 resulted from reciprocal translocation between *K. shandongensis* chrs. 6 and 7 (event C in Fig. 2).

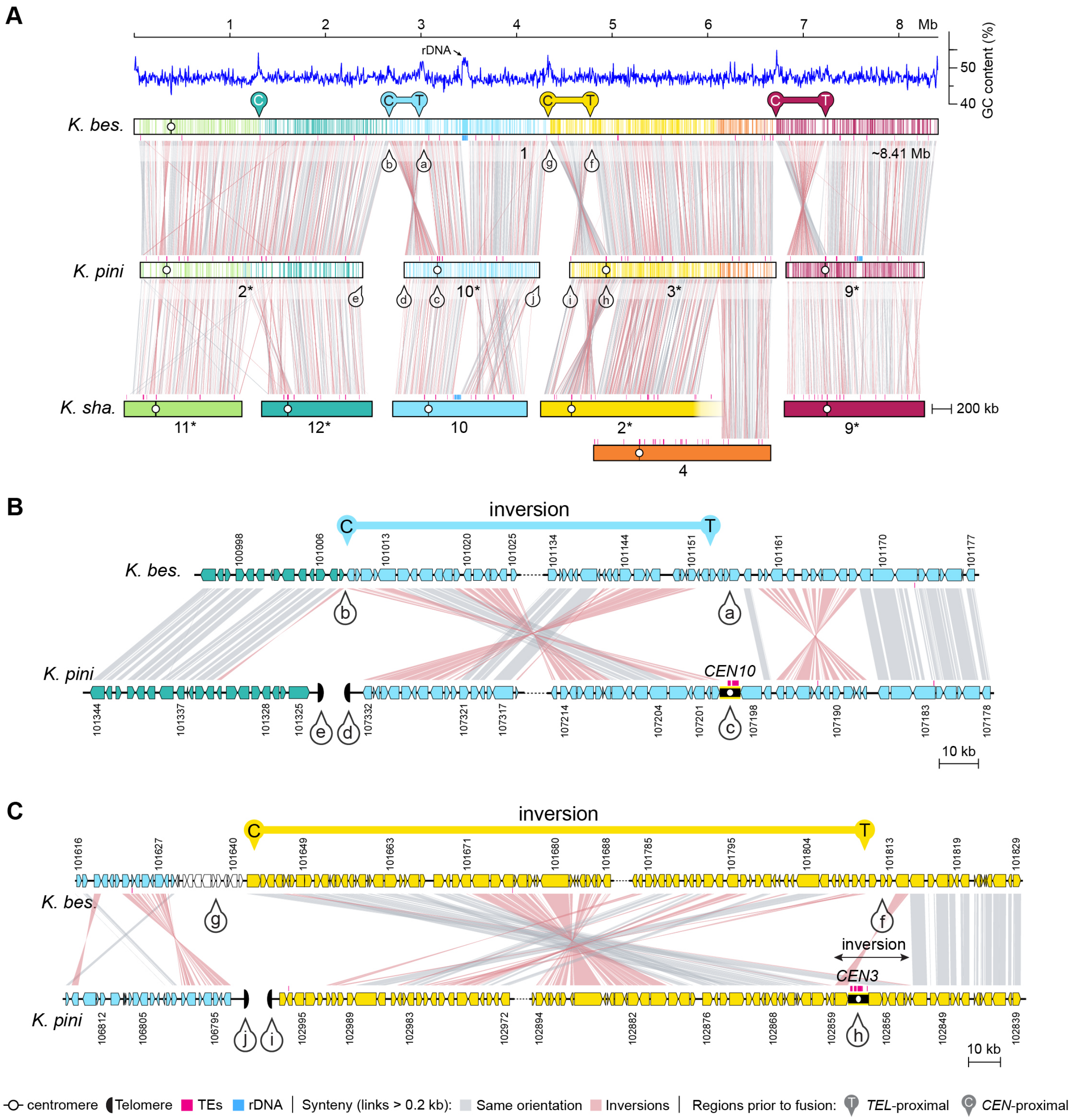


**S6 Fig. Formation of a large chromosome in *K. bestiolae*. (A)** Synteny comparison showing that *K. bestiolae* chr. 1 resulted from fusion of 4 chromosomes, extant in *K. pini*. Two of the ancestral chromosomes had been fused prior to this event (chrs. 11 and 12). *K. pini* chr. 3 resulted from a translocation between *K. shandongensis* chr. 4 and an ancestrally formed chromosome resulting from fusion of *K. shandongensis* chrs. 2, 13 and 14 (event B in Fig. 2). **(B-C)** Zoomed-in synteny views of the regions marked in panel A (pins with lowercase letters from a – h). Note that most of centromere-proximal regions of *K. pini* chromosomes are located at or near the fusion points on the giant chromosome, whereas the telomere-proximal regions are more internalized, suggesting that a large inversion targeting the centromeric region is associated with each fusion event. A secondary inversion (double-sided black arrow) likely occurred in *K. pini* reversing the relative orientation of the centromere and a few flanking genes. Species-specific differences in gene content and relative orientation are expected as these species have diverged for a long time.

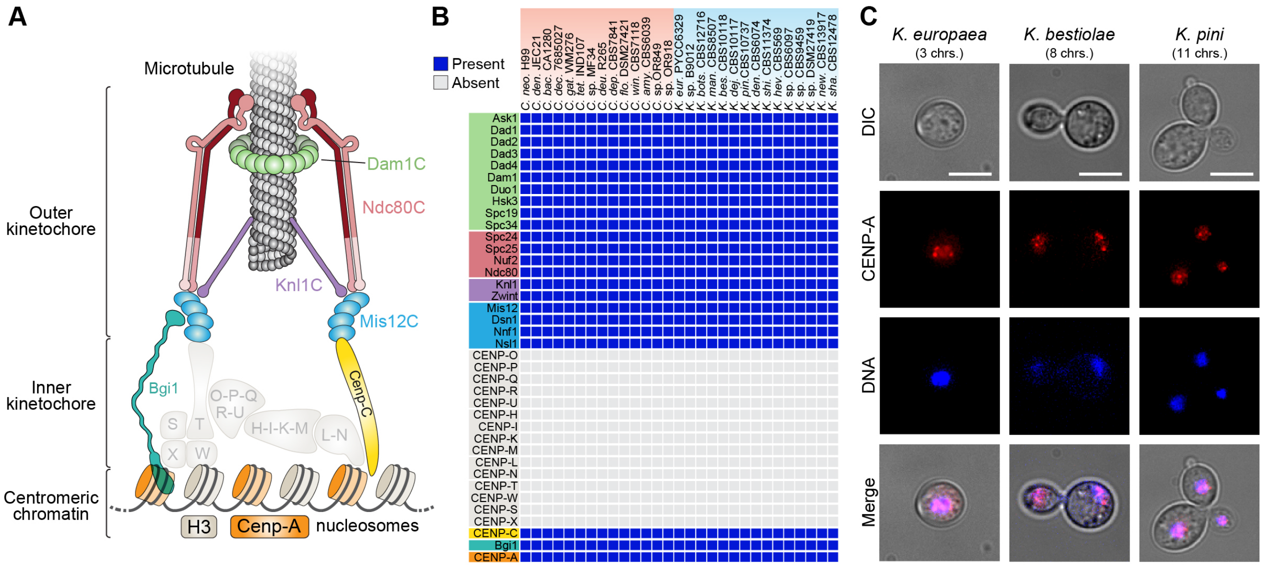


**S7 Fig. CENP-A expression and kinetochore components across *Cryptococcus* and *Kwoniella*. (A)** kinetochore ensemble schematic with color-coded protein complexes or faded out to indicate absence. **(B)** Matrix showing the presence (blue) and absence (grey) of the kinetochore components proteins depicted in panel A. **(C)** Live cell imaging of *K. europaea*, *K. bestiolae* and *K. pini* expressing mCherry-tagged CENP-A, showing nucleus subcellular localization (scale bar, 10 μm).

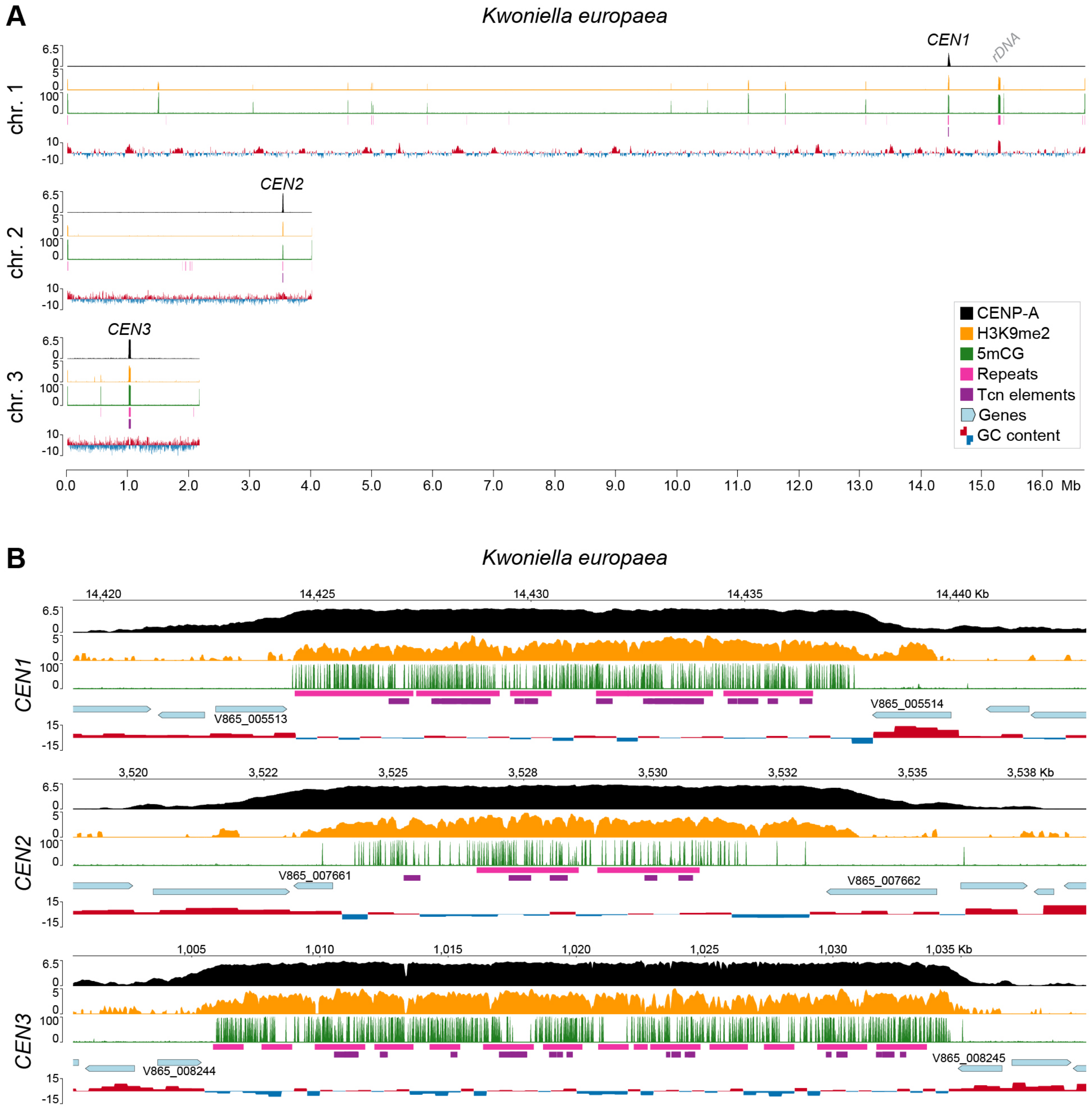


**S8 Fig. Localization of CENP-A to contiguous regions defines centromeres on each of the 3 chromosomes of *K. europaea*. (A)** Whole chromosome plots showing CENP-A (black) and H3K9me2 (orange) enrichment, CG cytosine DNA methylation (5mCG, green) derived from WGBS, repeat content (pink), TCN-like LTR elements (purple) and GC content show as deviation from the genome average (red, above; blue, below). CENP-A and H3K9me2 enriched regions were normalized to input DNA. The data is computed in 5-kb non-overlapping windows. **(B)** zoomed-in sections show the regions spanning the centromeres and adjacent genes (light blue). Note that centromeres are enriched for CENP-A, H3K9me2, and 5mCG DNA methylation marks and contain repeat elements.

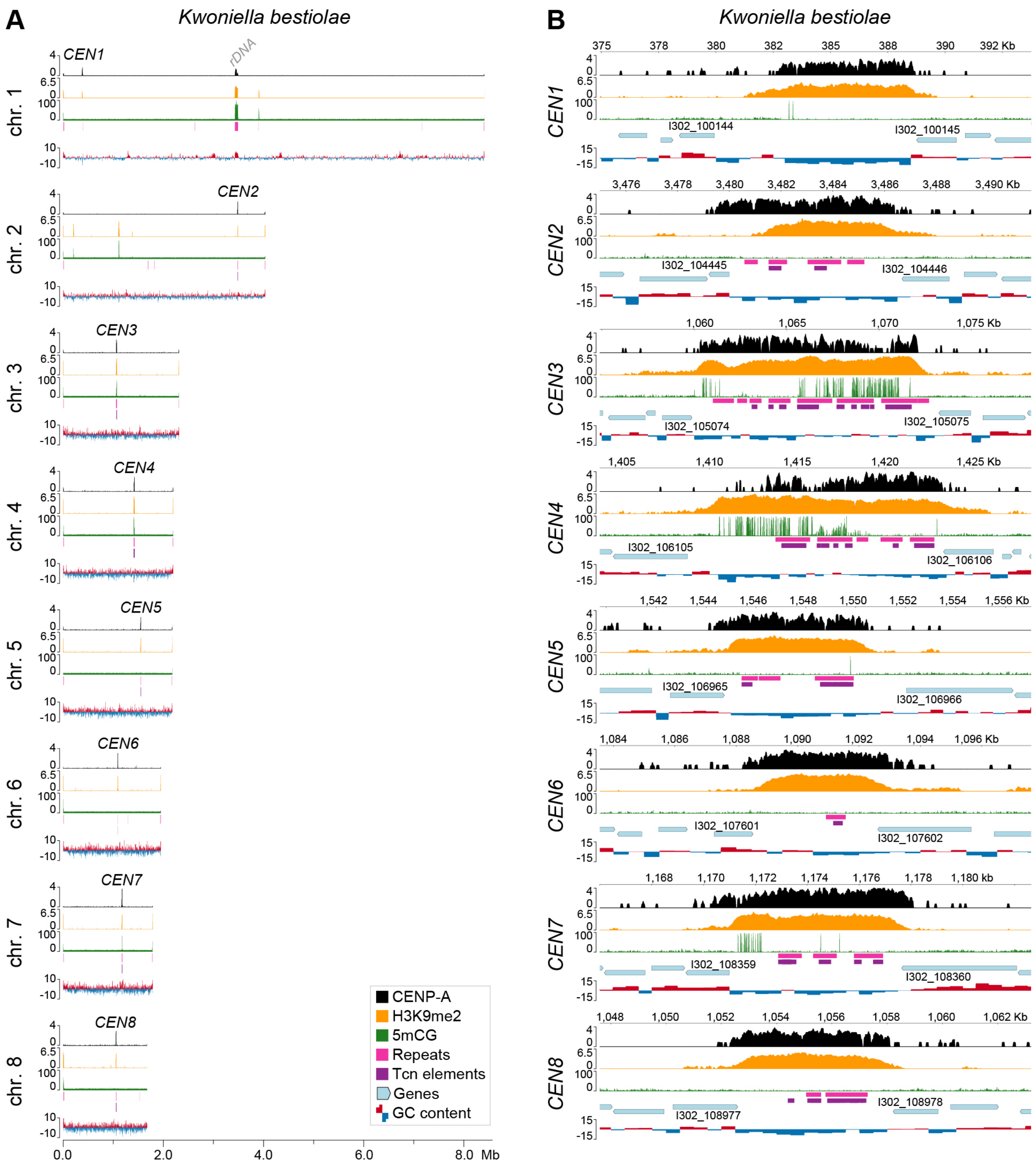


**S9 Fig. Localization of CENP-A to contiguous regions defines centromeres on each of the 8 chromosomes of *K. bestiolae*. (A)** Whole chromosome plots showing CENP-A (black) and H3K9me2 (orange) enrichment, CG cytosine DNA methylation (5mCG, green) derived from WGBS, repeat content (pink), TCN-like LTR elements (purple) and GC content show as deviation from the genome average (red, above; blue, below). CENP-A and H3K9me2 enriched regions were normalized to input DNA. **(B)** Zoomed-in sections show the regions spanning the centromeres and adjacent genes (light blue). Note that centromeres are enriched for both CENP-A and H3K9me2 marks but 5mCG enrichment was only observed in a subset of centromeres, and even in these cases, it was localized to specific regions instead of the whole-centromere. The data is computed in 5-kb non-overlapping windows.

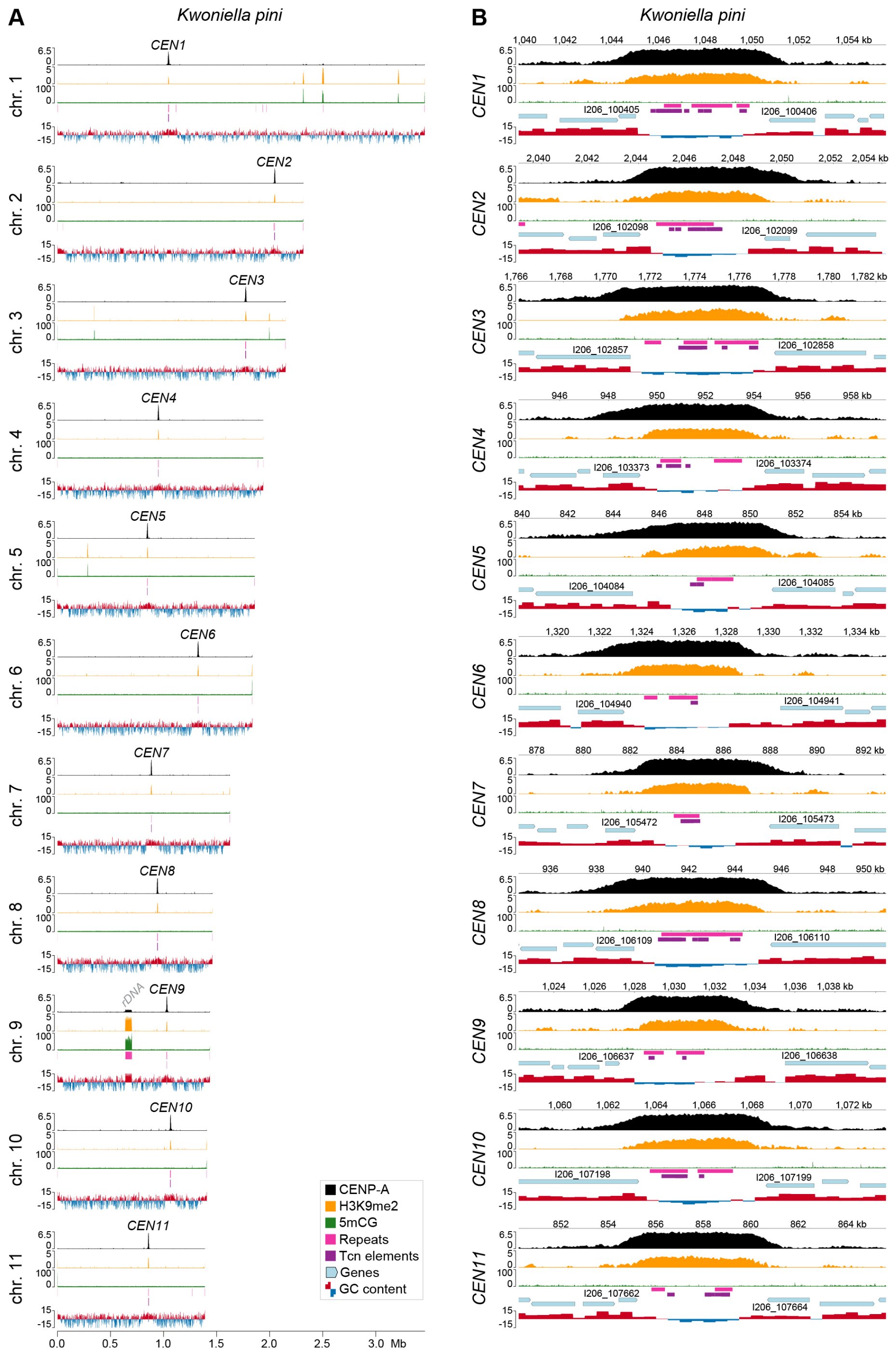


**S10 Fig. Localization of CENP-A to contiguous regions defines centromeres on each of the 11 chromosomes of *K. pini*. (A)** Whole chromosome plots showing CENP-A (black) and H3K9me2 (orange) enrichment, CG cytosine DNA methylation (5mCG, green) derived from WGBS, repeat content (pink), TCN-like LTR elements (purple) and GC content show as deviation from the genome average (red, above; blue, below). CENP-A and H3K9me2 enriched regions were normalized to input DNA. **(B)** Zoomed-in sections show the regions spanning the centromeres and adjacent genes (light blue). Note that centromeres are enriched for both CENP-A and H3K9me2 marks but are completely devoid of 5mCG DNA methylation despite the presence of this heterochromatic mark in other genomic regions. The data is computed in 5-kb non-overlapping windows.

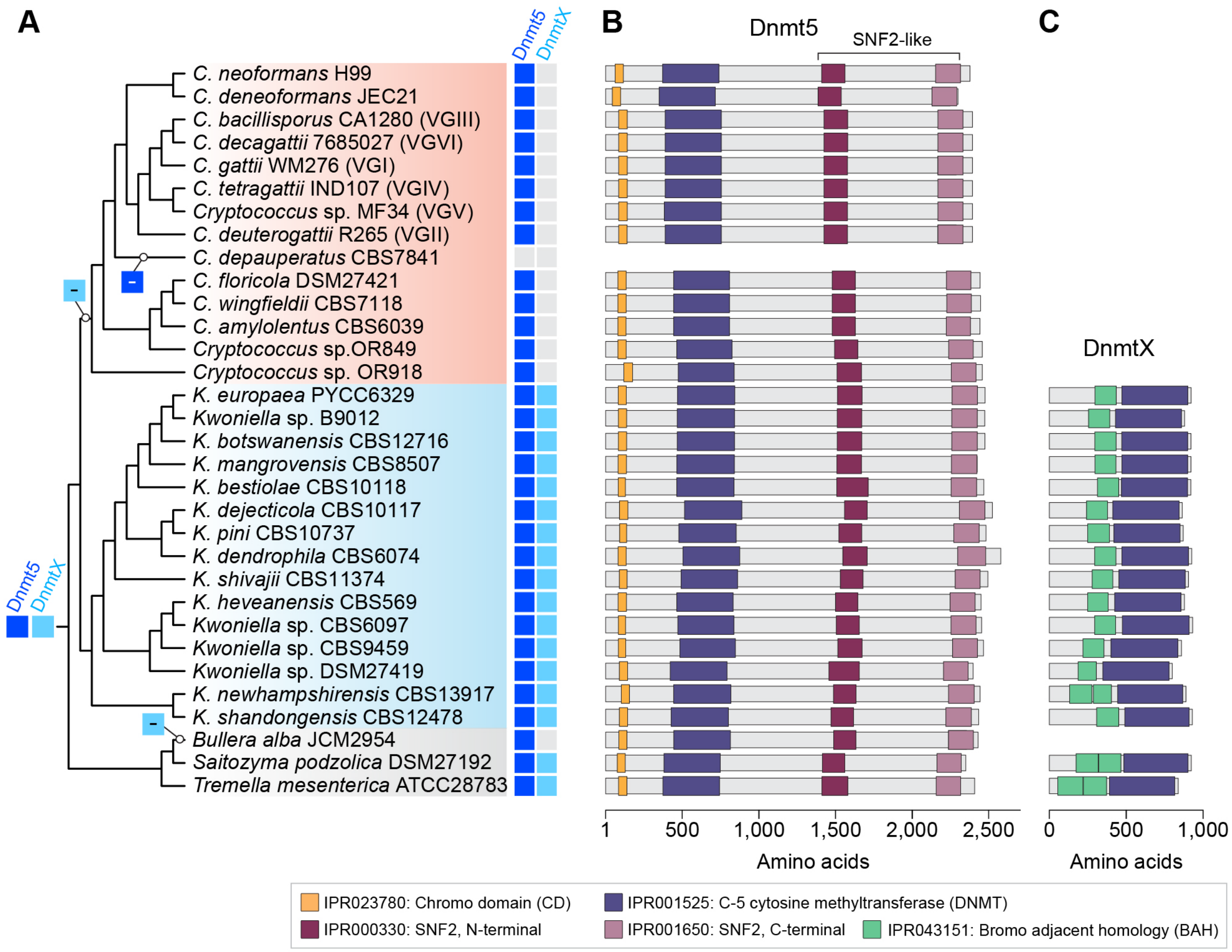


**S11 Fig. Evolution of DNA methyltransferases Dnmt5 and DnmtX in *Cryptococcus* and *Kwoniella* involved in cytosine methylation of DNA.** (A) Species tree topology indicating the presence/absence of two previously characterized DNA methyltransferases: Dnmt5 (encoded by DMT5 gene) is a maintenance-type DNA methyltransferase and DnmtX (encoded by DMTX gene) is a de novo methylase. Phylogenetic and BLAST analysis confirmed initial reports that the ancestral species of the two clades likely had both genes, but *DMTX* was lost in the *Cryptococcus* common ancestor, including in the early-branching *Cryptococcus* sp. OR918 lineage analyzed in this study. The *DMTX* gene is also absent in *Bullera alba* indicating additional losses of are expected to have occurred within the Tremellomycetes. The only instance of *DMT5* loss in our dataset is observed in *C. depauperatus* and our Nanopore data confirms absence of 5mC methylation in this species. **(B)** Similar structure of Dnmt5 across species, characterized by an N-terminal chromodomain (CD) followed by a cytosine methyltransferase catalytic domain (DNMT), and a domain related to those of SNF2-type ATPases **(C)**. The DnmtX protein is shorter and contains a bromo-associated homology (BAH) domain and a DNMT catalytic domain.

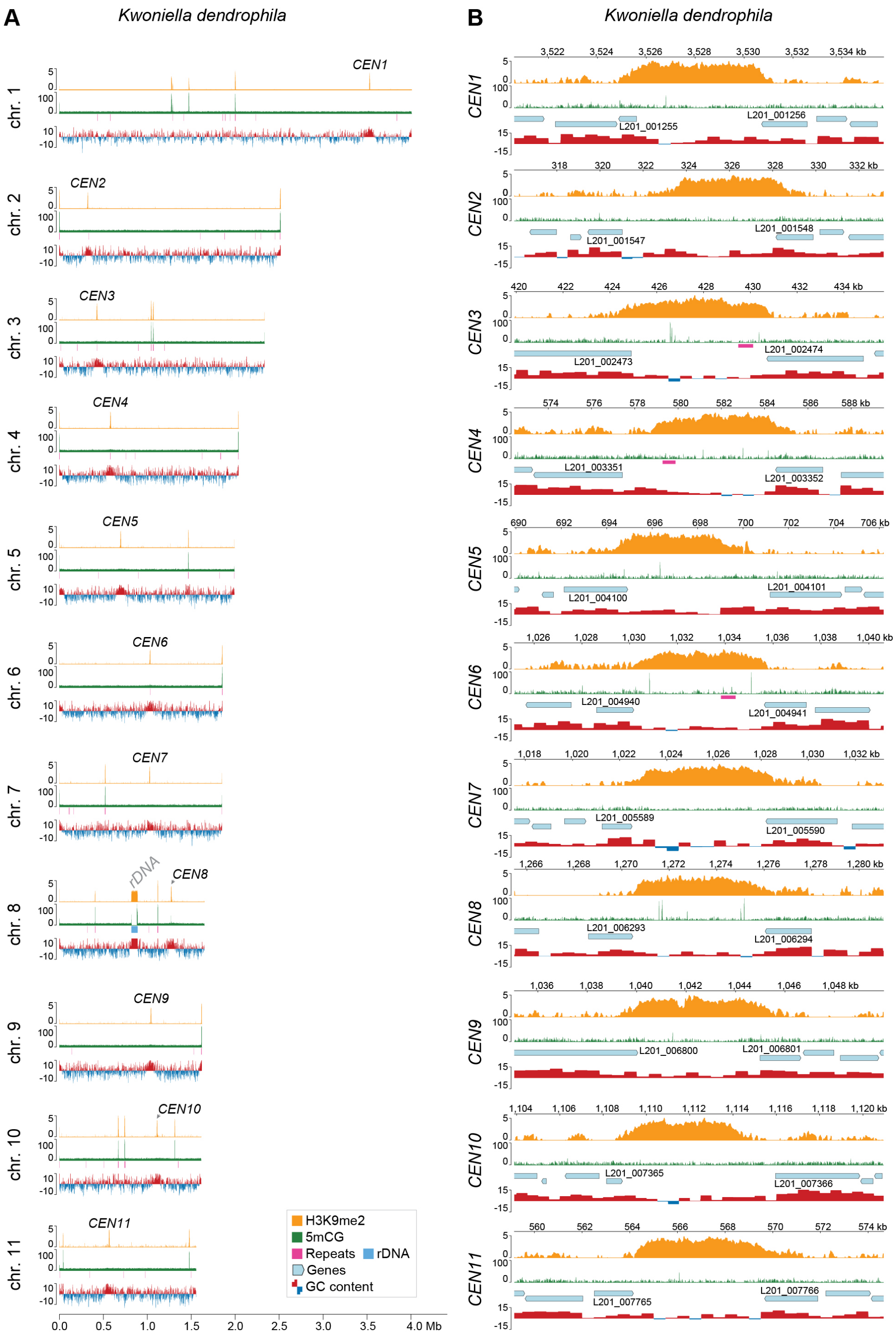


**S12 Fig. The predicted centromeres of *K. dendrophila* lack repeat elements and 5mC DNA methylation but are enriched for H3K9me2*.* (A)** Whole chromosome plots displaying H3K9me2 enrichment (orange), CG cytosine DNA methylation (5mCG, green) from WGBS, repeat content (pink), and GC content show as deviation from the genome average (red, above; blue, below). H3K9me2 enriched regions were normalized to input DNA. **(B)** Close-ups of predicted centromeres and adjacent genes (light blue) highlights H3K9me2 enrichment but absence of 5mCG methylation and transposable elements. The data is computed in 5-kb non-overlapping windows.

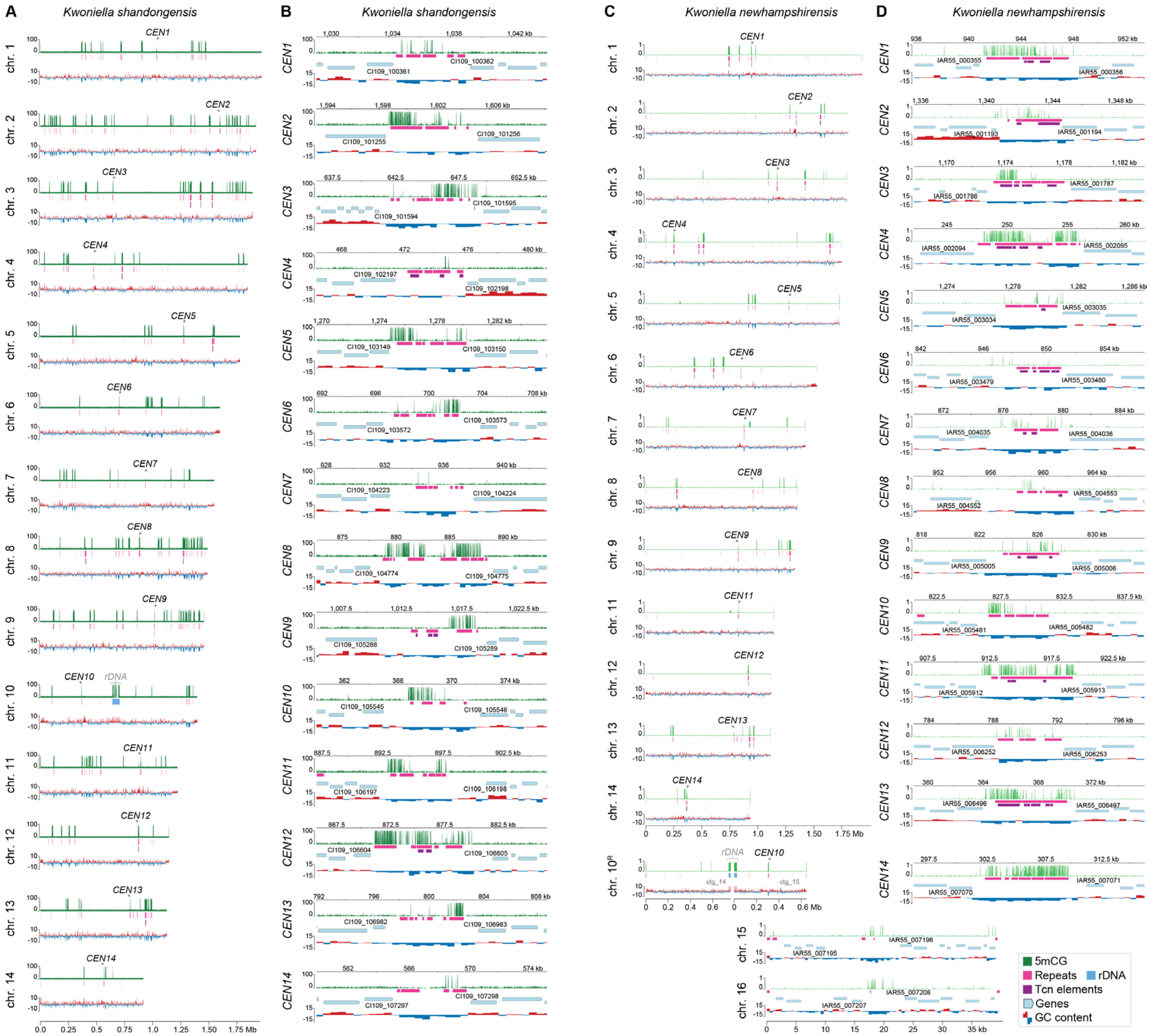


**S13 Fig. Variable presence of LTR retrotransposons within predicted centromeres of *K. shandongensis* and *K. newhampshirensis*.** **(A** and **C)** Whole chromosome plots displaying CG cytosine DNA methylation (5mCG, green) from WGBS (*K. shandogenensis*) or ONT data (*K. newhampshirensis*), alongside repeat content (pink), and GC content show as deviation from the genome average (red, above; blue, below). **(B** and **D)** Close-ups of predicted centromeres and adjacent genes (light blue) highlight diverse 5mCG methylation patterns. While most centromeres encompass unclassified repeats, only a few contain LTR retrotransposons (purple). In *K. newhampshirensis*, chr. 10 corresponds to assembled contigs 14 and 15, broken at the rDNA array, and chrs. 15 and 16 are mini-chromosomes with yet-to-be-determined centromere positions; however, these might correspond to the regions high in 5mCG and low in GC content. The data is computed in 5-kb non-overlapping windows.

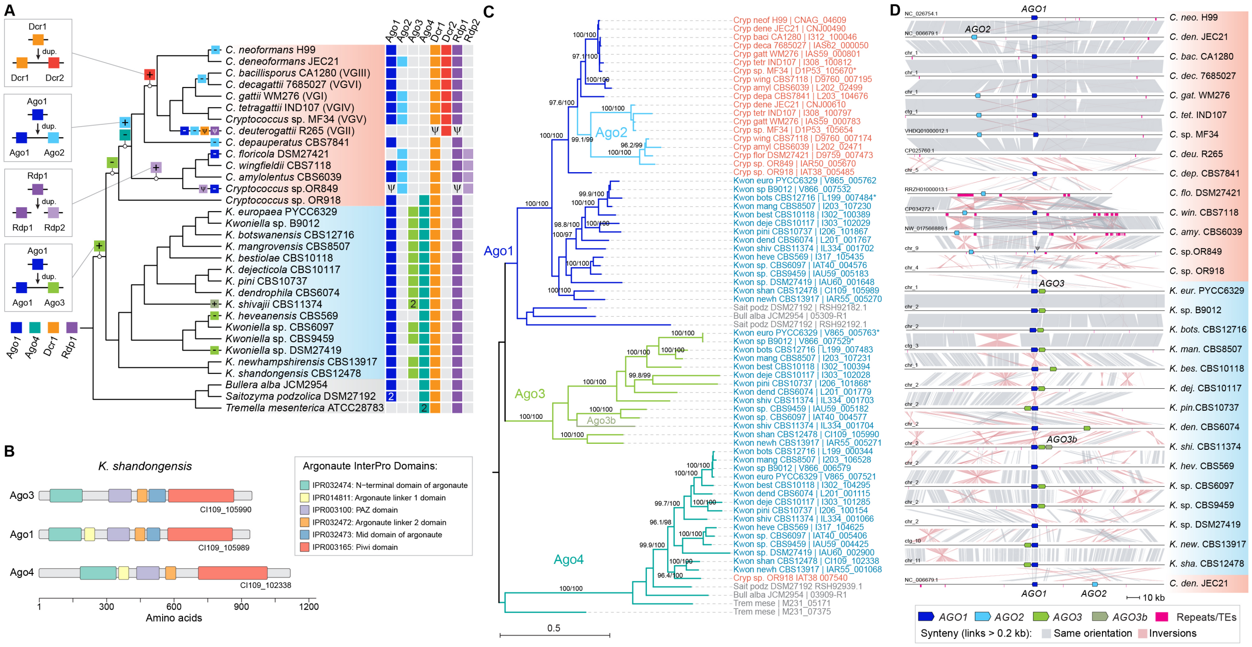


**S14 Fig. Evolution of the core RNAi components in *Cryptococcus* and *Kwoniella*. (A)** Species tree topology indicating the presence/absence of the core RNAi components (Argonaute, Dicer, and RNA-dependent RNA polymerase) across species as well as the inferred pattern of gene gain (via duplication) and loss during evolution. The common ancestor of the two groups was an RNAi-proficient organism, likely expressing two Argonaute proteins (Ago1 and Ago4), one Dicer (Dcr1), and one RNA-dependent RNA polymerase (Rdp1). Psi symbols indicate pseudogenization events and numbers indicate additional species-specific copies. **(B)** Protein domain organization of the three Argonaute proteins found in *K. shandongensis*, depicted here as a representative. **(C)** ML phylogeny of the different Argonaute proteins. The tree was constructed with IQ-TREE2 (model LG+F+R5) and rooted at the midpoint. Internal branch support was assessed by 10,000 replicates of the Shimodaira–Hasegawa approximate likelihood ratio test (SH-aLRT) and ultrafast bootstrap (UFboot). Branch lengths are given in number of substitutions per site. **(D)** Genomic region containing the *AGO1, AGO2* and *AGO3* genes across species. For simplicity, all other genes were omitted. Within the clade comprising *C. floricola*, *C. wingfieldii*, *C. amylolentus*, and *Cryptococcus* sp. OR849, a high prevalence of repeat elements is observed in this region, which may have contributed to the loss of *AGO1* in *C. floricola* and its pseudogenization in *Cryptococcus* sp. OR849.

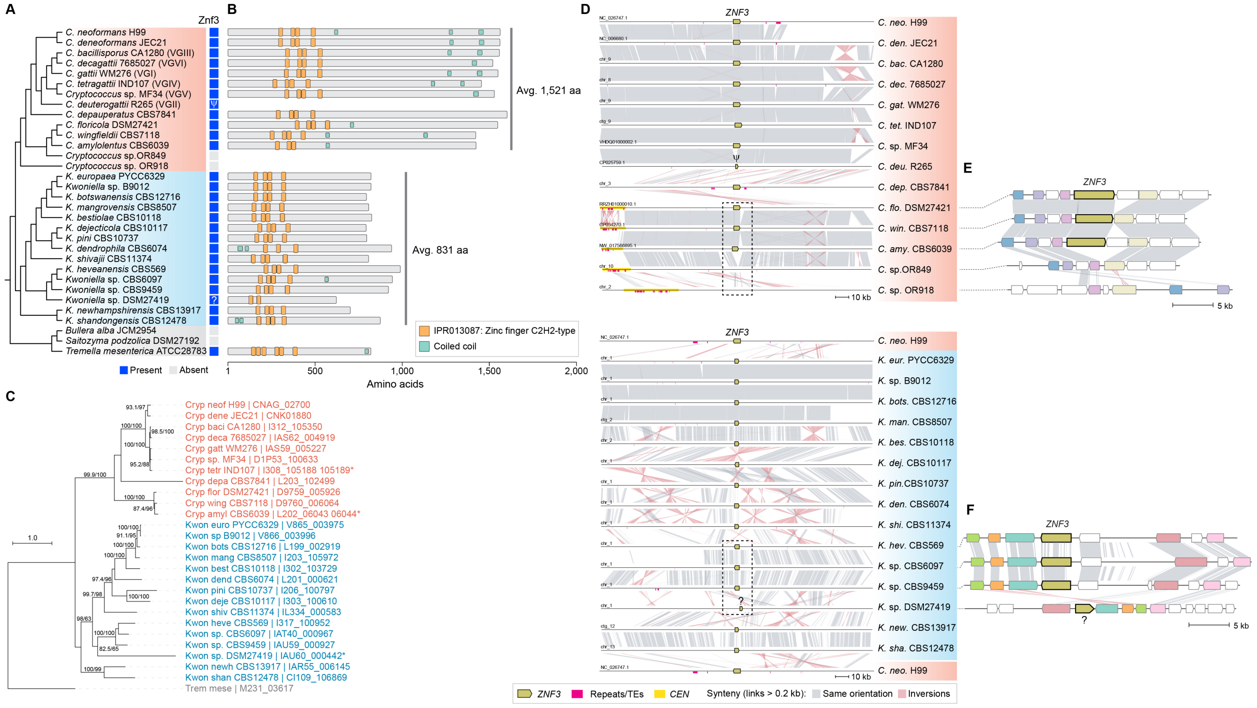


**S15 Fig. Evolution of Znf3 in *Cryptococcus* and *Kwoniella*.** **(A)** Species tree topology indicating the presence/absence of Znf3 across species. **(B)** Protein domain organization of Znf3 proteins. Most of the proteins exhibit four C2H2 zinc finger domains and a few conserved coiled coil regions, often involved in protein-protein interactions. **(C)** ML phylogeny of the different Znf3 proteins. The tree was constructed with IQ-TREE2 (model JTT+F+I+G4) and rooted with *T. mesenterica*. Internal branch support was assessed by 10,000 replicates of the Shimodaira–Hasegawa approximate likelihood ratio test (SH-aLRT) and ultrafast bootstrap (UFboot). Branch lengths are given in number of substitutions per site. **(D)** Genomic region encompassing the *ZNF3* gene across species. For simplicity, all other genes were omitted. The genomic region containing the *ZNF3* in *C. neoformans* H99 was plotted three times for comparison. **(E)** Detailed view depicting newly identified losses of *ZNF3* in *Cryptococcus* sp. OR849 and *Cryptococcus* sp. OR918. **(F)** Detailed view depicting a smaller *ZNF3* gene in *Kwoniella* sp. DSM27419. Colored genes in panels E and F denote those consistently present across all species within the compared region.

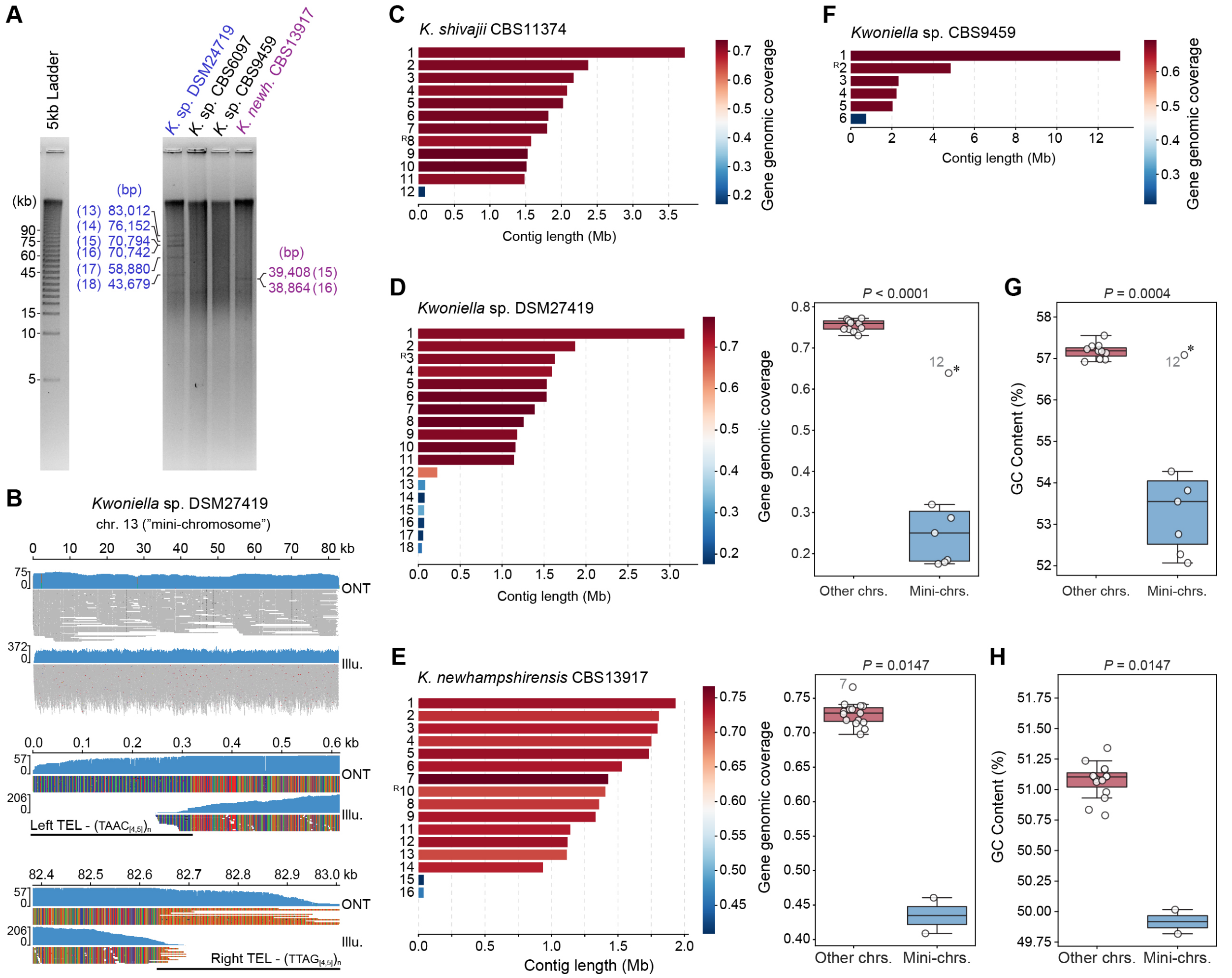


**S16 Fig. *Kwoniella* species with "mini-chromosomes". (A)** Validation of "mini-chromosomes" (less than 100 kb) via PFGE in *Kwoniella* sp. DSM27419 and *K. newhampshirensis* CBS13917, and their absence in *Kwoniella* sp. CBS6097 and *Kwoniella* sp. CBS9459. Contig sizes and respective chromosome numbers are shown next to the gel lane. **(B)** Example of a mini-chromosome assembly validation, evidenced by uniform coverage mapping (blue areas) of both short (Illumina) and long (ONT) reads spanning the full chromosome. The panels below show close-up views of the left and right ends of the mini-chromosome emphasizing the presence of telomeric repeats. **(C-F)** Plots displaying individual chromosome sizes, color-coded based on gene genomic coverage (defined as the ratio of total gene length on a contig to the total length of that contig). In panels D and E, the adjacent boxplots, illustrate statistically significant difference (Mann-Whitney U test) in gene genomic coverage between mini-chromosomes and other chromosomes. **(G-H)** Boxplots showing a notable difference (Mann-Whitney U test) in GC content between mini-chromosomes and other chromosomes in *Kwoniella* sp. DSM27419 and *K. newhampshirensis* CBS13917, respectively. While chr. 12 of *Kwoniella* sp. DSM27419 was categorized as a mini-chromosome in these analyses due to its reduced size (though still larger than 100 kb and thus not visible in the gel in panel A), it exhibits intermediate gene genomic coverage, and a GC content more closely aligned with that of non-mini-chromosomes (refer to **S1 Appendix** for numerical data). Chromosomes containing rDNA are indicated by an R and their size is likely underestimated.

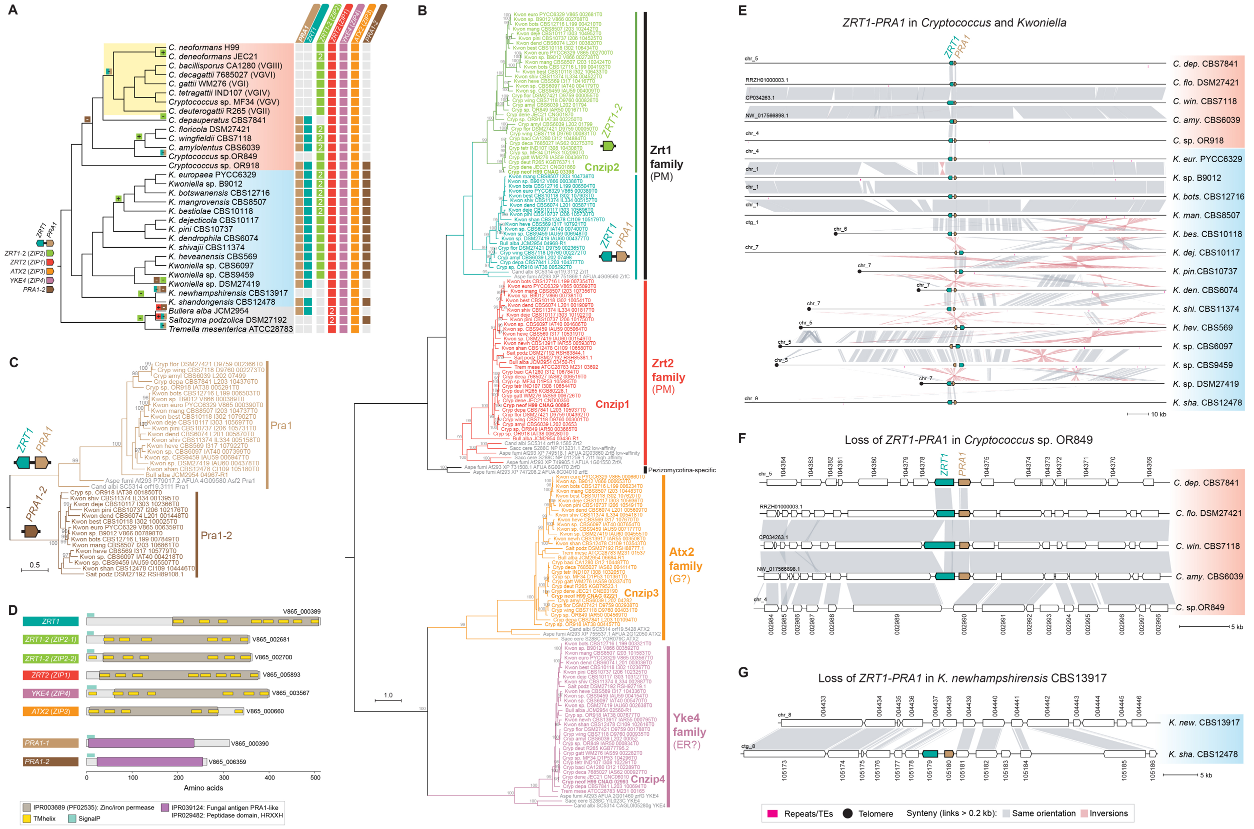


**S17 Fig. Evolution of the *ZRT1-PRA1* pathogenesis gene cluster in *Cryptococcus* and *Kwoniella*. (A)** Species tree topology illustrating the evolutionary trajectory of zinc transporter families and the Pra1 zincophore in *Cryptococcus* and *Kwoniella*. Gene presence in extant species is indicated in color, while absence is shown in grey. **(B)** ML phylogeny of Pra1 with sequences identified across *Cryptococcus* and *Kwoniella*. A diverged copy of *PRA1* (labelled as *PRA1-2*) is present in the early derived *Cryptococcus* sp. OR918 as well as in most *Kwoniella* species. Midpoint rooted phylogenetic trees in panels B and C, with branch lengths representing number of substitutions per site, were constructed with IQ-TREE2 (using models LG+R6 and WAG+G4, respectively), and with internal branch support assessed by 10,000 replicates of Shimodaira–Hasegawa approximate likelihood ratio test (SH-aLRT) and ultrafast bootstrap (UFboot). **(C)** Protein domains of zinc transporters and putative Pra1 zincophores identified in *K. europaea*, shown here as an example. **(D)** ML phylogeny of major zinc-related transporters in our dataset. The tree shows a distinct separation into 5 groups. Orthologs from *Saccharomyces cerevisiae*, *Candida albicans*, and *Aspergillus fumigatus*, previously characterized (show in grey), are included for comparative purposes and to aid in functional prediction. Genes included in the Zrt1 and Zrt2 families are predicted to transport zinc and localize to the plasma membrane (PM); those included in the Atx2 family may have more affinity to manganese transport and localize to Golgi (G); and those within the Yke4 family may function as bidirectional zinc transporters located in the endoplasmic reticulum (ER). The Zrt1 family encompasses two different sets of proteins: Zrt1 and Zip2. The *ZRT1* gene is always clustered with *PRA1*, whereas *ZIP2* is found elsewhere in the genome. Note that the *ZRT1-PRA1* gene cluster was lost in all of the *Cryptococcus* pathogenic species. **(E)** Genomic region encompassing the *ZRT1-PRA1* gene cluster in species where it is present. For simplicity, all other genes were omitted. **(F-G)** Synteny analysis illustrating the species-specific losses of the *ZRT1-PRA* gene cluster in *Cryptococcus* sp. OR849 and *K. newhampshirensis*.

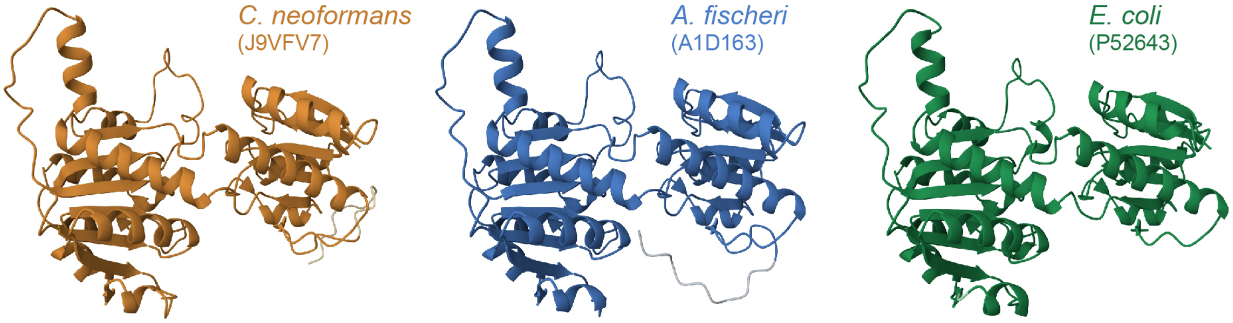


| Species | Uniprot ID | Chain | RMSD | TM-score | Identity | Equivalent Residues | Sequence length | Modelled Residues |
|---|---|---|---|---|---|---|---|---|
| *C. neoformans* | J9VFV7 | A | - | - | - | - | 348 | 348 |
| *A. fischeri* | A1D163 | A | 0.38 | 0.96 | 80% | 336 | 347 | 347 |
| *E. coli* | P52643 | A | 0.72 | 0.93 | 53% | 328 | 329 | 329 |

**S18 Fig. Structural comparison of D-lactate dehydrogenase proteins from AlphaFold predictions.** AlphaFold-predicted structures of *Cryptococcus neoformans* (UniProt J9VFV7), *Aspergillus fischeri* (UniProt A1D163), and *Escherichia coli* (UniProt P52643) are individually displayed at the top. Pairwise structure alignments with the *C. neoformans* protein were conducted with the JjFATCAT-rigid algorithm on the Protein Data Bank website (https://www.rcsb.org/alignment). The table below presents the resulting root mean square deviation (RMSD) and template modeling (TM) scores, among other metrics. The comparison reveals high structural similarity across these proteins, evidenced by low RMSD and high TM scores. The notably lower RMSD score between the two fungal proteins, aligns with the proposed hypothesis of a horizontal gene transfer event from an Aspergilli donor lineage to pathogenic *Cryptococcus* species.

**S1 Text.** Identification and evolutionary analysis of RNAi components in *Cryptococcus* and *Kwoniella*.

**S2 Text**. Identification of shelterin complex and telomere maintenance genes in *Cryptococcus* and *Kwoniella*.

**S1 Appendix**. Genome assembly, genomic features and information on raw sequencing data generated in this study. (A) List of *Cryptococcus* and *Kwoniella* isolates used in this study and summary of genome assembly statistics and other genomic features. (B) Genome sequencing, assembly, and polishing approaches. (C) NCBI accession numbers of each genome and raw read data generated and used in this study. (D) Centromere coordinates and telomeric sequences. (E) Gene genomic coverage and GC content in Kwoniella species containing mini-chromosomes.

**S2 Appendix**. List of genes analyzed in this study with a presumed role in chromosomal integrity. (A) Kinetochore components. (B) RNAi and SCANR complex components. (C) Shelterin and other predicted genes presumably involved in telomere maintenance. (D) List of *S. cerevisiae* essential-DAmP (Decreased Abundance by mRNA Perturbation) genes that exhibit short telomere phenotype and corresponding *Cryptococcus* and *Kwoniella* orthologs. (E) List of *S. cerevisiae* genes that results in shorter telomere length when deleted and corresponding *Cryptococcus* and *Kwoniella* orthologs. (F) DNA and histone methyltransferases

**S3 Appendix.** Significance of branch model fit for *Kwoniella* chromosome-number-based subset, dN/dS results from branch model.

**S4 Appendix**. List of genes absent in *C. depauperatus* and in *Cryptococcus* pathogens, and those that are specifically present in *Cryptococcus* pathogenic species. (A) OGs absent in *C. depauperatus* but present in all other species. (B) OGs absent in *Cryptococcus* pathogens but present in all other species. (C) OGs absent in *Cryptococcus* pathogens but present in 95% of the other species. (D) OGs present in *Cryptococcus* pathogens and absent in all other species.

**S5 Appendix**. Genes involved in capsule and melanin production, growth at 37 ºC, and those that are highly expression in human cerebrospinal fluid (CSF) for *Cryptococcus* pathogens, non-pathogenic *Cryptococcus*, *Kwoniella*, and outgroup species.

**S6 Appendix.** Transposable element content in *Cryptococcus* and *Kwoniella* genomes. (A) EarlGrey results for each TE category and species (related to Fig 5B). (B) Relative percentage of LTR retrotransposons found in centromeric (CEN) versus non-centromeric (Non-CEN) regions (normalized by the total percentage of LTRs; related to Fig 5C). (C) Percentage of TEs versus and average centromere length (related to Fig 5F).

**S7 Appendix.** List of strains, primers and plasmids used in this study.
