## Supplementary material for "Comparative genomics of *Cryptococcus* and *Kwoniella* reveals pathogenesis evolution and contrasting karyotype dynamics via intercentromeric recombination or chromosome fusion": S1 Text

RNA interference (RNAi) is important for genome protection from viral or TE invasion and cellular regulation, involving small-interfering RNA (siRNA) molecules produced by Dicer (Dcr) proteins, loaded onto Argonaute (Ago) proteins, and incorporated into RNA-induced silencing complexes (RISC), resulting in cleavage of target RNA molecules [1]. RNAi in fungi also frequently involves an RNA-dependent RNA polymerase (Rdp), which produces dsRNA [2]. Centromeres are siRNA production sites for transposon inhibition via RNAi in *C. neoformans* and *C. deneoformans*. These species have retained all of the essential RNAi components (Ago, Dcr, and Rdp). In contrast, RNAi-deficient *C. deuterogattii*, lost many of the RNAi components, and has shortened centromeres.

Phylogenetic and synteny analyses across *Cryptococcus* and *Kwoniella* allowed inferring that the common progenitor of the two groups was an RNAi-proficient organism, likely expressing two Argonaute proteins (Ago1 and Ago4), one Dicer (Dcr1), and one RNA-dependent RNA polymerase (Rdp1). (**S14A Fig**). The *AGO1* gene subsequently underwent duplication prior to the divergence of the two groups, giving rise to a third Argonaute copy (*AGO3*) located adjacent to *AGO1* (**S14D Fig**). While *AGO3* was retained in nearly all *Kwoniella* species (except for *K. heveanensis* and *Kwoniella* sp. DSM27419, which independently lost *AGO3* more recently) this copy was apparently lost before diversification of the *Cryptococcus* lineages **(S14A Fig)**. *K. shivajii* has an extra Argonaute gene (*AGO3b*) adjacent to *AGO3*, which is the product of a duplication specific to this species (**S14C and D Figs**). As for *AGO4*, this gene was retained in all *Kwoniella* species and in the early-branching species *Cryptococcus* sp. OR918 but was lost subsequent to the divergence from the shared ancestor with *Cryptococcus* sp. OR918 (**S14A-C Figs**). Remarkably, this loss may have been offset by another duplication of *AGO1*, giving rise to *AGO2*, located ~45 kb away on the same chromosome (**S14D Fig**). Although *AGO1* is conserved among all RNAi-proficient pathogenic *Cryptococcus* species, the distribution of *AGO2* is patchy, punctuated by species-specific losses. This likely reflects previous findings suggesting that this gene seems to play only a minor role in the RNAi pathway in some species [3, 4]. However, among the non-pathogenic *Cryptococcus* species *AGO1*, but not *AGO2*, was lost twice. In *C. floricola*, the loss of *AGO1* may be linked to intrachromosomal rearrangements potentially triggered by repetitive elements, which seem to be abundant in this genomic region (**S14D Fig**). In the case of *Cryptococcus* sp. OR849, a remnant of this gene could still be detected, suggesting a recent pseudogenization event (**S14D Fig**).

Beyond the turbulent evolution of the Argonaute proteins, the Dicer 1 gene (*DCR1*) also underwent duplication specifically within the pathogenic *Cryptococcus* species to produce *DCR2*. Both copies have been retained in all species except *C. deuterogattii*, which lost *DCR1*. Lastly, in the clade encompassing *C. amylolentus* and three related species, *RDP1* was duplicated but then lost in *Cryptococcus* sp. OR849, leaving only a relic of this gene.

Beyond these core components of the RNAi-pathway, we examined eight additional genes required for the global siRNA production in *C. neoformans*: *ZNF3*, *GWC1*, *QIP1*, *RDE1*, *RDE2*, *RDE3*, *RDE4*, and *RDE5* [4-6]. Examination of these genes revealed broad conservation, with *ZNF3* (CNAG_02700) standing out as an exception. *ZNF3* encodes a C2H2-type zinc-finger protein that localizes to P-bodies, and mutations within this gene in *C. neoformans* result in enhanced transposon activity [4]. Additionally, a naturally occurring *znf3* nonsense mutation abolishing siRNA production has been linked to hypermutation in two *C. neoformans* clinical isolates, coinciding with a surge in the accumulation of the non-LTR retrotransposon Cnl1 [7]. Given these findings, phylogenetic and synteny analysis were employed to deepen our understanding of *ZNF3* gene evolution within our focal group of species. We found that *ZNF3* has been lost independently on three separate occasions within *Cryptococcus*: in *C. deuterogattii*, where it became a pseudogene [4], and in *Cryptococcus* sp. OR849 and *Cryptococcus* sp.

OR918, where no traces of the gene were detected (**S15D Fig**). Similarly, *B. alba* and *Saitozyma podzolica*, representing two of the three outgroups, displayed no *ZNF3* counterparts. Interestingly, the predicted protein in *Kwoniella* species and in *Tremella mesenterica* (the third outgroup species) is nearly half the length (avg. 831 amino acids) compared to Znf3 proteins in *Cryptococcus* species (avg. 1,521 amino acids), with the *Kwoniella* sp. DSM27419 Znf3 protein being the smallest (622 amino acids). Despite size differences, most Znf3 proteins exhibit four conserved C2H2 zinc finger domains (**S14B Fig**). It is currently unclear whether all Znf3 proteins retain similar functions or if they have evolved different roles in RNAi in the two groups.

Future studies will address the functional consequences of the RNAi gene duplications and losses and how they might impact the ability to regulate gene expression or defend against transposable elements.
