## Supplementary material for "Comparative genomics of *Cryptococcus* and *Kwoniella* reveals pathogenesis evolution and contrasting karyotype dynamics via intercentromeric recombination or chromosome fusion": S2 Text

Fungal telomeres, like their counterparts in metazoans, are crucial for chromosome stability and integrity. These specialized DNA structures at the ends of chromosomes rely on telomerase for synthesis and a shelterin complex for protection and maintenance of telomere DNA. Telomerase functions by extending short DNA repeats at chromosome ends, while the shelterin complex acts as a guardian, protecting telomeres from degradation and preventing misrecognition of single-stranded telomeric repeats as DNA breaks [1, 2].

In mammals, the shelterin complex comprises six proteins: Trf1 and Trf2, which interact with double-stranded telomere DNA; Pot1, which recognizes the single-stranded G-overhang; and Rap1, Tin2, and Tpp1, which serve as bridging factors [3]. Some of these proteins, such as Tpp1, have roles in recruiting telomerase [4], while Trf1 is thought to recruit the Blm helicase to telomeres, aiding in unwinding G-rich structures for replication fork progression [5]. Fungi exhibit a notable variation in their shelterin components. For example, in *Saccharomyces cerevisiae*, Rap1 identifies telomeric dsDNA with Rif1/2 modulating its length [6], while *Schizosaccharomyces pombe* employs proteins such as Taz1 and Tpz1 without clear orthologous relationship (**S2 Appendix**), illustrating the diversity and evolutionary divergence in fungal shelterin proteins [2, 6]. This is further exemplified in the basidiomycete fungus *Ustilago maydis*, which has unique shelterin proteins such as UmTay1 and UmTrf2, with specific roles in telomere replication and protection, respectively [7].

In light of this diversity, we aimed to identify shelterin complex components in *Cryptococcus* and *Kwoniella* genomes, using sequences from *S. cerevisiae*, *S. pombe*, and *U. maydis* as queries. Our results indicated the presence of Pot1, Tpp1, and a Ten1-like protein in these species. However, other shelterin complex orthologs were not clearly identified, suggesting unique or divergent mechanisms for telomere length regulation and protection.

Complementing our shelterin analysis, we also explored the presence/absence of a set of 152 genes known in *S. cerevisiae* to cause telomere shortening when deleted or when their expression is compromised [8, 9]. While influencing telomere dynamics, these genes also participate in diverse cellular functions, including DNA replication, chromatin remodeling, protein degradation, pre-mRNA splicing, and vesicular transport [9]. This comprehensive analysis showed that an average of 124 out of the 152 genes (~82%) were conserved across the species in our dataset. Importantly, species with "giant" chromosomes, including *K. europaea*, *Kwoniella* sp. B9012, *K. botswanensis*, and *K. mangrovensis*, also exhibited a similar conservation rate (~82.9%) (**S2 Appendix**).

Given the unique and somewhat elusive nature of telomere biology in *Cryptococcus* and *Kwoniella*, as evidenced by our inability to identify clear orthologs for several key shelterin components, future research efforts will be directed towards experimentally elucidating these unique aspects and potentially revealing novel mechanisms of telomere maintenance that might be distinct from those observed in other eukaryotic organisms.
